# Mapping affinity and allostery in human IgG antibody Fc region-Fc γ receptor interactions

**DOI:** 10.1101/2025.03.28.645945

**Authors:** Alasdair D. Keith, Ting Xu, Tatiana A. Chernova, Meredith M. Keen, Marek Bogacz, Marko Nedeljković, Maria Flowers, Trenton Brown, Filipp Frank, Eric A. Ortlund, Eric J. Sundberg

## Abstract

IgG antibodies, required for a functional immune system, recognize antigens and neutralize pathogens using their Fab regions, while signaling to the immune system by binding to host Fc γ receptors (FcγRs) through their Fc regions. These FcγR interactions initiate and modulate antibody-mediated effector functions that are essential for host immunity, therapeutic monoclonal antibody effectiveness and IgG-mediated pathologies. FcγRs include both activating and inhibitory receptors and the relative binding affinities of the IgG Fc region to FcγRs that generate opposing signals is a key determinant of the immune response. Substantial research effort has been devoted to understanding and manipulating FcγR interactions to decipher their fundamental biological activities and to develop therapeutic monoclonal antibodies with tailored effector functions. However, a common Fc-FcγR binding interface, the high sequence identity of FcγRs, and the inherent conformational dynamics of the IgG Fc region, have prohibited a full understanding of these interactions, even when employing state-of-the-art biophysical and biological methods. Here, we used site-saturation libraries of the human IgG1 Fc region to determine the effective affinities of more than 98% of all possible single-site amino acid substitutions in the Fc to all human FcγRs, as well as the most common FcγR polymorphisms. We provide a comprehensive analysis of Fc amino acid variations that determine Fc stability, orthosteric control of FcγR binding, and short- and long-range allosteric control of FcγR binding. We also predict the relative activating versus inhibitory effector function capacity of nearly every possible single-site Fc mutation.

## INTRODUCTION

Immunoglobulin G (IgG) antibodies are critical to host immunity. They function by binding antigenic epitopes using their fragment antigen-binding (Fab) regions and signaling to other cells in the immune system through interactions with host receptors by their fragment crystallizable (Fc) regions. The neutralization capacities of IgG antibodies derive from Fab engagement with the infectious agent, while antibody-mediated effector functions rely on Fc region interactions with Fc γ receptors (FcγRs) and complement C1q. Beyond their importance in host immunity, Fc-FcγR interactions are often required for the effectiveness of therapeutic monoclonal antibodies (mAbs),^1,2^ such as in cancer treatment,^3,4^ and act as drivers of many IgG-mediated pathologies, including numerous autoimmune diseases.^5,6^

FcγRs act as regulators of immune responses, as they are expressed on the surfaces of effector leukocytes and include activating and inhibitory family members containing immunoreceptor tyrosine-based activation and inhibitory motifs (ITAMs and ITIMs, respectively).^7^ In humans, the FcγR family consists of one high-affinity receptor, FcγR1, and several low-affinity receptors, including FcγR2a, FcγR2b and FcγR3a,^8^ all of which are activating receptors with the exception of FcγR2b, which is inhibitory. The relative binding affinities of the Fc region to these low-affinity activating and inhibitory FcγRs dictates effector functions.^9^ For instance, differential binding to IgG Fc regions by FcγR2a, FcγR2b and FcγR3a influences antibody-dependent cellular cytotoxicity (ADCC) and antibody-dependent cellular phagocytosis (ADCP).^10^ FcγR2a, typically present on monocytes, neutrophils, macrophages, dendritic cells and platelets,^8,11^ induces ADCP to promote phagocytosis. FcγR3a, present on natural killer (NK) cells, neutrophils, monocytes and macrophages,^11^ is a key contributor to targeted cell death *via* ADCC.^12^ FcγR2b is highly expressed on B cells, where it is the only FcγR present, in addition to being expressed at lower levels on monocytes, macrophages, and dendritic cells,^8^ and on neutrophils and NK cells of individuals with certain genotypes.^13–15^ This cellular distribution allows FcγR2b to interfere effectively with the activity of other activating FcγRs, negatively regulating both ADCP^7,16^ and ADCC.^17,18^

An additional complicating factor in antibody-mediated effector functions is that both FcγR2a and FcγR3a have common polymorphisms in humans. For FcγR2a, a genotypic variation at position 131, encoded as either a histidine (H) or an arginine (R) residue, results in distinct clinical outcomes: patients with the homozygous H/H isoform benefit most from treatment with the therapeutic mAbs rituximab^19^ and trastuzumab^20^ compared to those with the H/R or R/R isoforms, due to a higher binding affinity to the Fc regions of these mAbs. Similarly, the polymorphisms of FcγR3a, encoded as either a phenylalanine (F) or valine (V) residue at position 158, have different binding affinities to the Fc domain; the FcγR3a-158V variant binds Fc regions with higher affinity than does FcγR3a-158F, resulting in increased binding by immune cells,^21,22^ which in turn enhances responses to monoclonal antibody therapy.^23^

Manipulating the relative binding affinities of inhibitory (I) versus activating (A) FcγRs – expressed here as the I:A ratio – is a key engineering strategy for tuning ADCP and ADCC responses of therapeutic mAbs. Increasing ADCC is of critical importance to many cancer-based therapeutics,^24–26^ and can be achieved by engineering the Fc region to exhibit increased binding to FcγR3a, decreased binding to FcγR2b, or a combination of the two. ADCP constitutes a more subtle immune response,^27^ but is also of significant consequence in cancer treatment^3,4^ and other immunotherapies;^1,2^ this response can be enhanced by engineering Fc regions to bind with higher affinity to FcγR2a and/or lower affinity to FcγR2b. Meanwhile, enhancement of binding to FcγR2b has been demonstrated to mitigate certain types of autoimmunity^5,28^ and preclinical candidates have been developed,^29,30^ but identification of suitable Fc mutations is difficult given that its extracellular domain has especially close sequence identity and structural similarity to that of FcγR2a.^31^

The structure of the IgG Fc region is conformationally dynamic, adopting both “open” and “closed” conformations due to the presence and absence, respectively, of a conserved *N*-linked glycan at residue Asn297; although, the entire ensemble of physiologically-relevant Fc conformations is likely much more extensive.^32^ This conformational flexibility of the Fc region results in allosteric effects on Fc-FcγR interactions that have not been fully elucidated despite the many Fc mutations interrogated for manipulating activating versus inhibitory FcγR binding affinities. No systematic, comprehensive evaluation of IgG Fc variants for FcγR binding, though, has yet been performed. Here, we employed deep mutational scanning (DMS) analysis using a mammalian surface-display platform to measure the effect of more than 98% of all possible single-site mutants of the human IgG1 Fc region on binding to each of the human FcγRs and their common polymorphisms. Since our site-saturation mutagenesis approach was untargeted, we were able to identify myriad IgG Fc sites that affected FcγR interactions that were previously overlooked by mutagenesis experiments designed based on structural analyses. This approach has allowed us to comprehensively map affinity and allostery in human Fc-FcγR interactions.

## RESULTS

### Establishing a mammalian cell surface display platform for the IgG1 Fc region

We adapted our previously reported lentiviral expression system^33^ to contain the hinge and Fc regions (inclusive of residues 216-447) of human IgG1. In this construct, IgG1_216-447_ is framed by an N-terminal signal peptide (SP) derived from IgG4 and a C-terminal transmembrane (TM) region from the platelet-derived growth factor receptor (PDGFR), with a Myc-tag inserted between the SP and IgG1_216-447_ to monitor expression variation.^34^ We cloned this construct into a lentiviral expression plasmid containing a 3’ internal ribosomal entry site (IRES) and green fluorescent protein (GFP), which acts as a selection marker (**Figure 1A**). We produced a stable cell line expressing IgG1_216-447_ on the surface of mammalian HEK293T cells and assessed human FcγR binding to validate the system. We incubated cells with titrated concentrations of individual FcγRs and subsequently stained the cells with fluorescently-labeled streptavidin and anti-Myc antibodies to quantify FcγR binding and IgG1_216-447_ cell surface expression, respectively (**Figure 1B**). We analyzed cells by flow cytometry to determine dissociation constants for all human FcγRs and common polymorphisms, including FcγR1, FcγR2a-131H, FcγR2a-131R, FcγR2b, FcγR3a-158F and FcγR3a-158V, by gating on GFP^+^/Myc^+^ cells and quantifying normalized median fluorescence intensity (MFI) of the streptavidin signal (i.e., FcγR; **Figure 1C**). The *K*_D_ values that we measured in this way for surface-displayed IgG1 Fc-FcγR interactions were consistent with published data derived by surface plasmon resonance (SPR) analysis of the same Fc-FcγR interactions^35^ (**Figure 1D**). IgG Fc regions are obligate homodimers and FcγRs bind only to homodimeric Fc regions, indicating that when expressed on the HEK293T cell surface as described, individual IgG1_216-447_ Fc protomers pair as homodimers as depicted in **Figure 1B** to become competent for FcγR binding.

**Figure 1.**
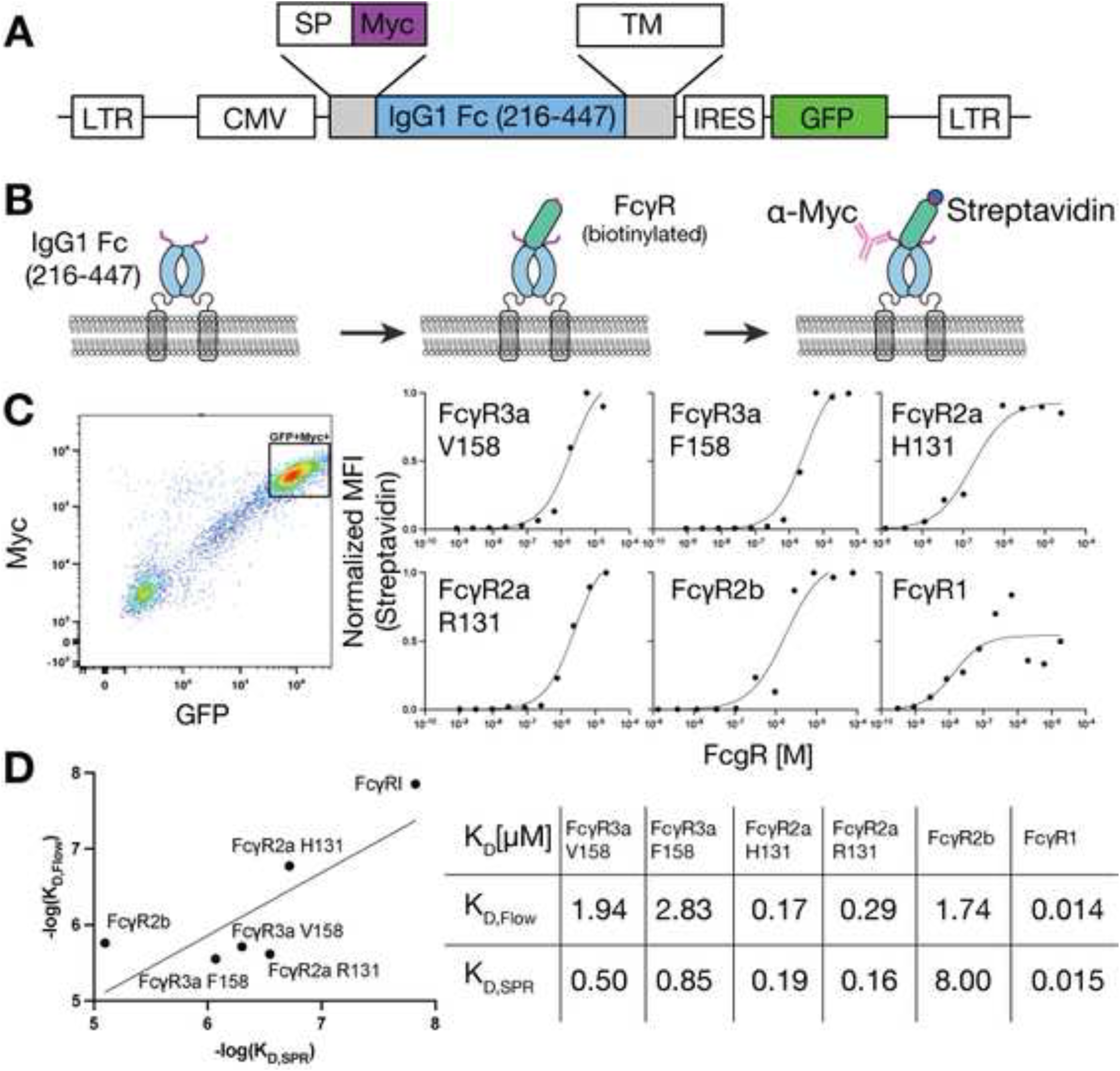
IgG1 Fc cell surface display system. **(A)** Construct design for the surface expression of IgG1 Fc on HEK293T cells. Additional elements required for deep mutational scanning include a cytomegalovirus (CMV) promoter, a signal peptide (SP) for translocation to the cell membrane, a transmembrane (TM) region and tags. Following cloning into a lentiviral expression plasmid, co-expression of the target protein and a GFP marker was achieved from the same mRNA via an internal ribosome entry site (IRES). **(B)** Schematic for the detection of FcγR binding to cell surface-displayed IgG1 Fc. **(C) *Left*,** Flow cytometry analysis of cells stably expressing IgG1 Fc on the cell surface. The majority of cells are GFP^+^/Myc^+^. ***Right*,** Titration curves for all FcγRs binding to cell surface-displayed IgG1 Fc. GFP^+^/Myc^+^-gated cells were analyzed for biotinylated FcγR binding signal via streptavidin normalized median fluorescence intensity (MFI) measurements. **(D)** Validation of dissociation constants (*KD*s) determined from cell surface binding titrations, as in (C), between WT IgG1 Fc and all FcγRs with those published in the literature^35^ using soluble recombinant protein and surface plasmon resonance (SPR) analysis.

### Cell surface display of an IgG1 Fc deep mutational scanning library

Our strategy for implementing an IgG1 Fc deep mutational scanning library is depicted in **Figure 2A**. We designed a site-saturation library encompassing all possible amino acid substitutions of IgG1_216-447_ and retaining the flanking regions from wild type surface display, as shown in **Figure 1A**. We then used PCR amplification to insert 15-nucleotide randomized barcodes immediately downstream of the IgG1_216-447_ stop codon. Following cloning of the library into the previously described lentiviral expression vector, we used PacBio long-read sequencing to produce a lookup table bracketing unique barcodes with distinct amino acid mutants. Our mutational coverage was 4,359 out of 4,408 (98.9%) total possible single amino acid substitutions, with only two positions (242 and 431) predominantly missing mutations, due to the input library failing to produce mutants (**Figure 2B-C**). Each variant was coupled to nine distinct barcodes, on average, and the vast majority of variants were single-site substitutions, as intended by the library design (**Figure 2B**). However, substantial numbers of variants with zero (wild type IgG1_216-447_) and two mutations (combinatorial mutants) also exist in our IgG1_216-447_ library. For subsequent experiments, we packaged the above-described library into lentiviral particles and then transduced HEK293T cells with a multiplicity of infection (MOI) close to 0.1, ensuring that only a single IgG1_216-447_ mutant is expressed by the majority of individual cells. We used fluorescence-activated cell sorting (FACS) in a two-step verification process to determine the success of transduction (i.e., GFP^+^ cells) and expression of the variant on the cell surface (i.e., Myc^+^ cells).

**Figure 2.**
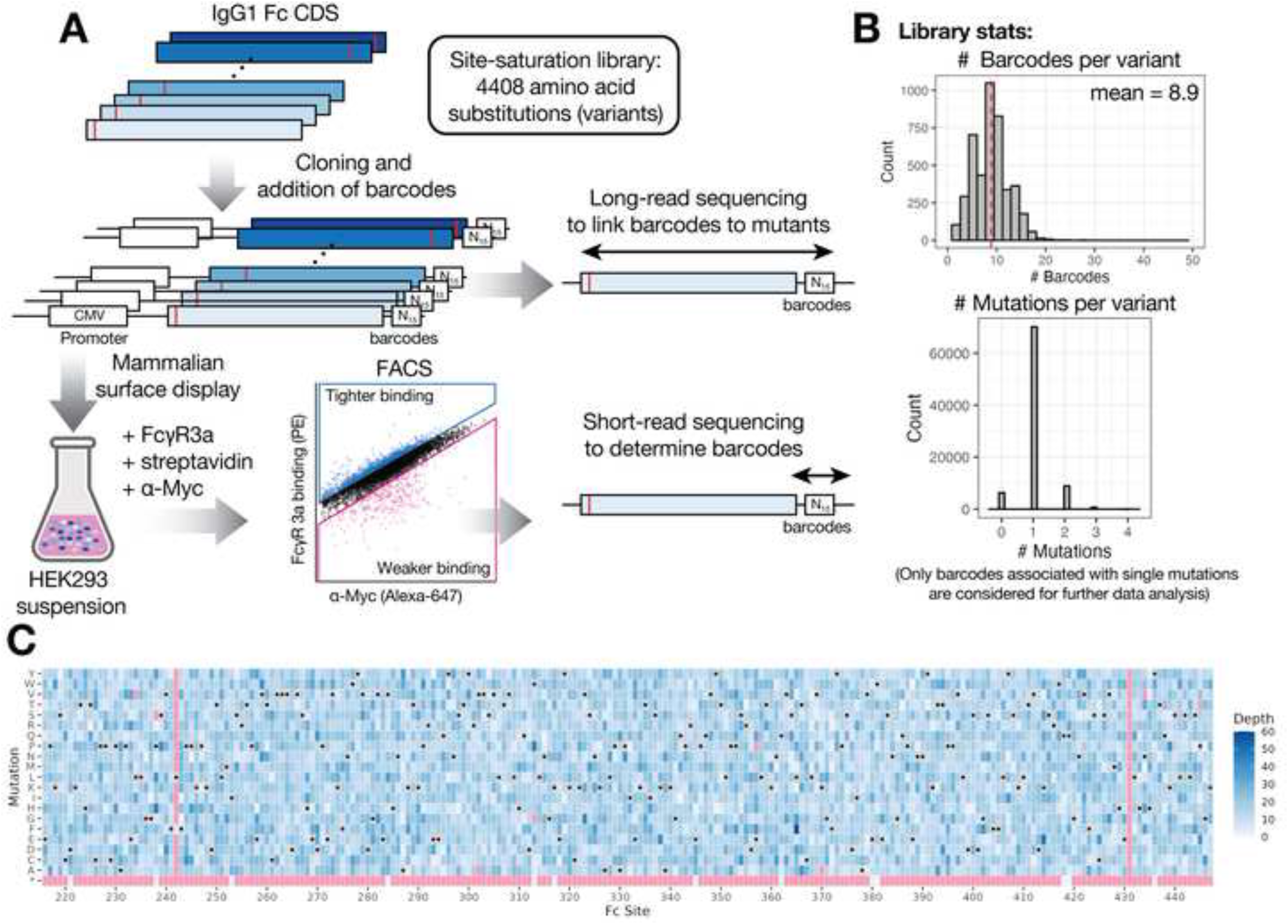
IgG1 Fc deep mutational scanning workflow. **(A)** Schematic of the deep mutational scanning approach for measuring FcγR binding to a site-saturation mutant library of IgG1 Fc. **(B)** PacBio long-read sequencing library statistics. Each mutation is represented by multiple, independent barcodes (***top***), and the majority of variants in the library contain a single-point mutation (***bottom***). **(C)** PacBio long-read sequencing read depth. Library coverage was 4,359 out of 4,408 (98.9%) total possible amino acid substitutions, with only two sites (242 and 431) largely failing.

### IgG1 Fc mutations that inhibit cell surface display are predominantly localized to the core of the CH2 domain

To identify single-site IgG1 Fc variants that negatively impact cell surface expression, likely due to their effects on protein stability and/or trafficking to the plasma membrane, we sorted for GFP^+^/Myc^-^ populations (**Figure 3A**) followed by barcode sequencing. A heatmap highlighting the residue and site-level effects on cell surface expression (**Figure 3B**) reveals a discontinuous pattern of enrichment located predominantly in solvent inaccessible sites within the N-terminal CH2 domain, inclusive of the disulfide bond between residues Cys261 and Cys321 (**Figure 3C-D**). Most sites were distal to the Myc tag, which is consistent with the hypothesis that these mutations have a significant role in altering protein stability, rather than in any direct interference with the Myc tag recognition. Not only were the number of positions that negatively impacted Fc cell surface display in the CH2 domain much greater than in the CH3 domain – thirteen versus two – each of these positions in CH2 was also much less tolerant to substitution by different amino acids than those in CH3 (**Figure 3B**). For most of these positions in the CH2 domain, substitution by nearly any amino acid other than the wild type residue had a sizable negative effect on surface expression. We found few residues in the CH3 domain that affect Fc surface display, with the exception of residues Cys367 and Cys425, which form a disulfide bond, and residues Leu365 and Val367 that are adjacent to this disulfide bond (**Figure 3E**). Although these residues are central to the hydrophobic core of the CH3 domain, they were nonetheless tolerant to substitution by a wide range of amino acids; substitutions not tolerated at these positions were almost exclusively charged amino acids (**Figure 3B**). Increased sensitivity to mutation of the CH2 domain is consistent with its lower thermal stability compared to the CH3 domain.^36,37^ The CH2 domain generally exhibits the lowest melting point of the cooperative folding units, i.e. the individual Ig domains, in a mAb and increasing its stability is of great value in the design of therapeutic mAbs with improved biophysical and developability properties.

**Figure 3.**
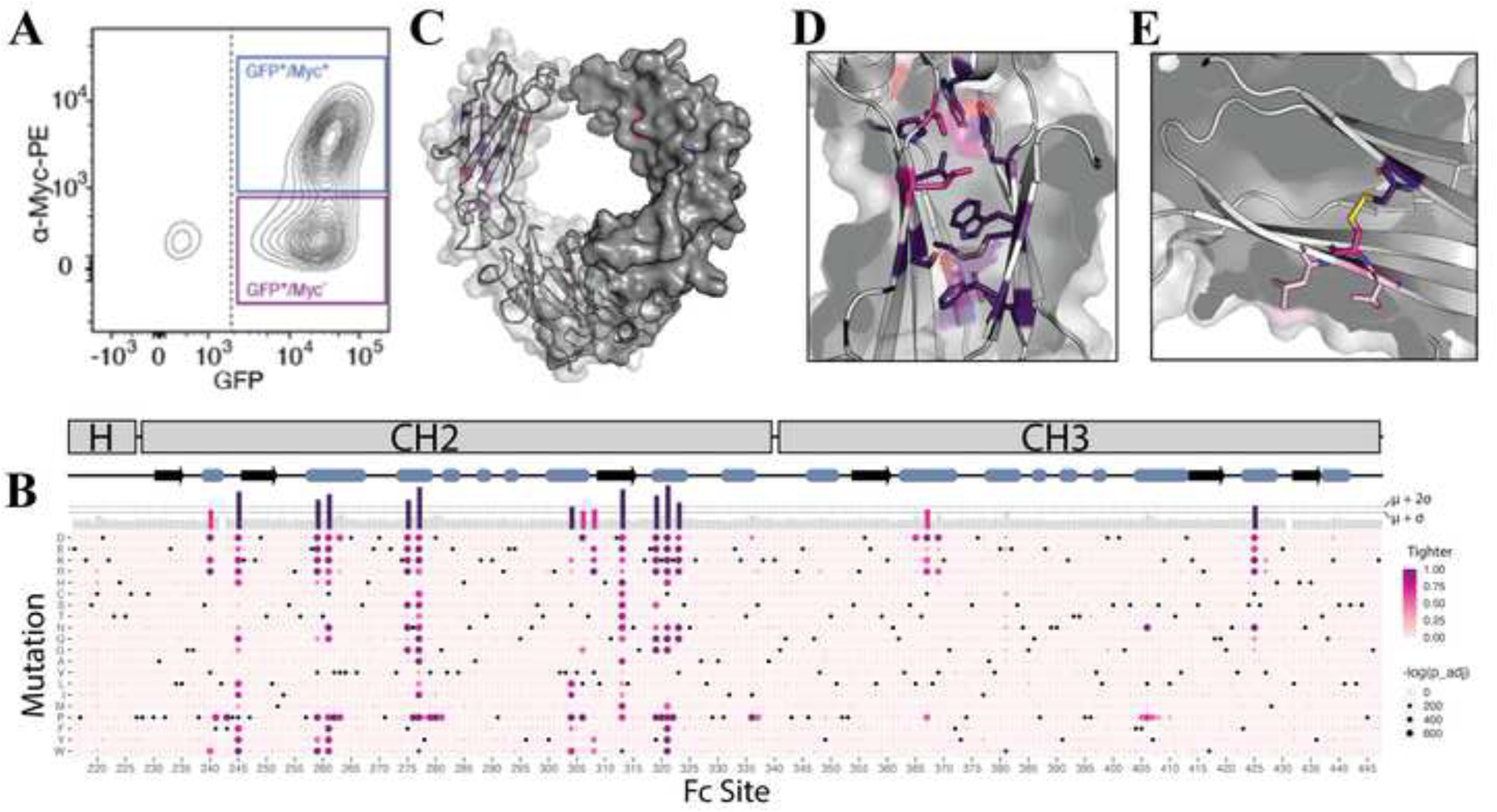
IgG1 Fc mutations that disrupt cell surface expression. **(A)** Fluorescence-activated cell sorting (FACS) strategy. The GFP^+^/Myc^-^ population was selected for this analysis as it gives a measure of protein instability and/or inability of trafficking for cell surface expression. The GFP^+^/Myc^+^ population is typically chosen for measurements of Fc-FcγR binding as it denotes IgG1 Fc variants that are functionally folded and expressed on the cell surface. **(B)** Heatmap of deep mutational scanning results from the IgG1 Fc GFP^+^/Myc^-^ population. The Fc sequence is charted along the x-axis, and mutations along the y-axis. The wild type sequence is represented by a series of black dots. Individual mutations are described by colored dots, which scale in shading according to enrichment vs a reference population, and in size according to their associated p-values. The bar graph situated above the heatmap shows total site enrichment scores (i.e., the sum of all 19 scores at each Fc position). **(C)** Structural representation of the Fc dimer, with individual residues highlighted if the total enrichment score exceeds one (pink) or two (magenta) standard deviations, s, above the mean, m. **(D)** Magnification of the Fc CH2 domain core region, revealing that the most destabilizing mutations are typically transformations of hydrophobic residues in the protein core. **(E)** Magnification of the Fc CH3 domain core region, revealing that it is mainly tolerant to destabilizing mutations.

### Mapping IgG1 Fc mutations that lead to tighter and weaker FcγR binding

As for the cell surface display analysis described above, we expressed our IgG1 Fc deep mutation scanning library in HEK293T cells, but this time sorted for GFP^+^/Myc^+^ cells, incubated these cells with biotinylated FcγRs at concentrations equivalent to the *Kd* of the individual Fc-FcγR interactions as determined by titration analysis (**Figure 1C**), and stained with streptavidin-PE. Using FACS, we selected cells along the FcγR-binding/α-Myc diagonal to isolate tighter and weaker Fc-FcγR binding populations corrected for mammalian surface expression. We deep-sequenced these two cell populations, along with an unsorted reference library, identified and counted barcodes for each population, and linked these data to the amino acid variants that they represent. Using **Equation (1)** we determined the ratios of these sorted populations versus the reference library to generate scores for tighter and weaker binding for all IgG1 Fc amino acid variants in our library binding to each of the six FcγRs. Results for IgG1 Fc variants binding to FcγR3a-158V are shown in **Figure 4A-B**. As expected, all mutations of Asn297, onto which an *N*-glycan critical for FcγR binding is linked,^38^ abrogated FcγR3a-158V binding, as indicated by the high weaker binding scores for all non-asparagine amino acids at this position. We mapped the cumulative tighter and weaker binding scores at each IgG1 Fc position to the Fc-FcγR3a crystal structure (PDB: 3SGJ)^39^ (**Figure 4C**) in order to visualize the location of mutations that had substantial effects on FcγR3a-158V binding. While we could clearly identify mutations in the IgG1 Fc residues that form the interface with FcγR3a-158V due to their high scores for weaker binding, we found numerous residues distal to the Fc-FcγR3a-158V interface that exhibited both significant tighter and weaker binding scores, indicating a range of allosteric effects that can lead to both increased and decreased Fc-FcγR3a-158V binding affinity. We performed the FcγR3a-158V experiments in replicate using two independent replicate cell lines with completely independent sets of barcodes and found that our results were highly reproducible for both tighter (**Figure 4D)** and weaker (**Figure 4E**) binders. By analogous methods, we evaluated IgG1 Fc variants for binding to FcγR1, FcγR2a-131H, FcγR2a-131R, FcγR2b and FcγR3a-158F. We observed the same general pattern of mutations in, and distal to, the Fc-FcγR interface leading to tighter and weaker binding for all FcγRs, as shown in the heatmaps in **Figure S1**, as well as mapped to the structures of the respective Fc-FcγR complexes (**Figure 4F-J**).

**Figure 4.**
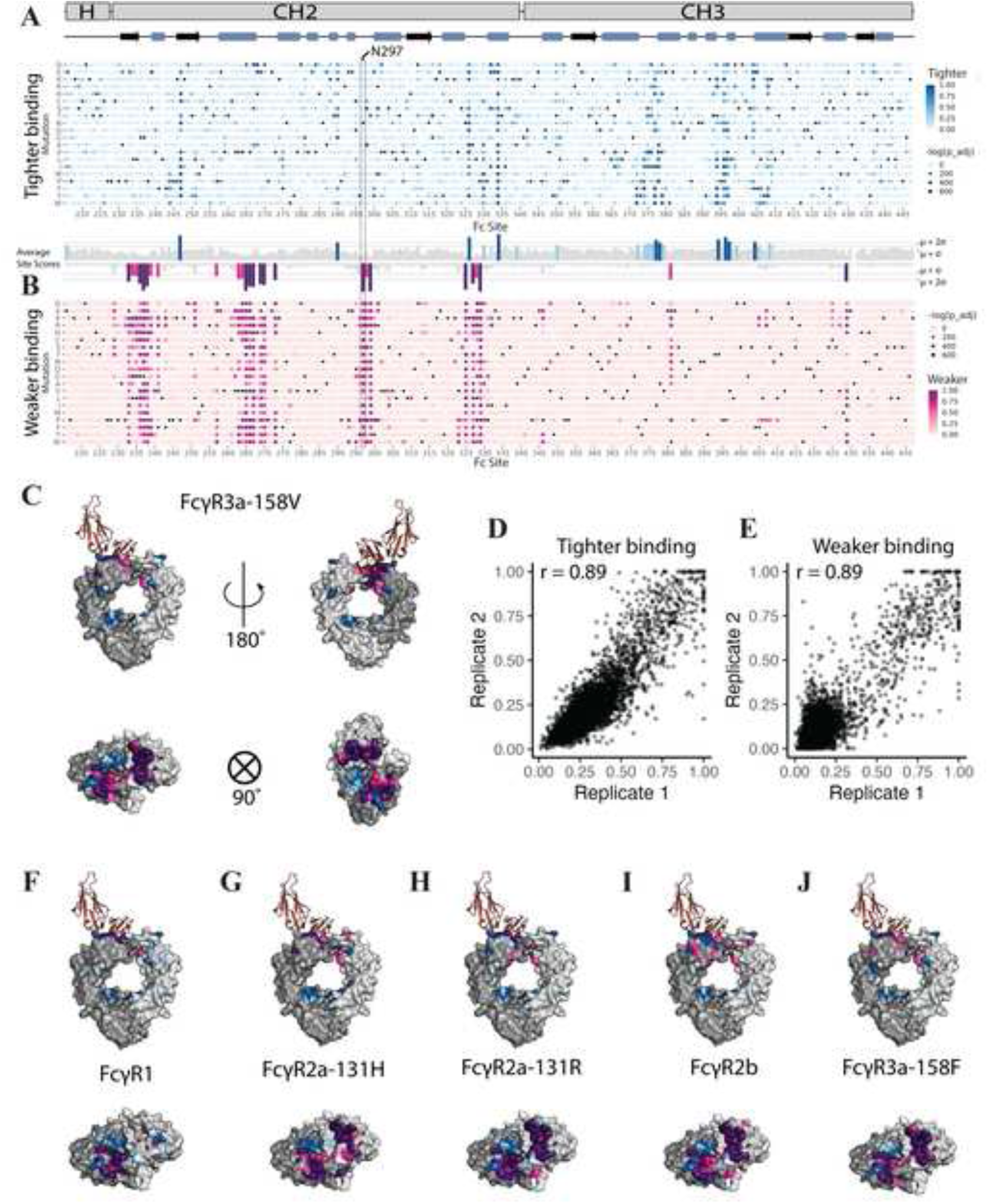
IgG1 Fc deep mutational scanning at a single concentration. Heatmaps of deep mutational scanning results for IgG1 Fc mutants with **(A)** tighter and **(B)** weaker binding to FcγR3a-158V. Average tighter/weaker binding scores per site are depicted as bar charts between the heatmaps, with sites highlighted if the score exceeds one (light blue/pink) or two (dark blue/magenta) standard deviations above the mean. N297 is boxed, highlighting reduced binding upon loss of glycosylation. **(C)** Average tighter (dark/light blue) and weaker (magenta/pink) binding scores, color-coded according to the threshold values described in (B), mapped onto Fc dimer structures. FcγR3a-158V is depicted in orange. Correlations between two replicates of IgG1 Fc DMS analyses with FcγR3a-158V with **(D)** tighter and **(E)** weaker binding populations. Average tighter and weaker binding scores mapped onto Fc dimer structures for **(F)** FcγR1, **(G)** FcγR2a-131H, **(H)** FcγR2a-131R, **(I)** FcγR2b, and **(J)** FcγR3a-158F.

### Determining the effective affinities of single-site amino acid substitutions in the IgG1 Fc to all low-affinity FcγRs

Although our deep mutational scanning strategy described above provides a comprehensive analysis of the effects of nearly all possible Fc mutations on binding to individual FcγRs, the tighter and weaker binding scores are not directly comparable to biophysical analyses of recombinant proteins (such as biolayer interferometry [BLI] or surface plasmon resonance [SPR] analyses), nor do they provide a convenient means for comparing scores between assays. Since antibody-mediated effector functions are dependent on the relative binding of activating (FcγR2a, FcγR3a) versus inhibitory (FcγR2b) low-affinity FcγRs, we sought to determine effective affinities that would be directly comparable for all possible Fc mutations to all of the low-affinity FcγRs. In order to do so, we tested Fc variant binding against ten concentrations for each FcγR, spanning more than two orders of magnitude both above and below the *K*_D_ values for each wild type interaction (**Figure 5A**). Following the methodology introduced by Starr *et al.*,^34^ we employed a partitioning strategy by separating cells into four indexed bins, from lowest binding to highest binding, for each concentration, resulting in 40 distinct populations of cells per experiment (**Figure 5B**). After sequencing the barcodes from all populations and grouping barcodes to their relevant variants, we used the distribution of reads across the bins to calculate a mean bin for each variant at each concentration. The resulting titration curves allowed us to calculate effective dissociation constants, *K*_D_s, for each single-site variant (i.e., a ‘per variant binding phenotype’), as shown for one library member for which the barcode corresponds to wild type Fc (**Figure 5C**); the *K_D_* value for this interaction that we derived by this method is nearly identical to that determined independently by SPR.^35^ We then plotted these data in heatmaps, color-coded according to log*K*_A_ values (where *K*_A_ is an effective association constant, with log*K*_A_ = -log*K*_D_), as shown for the Fc–FcγR3a-158V interaction (**Figure 5D**). We normalized these heatmaps such that the log*K*_A_ value of each wild type Fc residue is set to white in the color-coded spectrum of log*K*_A_ values. We used association constants to avoid negative signs and, therefore, simplify the interpretation of differential heatmaps as described below. For the Fc–FcγR3a-158V interaction, we compared the weaker minus tighter binding scores and titration log*K*_A_ values for all Fc mutants analyzed and found them to be highly correlated (**Figure 5E**). We averaged these log*K*_A_ values to calculate per-site values and mapped all of the effective affinities for the Fc–FcγR3a-158V interaction to the structure of this complex (**Figure 5F**). Using analogous methods, we generated effective affinities for all Fc mutants binding to each of the remaining low-affinity FcγRs – FcγR2a-131H, FcγR2a-131R, FcγR2b and FcγR3a-158F – which we plotted in heatmaps (**Figure S2**) and mapped to Fc-FcγR complex structures (**Figure 5G-K**).

**Figure 5.**
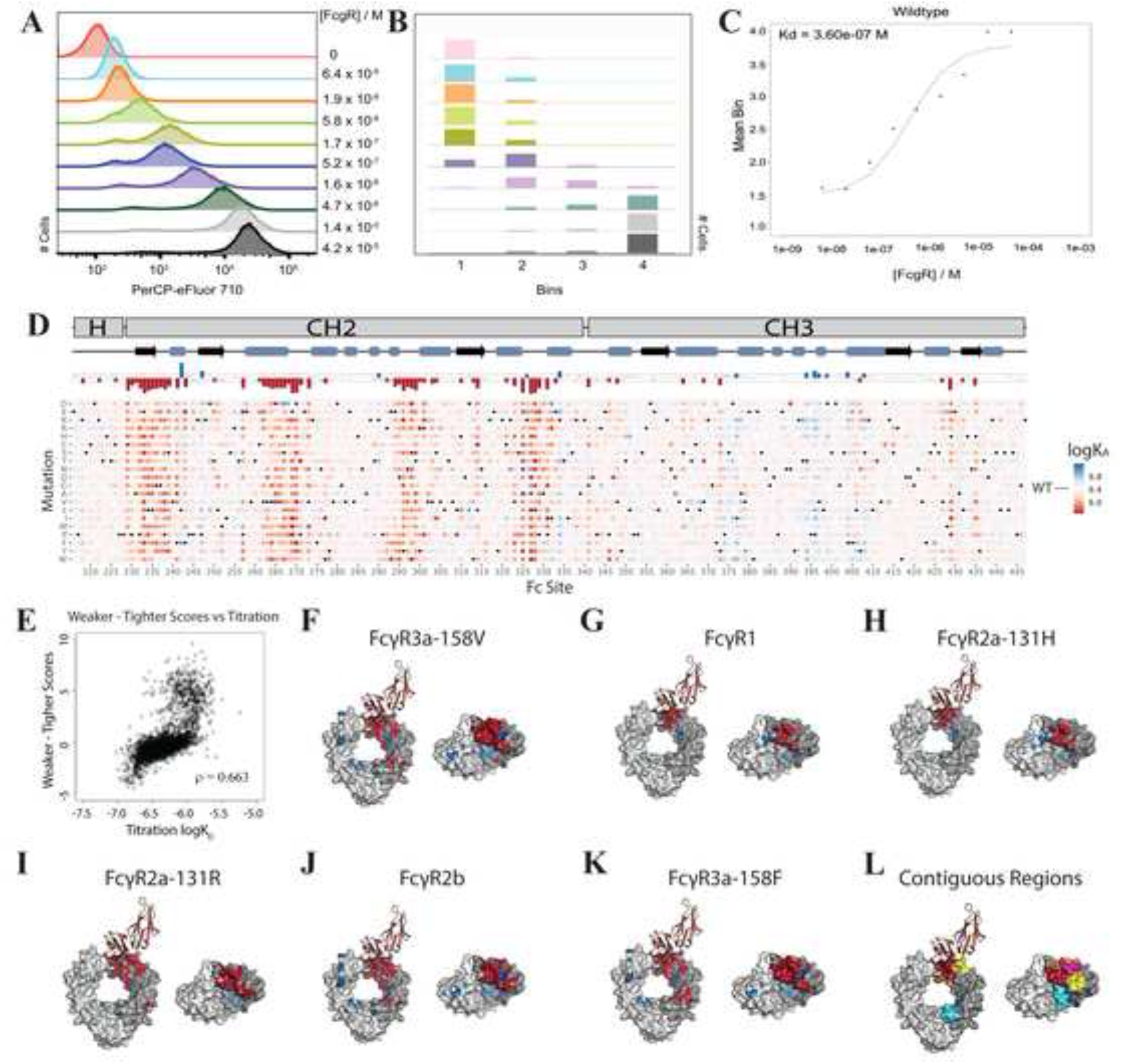
IgG1 Fc deep mutational scanning titrations. **(A)** A shift in fluorescence is observed with increased FcγR3a-158V concentration. **(B)** Panel (A) is reflected by the number of cells collected in each of the 4 bins. **(C)** A mean bin can be calculated against each concentration and a curve fitted per barcode. Effective *KD*s can be calculated from these curves and a weighted average used to calculate the *KD* (or K*A*) per variant. **(D)** Heatmap of deep mutational scanning titration for IgG1 Fc mutants with FcγR3a-158V. Average log*KA* scores per site are depicted on a bar chart above the heatmaps, with sites highlighted if the score exceeds 0.75 standard deviations above (blue) or below (red) the WT log*KA*. **(E)** Scatterplot showing the correlations between variant weaker minus tighter binding values and titration log*K*Ds for experiments with FcγR3a-158V. Average log*KA* values mapped onto Fc dimer structures for **(F)** FcγR3a-158V, **(G)**FcγR1, **(H)** FcγR2a-131H, **(I)** FcγR2a-131R, **(J)** FcγR2b, and **(K)** FcγR3a-158F. **(L)** Titration data revealed four sequence-defined contiguous regions where point mutations typically result in weaker Fc-FcγR binding, and two regions which typically enhance binding. These regions are color-coded on a representative Fc structure (which otherwise consists of a grey surface) bound to FcγR3a-158V. Regions giving weaker binding: sites 230-240, red; sites 262-272, pink; sites 291-300, orange; and sites 323-331, yellow. Regions giving stronger binding: sites 370-379, dark blue; and sites 394-407, cyan.

These effective affinities are in broad agreement with the corresponding tighter and weaker binding scores for each Fc-FcγR interaction (**Figure S3**). Therefore, the general trends identified in the analysis using a single concentration of FcγR largely hold when considering these titrations. Our high-resolution deep mutational scanning titration analysis revealed four contiguous sequence-based regions of human IgG1 Fc where mutations typically lead to weaker binding to all human FcγRs, encompassing approximately sites 230-240, 262-272, 291-300 and 323-331, inclusive of the lower hinge region and the three Fc loops that all contain at least some residues that make direct contacts with FcγRs. We also identified two Fc regions that typically result in stronger FcγR binding, consisting of sites 370-379 and 394-407, both of which are located in the Fc CH3 domain and span the distance from the CH2-CH3 joint to the CH3-CH3 interface. All regions that commonly decrease or increase Fc binding to all FcγRs are depicted in **Figure 5L**. In general, the Fc regions that typically reduce binding affinity upon mutation are situated at or near the Fc-FcγR interface, while the Fc regions that typically enhance binding upon mutation, conversely, are situated far from the Fc-FcγR interface.

In addition to these general trends, our deep mutational scanning titrations also allowed us to identify Fc variants that most increase or decrease binding to each of the human FcγRs. Effective log*K*_A_ values for each Fc variant with every FcγR are provided in GitHub (https://github.com/Ortlund-Laboratory/DMS_IgG1Fc/tree/main/Deposited_Data/Titration_Data). Additionally, the top ten strongest- and weakest-binding variants for each receptor are provided in **Table S1**. Some of these strongest/weakest variants are consistent with those reported previously, such as G236A,^40^ S267E,^41^ L328F^41^ and H268D.^42,43^ Furthermore, some of these strongest/weakest variants were common amongst more than one receptor. Of the variants that most influenced effective Fc-FcγR affinities to three or more receptors, two resulted in weaker binding (E269P and N297A) and four resulted in stronger binding (S239D, I332D, K334E and P396I). As mentioned above, N297 is an *N*-linked glycosylation site required for FcγR binding and induction of antibody-mediated effector functions, and our titration data do indeed show reduced binding upon mutation at this site. Of the 42 unique variants that appeared in the top ten strongest-binders against the respective receptors, 15 belonged to one of the four regions identified as typically reducing binding upon mutation. That is, these 15 variants each contain mutations at the Fc-FcγR interface, indicating that even though most mutations in this protein-protein interface reduce binding, there are numerous mutations that instead significantly increase affinity.

### Comparing the effects of Fc mutations across all low-affinity FcγRs

An important advantage of generating log*K*_A_ values for all Fc mutations is that it allows us to directly compare the results between FcγRs. The relative effects of individual mutations on increased or decreased binding to multiple receptors can be calculated as long as the results for each receptor are normalized to a consistent value for wild type Fc binding (here we set this value to 0). This normalization step allows us to plot ‘differential titration heatmaps’ that depict the relative effects of all mutations on binding to one selected FcγR versus binding to any other FcγR or FcγRs. The key parameter for how Fc mutations drive antibody-mediated effector functions is their relative affinities for activating (FcγR2a and FcγR3a) versus inhibitory (FcγR2b) low-affinity FcγRs, typically expressed as a value equivalent to the ratio of affinities to one inhibitory FcγR versus that to one activating receptor, or the I:A ratio. As FcγR2b is the only inhibitory FcγR, we plotted differential heatmaps for each of the activating FcγRs versus FcγR2b (**Figure S4**), as shown for FcγR3a-158V (**Figure 6A**). We mapped differential effective affinities between activating FcγRs and FcγR2b onto the respective activating Fc-FcγR complexes (**Figure 6B-E**). It is evident from these data that Fc mutations that result in substantial relative increases and decreases in binding to both the activating receptor and to the inhibitory receptor extend from the Fc-FcγR interface, through the Fc lower hinge and CH2 domain, and into the CH3 domain and CH3-CH3 interface. We generated tables (https://github.com/Ortlund-Laboratory/DMS_IgG1Fc/tree/main/Deposited_Data/Differential_Titration_Data) showing all (FcγR2b – FcγR2a-131H; FcγR2b – FcγR2a-131R; FcγR2b – FcγR3a-158F; and FcγR2b – FcγR3a-158V) differential log*K*_A_ scores and ordered each from lowest to highest I:A ratio. Finally, we generated differential heatmaps and mapped differential effective affinities comparing the major activating receptor polymorphisms - FcγR2a-131R/FcγR2a-131H and FcγR3a-158F/FcγR3a-158V (**Figure S5**).

**Figure 6.**
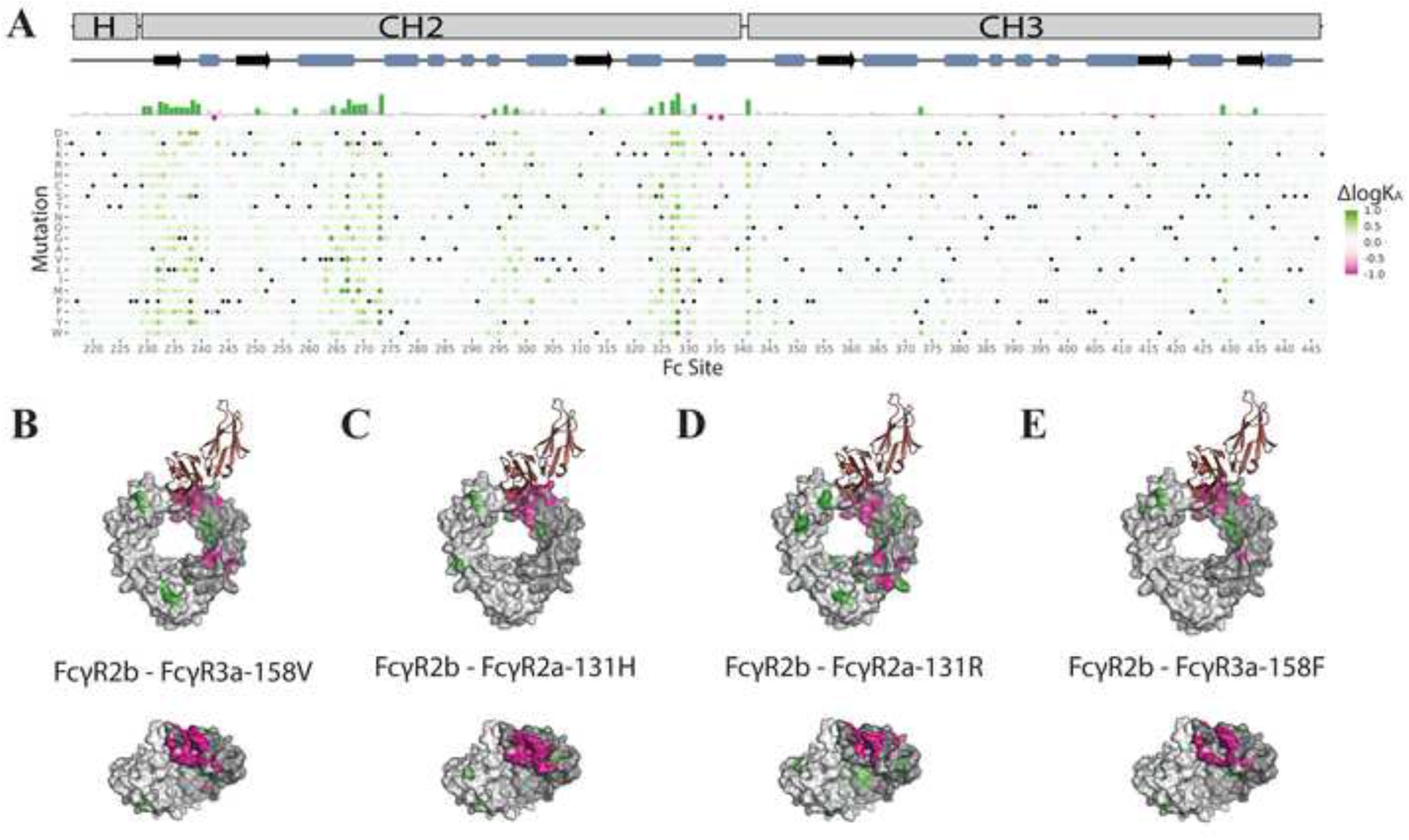
Differential titration data for FcγR2b versus low-affinity activating FcγRs. **(A)** Heatmap showing the difference in log*KA* values for all single-point variants of FcγR2b versus FcγR3a-158V. The color scale is normalized to the difference in wild type log*KA* values for FcγR2b and the selected receptor, respectively. Positive scores (green) indicate stronger binding to FcγR2b, and negative scores (pink) indicate stronger binding to the other receptor. Average scores per site are given in the bar chart above the heatmap. Per-site average scores exceeding 1.0 standard deviation above (green) or below (pink) the mean are highlighted on Fc dimer structures. Average differential logKA values mapped onto Fc dimer structures for **(B)** FcγR2b-FcγR3a-158V, **(C)** FcγR2b-FcγR2a-131H, **(D)** FcγR2b-FcγR2a-131R, and **(E)** FcγR2b-FcγR3a-158F.

We also identified Fc mutations from our deep mutational scanning titration dataset that exhibited the highest (i.e., most inhibitory) or lowest (i.e., most activating) I:A ratios. To determine which variants broadly cause the greatest increase/decrease in the I:A ratio, we averaged and resorted the results from these four differential tables – one for each activating FcγR, with the exception of the high-affinity activating receptor FcγR1. We further identified those Fc mutants for which the relative log*K*_A_ was within a factor of 2 of the average, the purpose of which was to ensure that the variant had approximately the same effect on each activating receptor (see https://github.com/Ortlund-Laboratory/DMS_IgG1Fc/blob/main/Deposited_Data/Differential_Titration_Data/Collected_

<u>Differential_Titration_Data/sorted_Collected_Titration_Data_Github.xlsx</u>). In addition to these detailed tabulations, we provide the ten variants that each exhibit the highest and lowest I:A values in **Table S2**, which we mapped to the Fc-FcγR structure in **Figure S6**. The positions of these Fc mutants indicate that those that should be most activating (i.e., lowest I:A) typically clustered at the Fc-FcγR binding interface, whereas those that should be most inhibitory (i.e., highest I:A) are dispersed throughout the Fc and often distal to the Fc-FcγR interface.

### Biophysical validation of deep mutational scanning data

It is evident that our deep mutational scanning experiments faithfully identified the binding interface on IgG1 Fc for each of the FcγRs, as the majority of Fc mutations that led to weaker FcγR binding were clustered in the known Fc-FcγR interaction sites (**Figure 4C&F-J**). To further validate the effective affinities of Fc-FcγR interactions derived from our deep mutational scanning data, we performed SPR analysis to measure the binding affinities of a set of Fc mutants to each of the low-affinity FcγRs. First, we expressed recombinant wild type IgG1 Fc, captured the Fc directly from the culture supernatant onto protein A-immobilized SPR sensor chips, individually flowed titrated concentrations of FcγR2a-131H, FcγR2a-131R, FcγR2b and FcγR3a-158F as analyte, and analyzed the resulting sensorgrams using a steady-state affinity model due to the fast association and dissociation rates of these low-affinity interactions (**Figure 7A**). Our measured wild type Fc-FcγR affinities are similar to SPR measurements reported by others^35^ (also see **Figure 1D**). Then, from the Fc mutants that exhibited the highest and lowest I:A ratios (see **Table S2**), we selected twelve single-site Fc mutants – P238F, L242F, S267Q, H268E, V273F, V273Q, N325M, I332R, S337G, D376A, N389T and V397L – and measured their affinities to FcγRs as described above for wild type Fc. We plotted the log*K*_A_ values derived above from deep mutational scanning experiments versus those derived by our SPR analysis (**Figure 7B**). For each of these mutant Fc-FcγR interactions, relative to its respective FcγR2b interaction, we calculated and compared the I:A ratio derived from deep mutational scanning analysis to those derived from SPR analysis (**Figure 7C**). All of these data suggest a strong correlation between effective affinities determined by deep mutational scanning experiments and affinities determined by SPR analysis. Additionally, we confirmed by differential scanning fluorimetry that none of the above IgG1 Fc variants exhibited substantially different thermostabilities relative to wild type (see **Table S3**).

**Figure 7.**
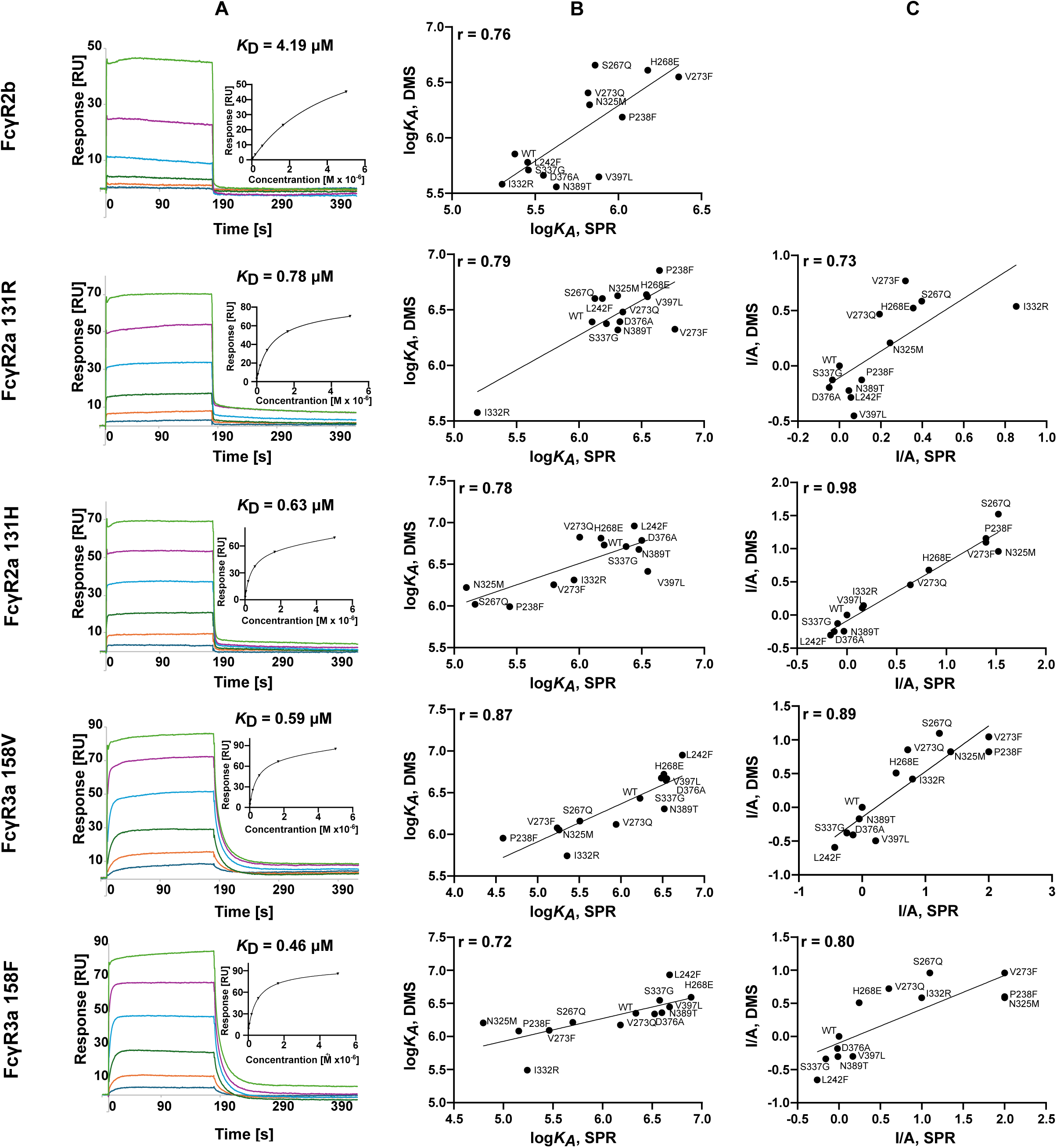
Biophysical validation of deep mutational scanning data by surface plasmon resonance (SPR) analysis. **(A)** Representative SPR data for affinity measurements between wild type IgG1 Fc and FcγR receptors. Sensorgrams represent injections of FcγRs at concentrations 5 µM, 1.67 µM, 0.56 µM, 0.19 µM, 0.06 µM, and 0.02 µM. Affinity plots used to calculate *K*D values are shown in the insets. **(B)** Comparison of log*K*A values determined by deep mutational scanning titrations and SPR analyses. Pearson correlation coefficient (r) was used to estimate similarity (p < 0.05). **(C)** Comparison of I:A values determined by DMS and SPR. Pearson correlation coefficient (r) was used to estimate similarity (p < 0.05).

### Allosteric effects on Fc-FcγR binding are localized to discreet areas of the Fc region

Inspection of logK_A_ values mapped onto structures (for example, in **Figures 5** and **S2**) suggests that the control that Fc variants exercise over binding affinity to the various receptors is strongly dependent on the positions of the mutated residues. We therefore charted a systematic analysis of this phenomenon (**Figures S7** and **S8**), showing that the Fc residues in closest proximity to, or in direct contact with, the receptors typically exert the strongest impact on FcγR binding. The region with the next highest impact on FcγR binding, however, was not the layer immediately adjoining residues with direct contact, but instead the layer 50-60 Å from the receptor center of mass. This layer comprises the residues at the CH2-CH3 interface on each of the Fc monomers, as well as the inward-facing residues at the Fc dimer interface (**Figure 8A**). These residues are integral to the structure of the Fc region, and the observed changes to binding affinities suggests that such structural changes are exerting allosteric effects on the Fc-FcγR binding interface. Importantly, this layer would not have been identified by approaches relying purely on structural analyses of the Fc-FcγR interfaces highlighting the power of our unbiased mutational scan for identifying interaction hot spots between two proteins. The average effect on Δlog*K*_A_ within this 50-60 Å layer is either positive or marginally negative, suggesting that this layer may be particularly promising for engineering Fc variants with differential affinities to activating and inhibitory FcγRs, alongside those at the Fc-FcγR binding interface (the 20-30 Å layer).

**Figure 8.**
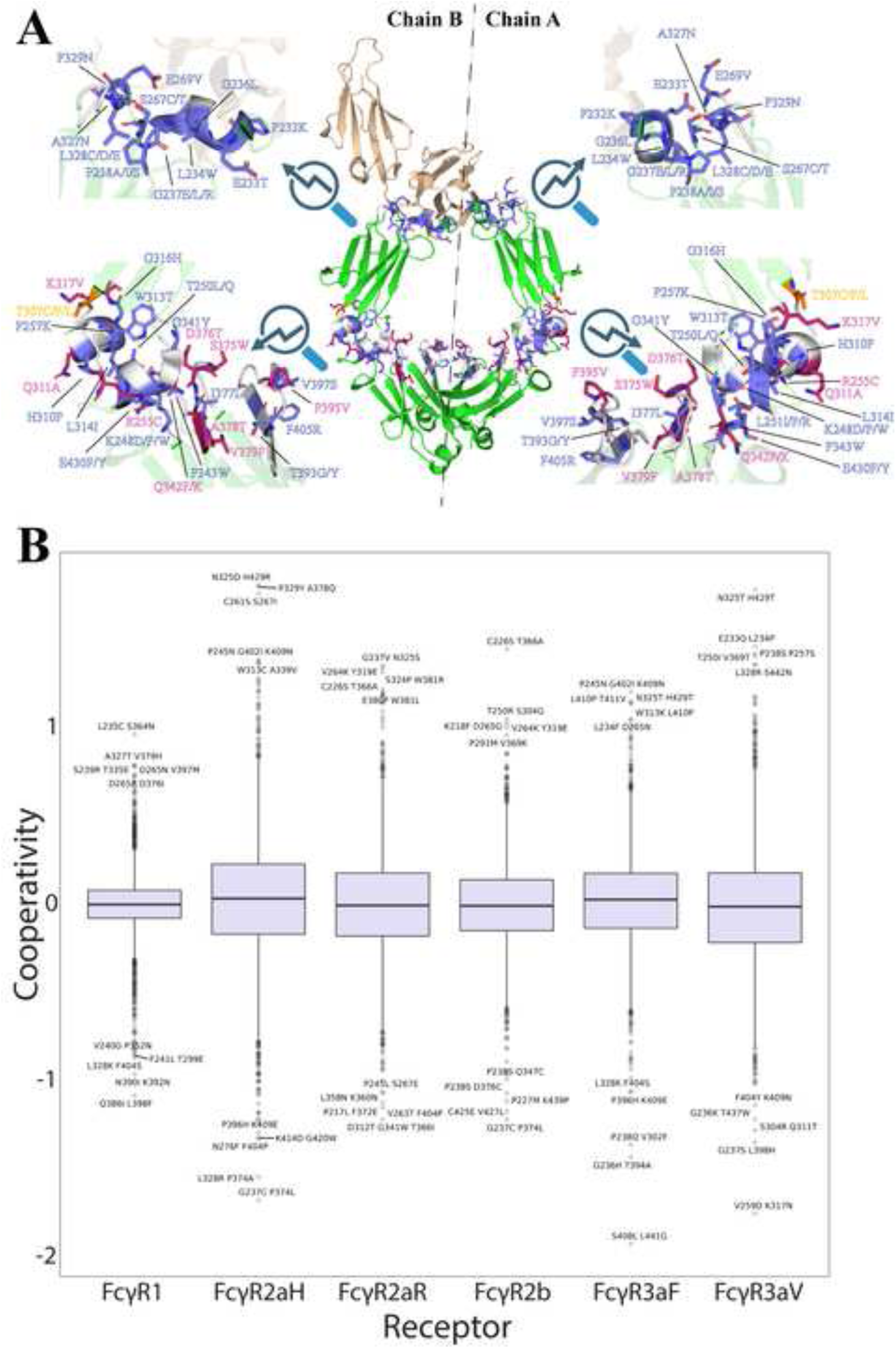
Mapping the locations of allosteric and cooperative Fc mutations within the Fc region. **(A)** The FcγR is represented in tan; Fc in green. Residues within the 20-30 Å and 50-60 Å are color-coded in white, by default. If a residue can be mutated to give an Fc with an increased log*KA* to FcγR2b whilst simultaneously giving a decreased log*KA* against the activating receptors (FcγR2aH, FcγR2aR, FcγR3aF and FcγR3aV), then the residue is colored blue and the wild type stick model explicitly shown. Residue labels give the amino acid(s) which produce this effect. If the mutated residue results in decreased binding affinity to FcγR2b and increased binding affinity to all of the activating receptors, the same applies but the residue is colored magenta instead. T307 was the only residue which showed both trends described above, depending on which amino acid it was mutated to, and so is colored orange. If converted to cysteine, binding to all activating receptors is increased and to FcγR2b is decreased, and if converted to phenylalanine or leucine, binding to FcγR2b is increased and to all activating receptors is decreased. Both chain A and chain B of Fc are shown. **(B)** Positive cooperativities indicate multi-site variants (e.g., L235C S346N) where the Δlog*KA* value exceeds the sum of the Δlog*K*A values of its constituent, single-point mutant, elements (e.g., L235C and S346N), whereas negative cooperativities indicate that the sum of the Δlog*K*A values of individual mutations exceed that of the corresponding multi-site variant. Cooperativity values close to 0 indicate additivity. The bold lines within each box defines the median, and the boxes themselves capture the interquartile range. The upper/lower whiskers extend to 1.5 times the interquartile range over the 75^th^ percentile/under the 25^th^ percentile. The outliers above/below these whiskers are plotted as individual points, and are labeled if they are ranked within the top five most/least cooperative variants.

We therefore extracted the Δlog*K_A_* values against each receptor for each mutation from all of the residues within these two layers (see https://github.com/Ortlund-Laboratory/DMS_IgG1Fc/tree/main/Deposited_Data/ΔlogKa_vs_Distance). Since FcγR2b is the only inhibitory receptor, we identified mutations that result in a positive Δlog*K*_A_ against this receptor while producing a negative Δlog*K*_A_ against all of the activating receptors (i.e., which increase I:A) and those that result in a negative Δlog*K*_A_ against FcγR2b while producing a positive Δlog*K*_A_ against all of the activating receptors (i.e., which decrease I:A) (**Figure 8A**). **Table S4** lists these mutations, alongside the difference between Δlog*K*_A_ to the inhibitory receptor and Δlog*K*_A_ to the average of the four activating receptors. Thus, this metric indicates mutations that are likely to enhance inhibition (a large, positive score) or activation (a large, negative score). Inspection of these data reveals that the mutation predicted to most enhance inhibition (i.e., which has the largest positive score) is P238S, although deep mutational scanning data for this mutation with FcγR2aH and FcγR3aF are missing. This was due to this particular mutation only being associated with one barcode in our library, thus increasing the likelihood for failing the stringent quality control steps we have in place for our curve fitting procedure. The mutation producing the next best enhancement in inhibition is F405R. This residue is situated at the interface between the two Fc chains, far from the receptor, suggesting a significant allosteric effect. All mutations identified as improving activation were situated in the layer 50-60 Å from the receptor center of mass. Of these, D376T was most effective at improving activation and appears to form part of a larger network of residues that could be mutated to improve activation (S375W, D376T and P395V, respectively, **Figure 8A**). These residues all point inwards (i.e., towards the receptor) and are proximal to the interface between the two Fc chains. T307 was the only residue that met both metrics: if converted to a cysteine, activation is improved; if converted to a phenylalanine or a leucine, inhibition is improved – suggesting this residue as a promising target for engineering. T307 is situated at the interface between the CH3 and CH2 domains, suggesting this portion of the Fc exerts allosteric control on receptor binding. We further found that all 18 mutations within the 20-30 Å layer improve inhibition, while none improve activation, suggesting that the residues at the Fc binding interface may have specifically evolved to bind to activating receptors rather than the inhibitory receptor.

### Cooperative effects of combinatorial Fc mutations

Our deep mutational scanning libraries contained numerous combinatorial mutations in addition to the nearly comprehensive single-site changes. Though a small fraction of the theoretical number of combinatorial mutations (see **Table S5** for the statistics), we hypothesized that this wealth of additional data could be used to measure cooperativity effects between mutations at pairs of distinct IgG1 Fc positions. These extra data could potentially be included as input for a machine learning approach to predict the effect of higher-order mutations on binding affinities at scale.

From each titration, we extracted all of the multi-mutation variants, along with their associated log*K*_A_ values. We then subtracted the log*K*_A_ value for the wild type against the relevant receptor since this normalizes the data, giving Δlog*K*_A_ values, where positive values now indicate that the mutation causes an increase in binding affinity and negative values a decrease in binding. We compared these data to the sum of the normalized single-point mutations constituting the variant (for example, by comparing the Δlog*K*_A_ value for the T394W R416A variant against the sum of the Δlog*K*_A_ values for the T394W and R416A variants, respectively). We achieved this comparison by subtraction of the single-point data from the multi-point data, so that a positive value indicates positive cooperativity between the mutated residues, a value approximately equal to 0 suggests strict additivity, and a negative value suggests negative cooperativity, which we plotted for each receptor (**Figure 8B**). These data indicate that the median cooperativity is very close to 0 for each receptor. Furthermore, the interquartile ranges for each receptor do not extend far from 0, indicating that the great majority of variants are simply additive. The interquartile range, whiskers and outliers are narrower for FcγR1 than for the other receptors. FcγR1 is known to bind to the opposite face of the same contact plane of Fc compared to the other receptors^44^ (for example, compare the crystal structures 4W4O^44^ and 3SGJ,^39^ or see our Alphafold simulations in **Figure S8**) and it could be that mutations at this face are generally less able to exert a significant effect upon binding, thus providing less scope for cooperative behaviors. There are, however, numerous outliers from these box plots, some of which exhibit cooperativity values approaching +2, and others approaching -2, which represent potential new targets for Fc engineering. As Δlog*K*_A_ values are being compared, this implies that careful consideration of mutational combinations can give up to two orders of magnitude changes in binding affinities in both directions compared to what would be expected from simple additive effects of single mutations, thus confirming the importance of epistatic effects to any Fc engineering strategy.

We then extended our analysis to determine whether cooperativity was influenced by (1) the distance of the mutants from the receptor and (2) the distance of the mutated residues from each other. We limited our study to double mutants (which we justified from **Table S5**, which shows that higher-order mutations only constitute a small fraction of the variants) in order to simplify the analysis. For each double mutant, we quantified the distance between the center of mass of the receptor from the center of mass of each changed residue. Because the mutated residues appear on both chains of Fc, and these chains bind asymmetrically to the receptor, we averaged the distances between chains. This strategy gives two distance metrics per double mutant, which constitute our binning strategy, as depicted in **Figure S9A** and in the respective plots of **Figure S9B-H**. While **Figure S9B** gives the data summed over all receptors, **Figure S9C-H** gives the data separated by receptor. The latter data were explicitly plotted in case any of the receptors acted differently to the overall trends; however, this proved not to be the case. The bars in **Figure S9B-H** are subdivided according to whether the double mutation is positively cooperative, simply additive, or negatively cooperative. Left-hand plots give the number of times a double mutant occurs in a particular bin, whereas right-hand plots give percentage values. These data show that the distances of the mutated residues from the receptor generally have little effect on the likelihood of cooperativity, demonstrating that there is no discreet Fc region (such as the interface between its respective chains, or its interface with the receptor) that is particularly prone to cooperativity.

## DISCUSSION

The interactions between IgG antibodies and FcγRs are of critical importance to the human immune system, both in its physiological function and its pathological dysfunction. Although Fab regions of these antibodies are needed for antigen recognition and neutralization, their Fc regions are responsible for all of their signaling capabilities. Without Fc regions, IgG antibodies are effectively inert binding molecules, able only to engage antigens and, in some instances, neutralize pathogens. Accordingly, certain bacteria that are pathogenic in humans have evolved enzymes that specifically degrade or deglycosylate IgG antibodies in order to defeat effector functions, and an entire class of IgG degrader drugs have already entered the clinic for treating a wide range of IgG-mediated pathologies.

The antibody-mediated effector functions of IgG antibodies that are the result of Fc-FcγR interactions are regulated by the binding of the IgG Fc region to both activating – FcγR2a and FcγR3a – and inhibitory – FcγR2b – low-affinity receptors. The balance of IgG Fc binding to inhibitory versus activating FcγRs ultimately dictates the strength of the inflammatory immune response, which can be expressed as the inhibitory:activating, or I:A ratio. Changes in the Fc regions of IgG antibodies in the periphery, such as modifications to the Asn297-linked glycan, can shift the balance of Fc binding to inhibitory versus activating FcγRs, thereby resulting in either more or less immune stimulation. For instance, a relative increase in afucosylated IgG1 antibodies in severe dengue disease triggers antibody-dependent enhancement (ADE) mediated through increased affinity to FcγR3a that correlates with morbidity and mortality in these patients.^45^ Such manipulation of immune signaling can be leveraged for therapeutic monoclonal antibodies (mAbs) for indications in which elevated antibody-mediated effector functions are beneficial; there are already seven FDA-approved afucosylated IgG mAbs.

The other major route to engineering IgG mAbs with bespoke antibody-mediated effector functions is to mutate the IgG Fc region such that it will preferentially bind inhibitory versus activating FcγRs, or vice versa, to manipulate the I:A ratio. For decades, Fc mutagenesis campaigns, both large and small, have been launched with the goal of improving the effectiveness of therapeutic mAbs. An early mutagenesis campaign was successful in determining the Fc binding site common to all FcγRs, which in turn led to the discovery of mutations that increased ADCC up to 2-fold.^46^ Since then, high-resolution crystal structures of many IgG Fc-FcγR complexes became available,^42,47,48^ further indicating which Fc residue interactions are common between certain receptors, and which are unique. Despite this additional information, screening strategies have typically still required the investigation of a vast number of variants. Notable campaigns include Chu *et al.*,^41^ who screened over 900 variants, ultimately identifying S267E/L328F as a mutational combination which enhances binding to FcγR2b 430-fold whilst acting minimally on FcγR1 and FcγR2a-131H, and eliminating binding to FcγR3a-158V; Mimoto *et al.*,^42^ who screened over 500 variants, to discover that the P238D mutation enhances binding to FcγR2b whilst simultaneously severely reducing binding to all of the activating FcγRs studied; and Richards *et al.*,^40^ who also screened over 900 variants, uncovering a G236A variant which improves ADCP through enhanced binding to both allotypes of FcγR2a, has a minimal effect on ADCC or FcγR3a binding, but also reduces binding to FcγR1, resulting in a suboptimal therapeutic capacity. Additional mutagenesis efforts have yielded combinatorial variants that can have even larger impacts on the I:A ratio, such as GAALIE (G236A/A330L/I332E),^49^ which preferentially promotes binding to activating FcγRs (relative fold increases of FcγR binding), as well as V11 (G237D/P238D/H268D/P271G/A330R) and V12 (E233D/G237D/P238D/H268D/P271G/A330R), which increase FcγR2b affinity compared to wild type 40-fold and 217-fold, respectively.^42,43^

Thus, even with a deep understanding of the sequences and structures of both IgG Fc regions and FcγRs, rational manipulation of inhibitory and activating Fc-FcγR interactions to shift the I:A ratio remains a major challenge. While all Fc-FcγR complexes are nearly indistinguishable from the structural standpoint, their biological effects are driven by differences in these interactions that are not inherent readouts of even the state-of-the-art biophysical and biological methods used to evaluate them. Indeed, several properties of Fc-FcγR interactions that are difficult to measure, including but not limited to the inherent conformational dynamics of the IgG Fc region, allosteric control of Fc-FcγR interactions, and cooperativity between Fc residues in their effects on FcγR binding, have prohibited a full understanding of these interactions.

We therefore devised a deep mutational scanning approach for interrogating Fc-FcγR interactions to provide a comprehensive and systematic analysis to better evaluate parameters governing Fc-FcγR complex formation and selectivity of binding to inhibitory versus activating FcγRs. In our approach, we constructed a library of nearly all possible human IgG1 Fc mutations, expressed these mutants on the surface of HEK293T cells, and sorted these cells on the basis of binding to each human FcγR and the most common FcγR polymorphisms. Using a variety of sorting methods, we were able to generate comprehensive datasets that address critical properties of Fc-FcγR interactions, such as stability, affinity, allostery and cooperativity.

As with other mutagenesis campaigns, the primary readout of our deep mutational scanning analysis was Fc-FcγR affinity. One major difference with previous mutagenesis efforts is the scale of our approach. We were able to evaluate virtually every possible amino acid substitution at every position of the IgG Fc region – well over 4000 total mutations. Not only does this provide a map of affinity throughout the entire Fc for binding to each FcγR, the unprecedented comprehensiveness of our data allows us to identify previously unrealized trends in Fc-FcγR affinity. For instance, we found that the Fc regions that typically enhance binding to all FcγRs upon mutation, somewhat counterintuitively, are situated far from the Fc-FcγR interface. Furthermore, these regions are located at the interface between the two chains of the dimer and are distal to the receptor, suggesting that there are mutations that strengthen the interaction between these chains, which in turn stabilizes and/or reorients the dimeric structure, thereby allowing the receptor to bind more effectively at the Fc-FcγR interface.

Because the main biological driver of antibody-mediated effector functions is the differential affinity of the Fc region for inhibitory versus activating FcγRs, we used a binning and sorting strategy with our deep mutational scanning library to produce effective affinities for all single-site Fc mutations that allowed us to directly compare affinities across any and all Fc-FcγR interactions. We identified not only those single-site Fc mutations that have already been reported to substantially influence the I:A ratio in both directions, but also many more. Again, due to the comprehensiveness of our mutagenesis data, we were able to identify general trends in differential affinities across Fc-FcγR interactions. Most notably we found that Fc mutations that most strongly affected the I:A ratio were located entirely outside of the Fc-FcγR binding interface and, furthermore, that those that increased and decreased the I:A ratio most clustered to discrete locations on the Fc region structure.

By sorting the cells in our deep mutational scanning library for the absence of the marker of cell-surface expression of the Fc region, we were able to produce a complete map of single-site mutations that affected the stability and/or trafficking of IgG1 Fc. The Fc region is an obligate homodimer that is primarily stabilized by the CH3-CH3 interface, yet we found that the CH3 domain is much more amenable to mutation than the CH2 domain. While there is no real CH2-CH2 interface – these domains are primarily tethered by the disulfide bonded hinge – our data indicated that the CH2 domain has many more sites than the CH3 domain that are intolerant of mutation and, of those, they are much less tolerant of the types of amino acids that can be substituted.

The IgG Fc region is well known to be conformationally dynamic. Crystallographic snapshots have identified “open” and “closed” conformations; NMR data has highlighted even greater flexibility of this protein.^32^ While the role of the Asn297-linked glycan – especially afucosylated and asialylated glycoforms – in allosteric control of Fc-FcγR binding and the resulting antibody-mediated effector functions has been shown previously, our deep mutational scanning analysis data provides new insights into the role of all single-site amino acid substitutions in the allosteric control of Fc-FcγR interactions.

Despite the power of site-saturation mutagenesis to inform protein-protein interactions, there are clear limitations to the data produced by a collection of single-site mutants, even if that collection includes all such mutations that are possible. Especially in allosterically regulated binding events, individual amino acids in a protein sequence do not function in isolation but, rather, in cooperation with other amino acids in the same protein. Because the fidelity of the synthesis of our site-saturation library was not perfect, in addition to all of the single-site Fc mutant library, numerous wild type and double mutant sequences are present in our library that we barcoded and identified in various sorted populations. The wild type sequences act as internal controls in our deep mutational scanning analysis experiments, while the double mutants serve as an unbiased collection of combinatorial variants that provide unique insight into the degree of cooperativity for pairs of mutations at different sites in the Fc region and their effect on FcγR binding.

In summary, we developed a cell-surface displayed deep mutational scanning library for evaluating the effects of virtually all possible single-site and approximately 1% of all possible double mutants of the human IgG1 Fc region to all human FcγRs. Our data provide the most comprehensive analysis of Fc-FcγR interactions to date, as well as unprecedented insights into how the interactions between the Fc region and activating and inhibitory FcγRs are regulated by allostery.

### Limitations of the Study

Although our deep mutational scanning library of Fc mutants is a nearly comprehensive set of single-site mutants and many double mutants, it is not a truly combinatorial library that includes a substantial number of higher-order combinatorial variants. Due to the conformational flexibility in the Fc region and allosteric regulation of Fc-FcγR interactions, such a combinatorial library could provide greater insight into these interactions; however, there are technical hurdles to displaying what would be a considerably larger library on the mammalian cell surface. Also, our cell surface display system relies on the endogenous glycosylation machinery of the mammalian cell on which library members are displayed, which results in Fc regions that are glycosylated in a consistent but limited way. Since Fc glycosylation is a major determinant of FcγR binding, we can only evaluate our Fc mutants in the context of a subset of glycosylation states, or IgG glycoforms. Glycan remodeling on the mammalian cell surface in combination with our deep mutational scanning library could enhance the evaluation of Fc-FcγR by expanding the number of distinct IgG glycoforms from which data could be derived.

## SUPPLEMENTAL INFORMATION

Supplemental information can be found online at DOI: XXXXXXXXXXXXXX

## ACKNOWLEDGEMENTS

Research reported in this publication was supported by the National Institute of Allergy and Infectious Diseases grant R01AI149297 (to E.J.S). A.D.K., M.M.K., F.F. and E.A.O. were supported by National Institute of Biomedical Imaging and Bioengineering of the National Institutes of Health grant 3U54EB027690. Next generation sequencing services were provided by the Emory NPRC Genomics Core which is supported in part by NIH P51 OD011132. Sequencing data was acquired on an Illumina NovaSeq 6000 funded by NIH S10 OD026799. FACS experiments were carried out in the Flow Cytometry Core Facility of the Emory University School of Medicine.

## AUTHOR CONTRIBUTIONS

Conceptualization, F.F., E.A.O. and E.J.S.; Data curation, A.D.K. and T.X.; Formal analysis, A.D.K., T.X., T.C. and E.J.S.; Funding acquisition, E.A.O. and E.J.S.; Investigation, A.D.K., T.X., T.C., M.M.K., M.B., M.N., M.F., T.B. and F.F.; Methodology, A.D.K., T.X., T.C., M.M.K. and F.F.; Project administration, T.C., E.A.O. and E.J.S.; Resources, E.A.O. and E.J.S.; Software, A.D.K.; Supervision, T.C., E.A.O. and E.J.S.; Validation, A.D.K., T.X., M.M.K. and T.C.; Visualization, A.D.K., F.F. and E.J.S.; Writing – original draft, A.D.K. and E.J.S.; Writing – review & editing, A.D.K., T.X., T.C., M.M.K. M.B., F.F., E.A.O. and E.J.S.

## STAR★METHODS

### KEY RESOURCES TABLE

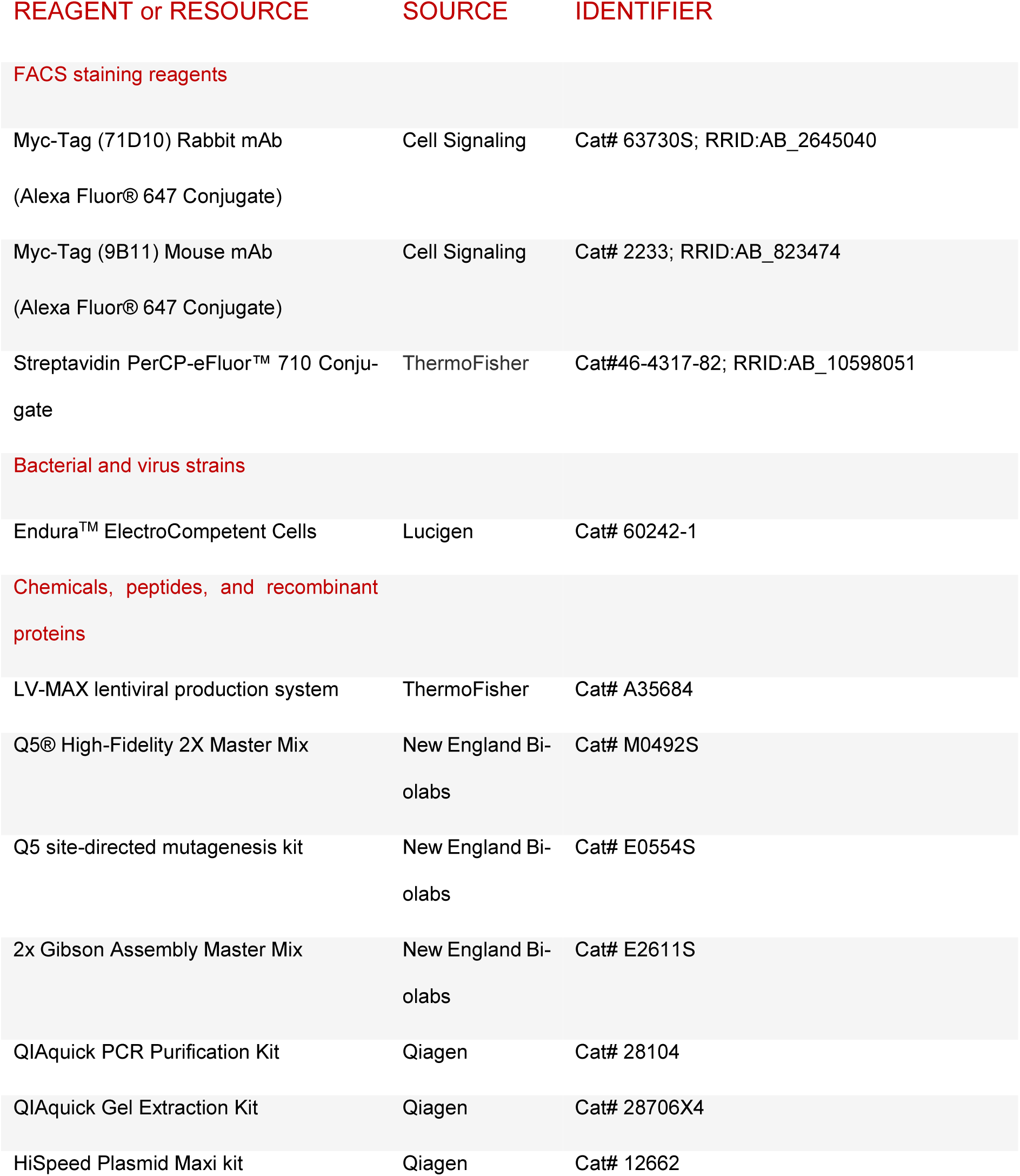
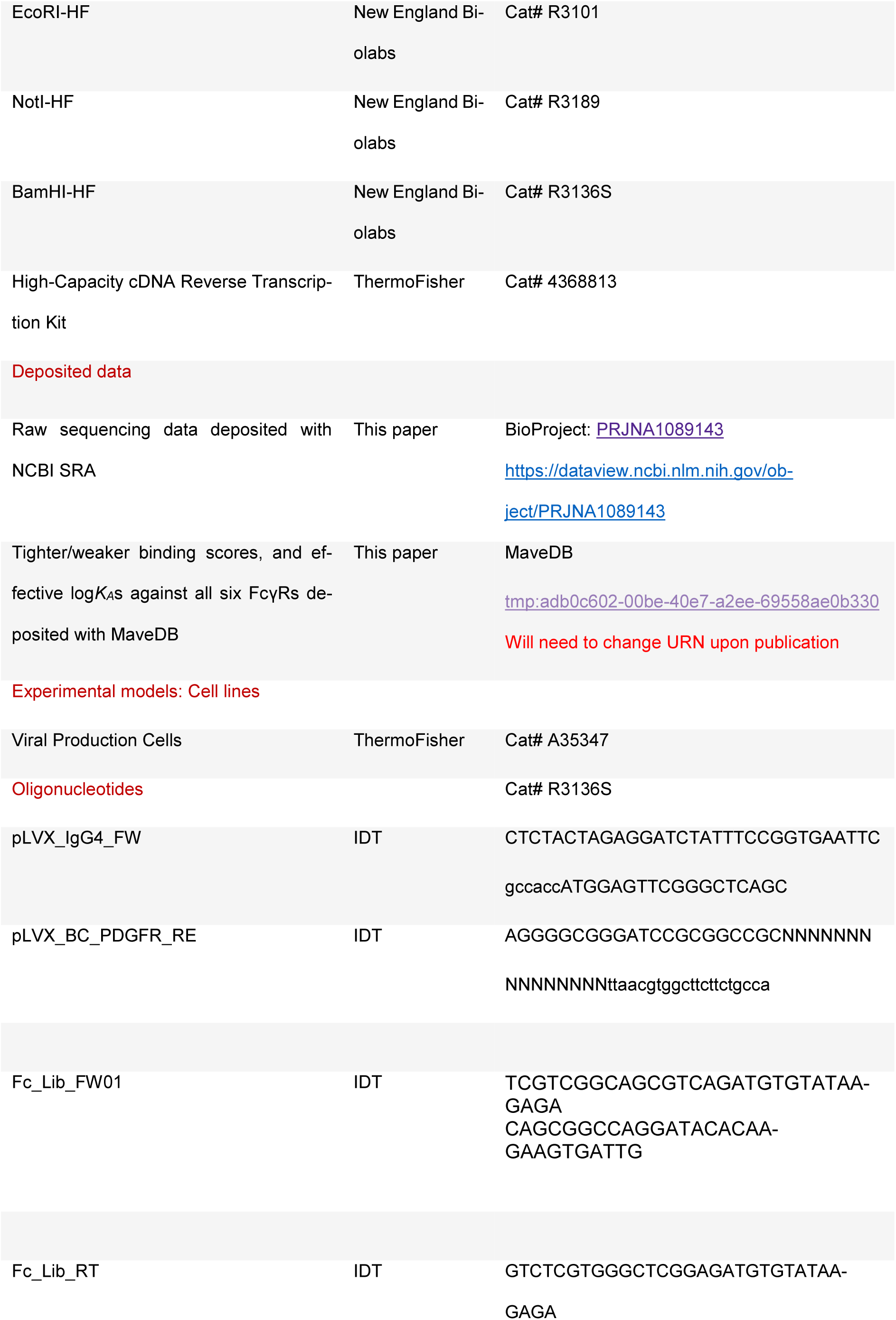
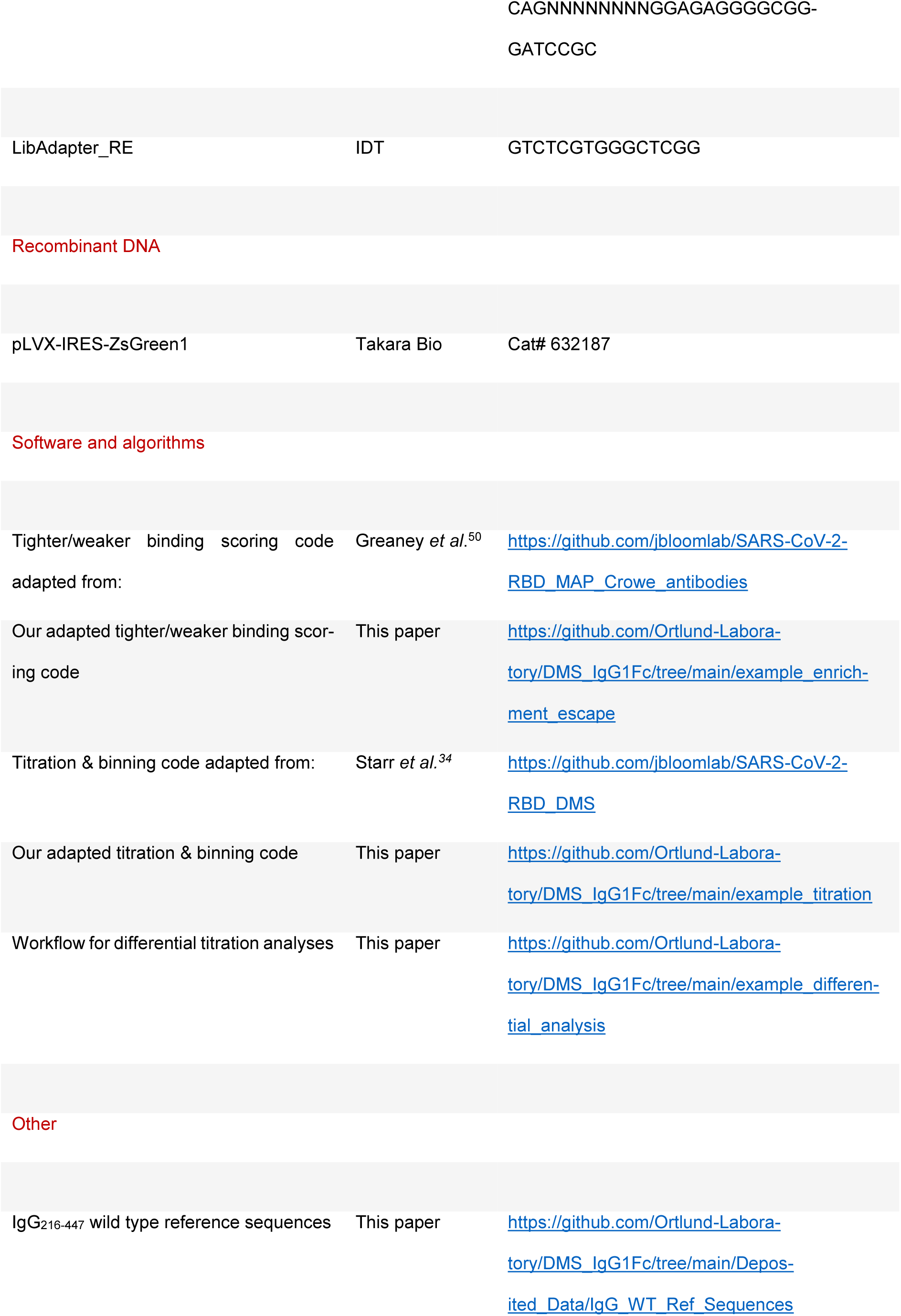

### RESOURCE AVAILABILITY

#### Lead contact

Further information and requests for resources and reagents should be directed to and will be fulfilled by the lead contact, Eric J. Sundberg.

#### Materials availability

IgG1 Fc mutant libraries generated in this study will be made available on request by the **Lead Contact** with a completed Materials Transfer Agreement.

#### Data and software availability

Data and code are provided in the following ways:

- IgG1 Fc coding sequence and mutant codons (https://github.com/Ortlund-Laboratory/DMS_IgG1Fc/tree/main/Twist_Library_QC).
- Raw sequencing data from deep mutational scanning experiments (NCBI BioProject PRJNA1089143, at https://dataview.ncbi.nlm.nih.gov/object/PRJNA1089143).
- The complete PacBio variant:barcode lookup table (https://github.com/Ortlund-Laboratory/DMS_IgG1Fc/tree/main/PacBio_Library_Sequencing).
- Tighter/weaker binding scores for IgG1 Fc against each FcγR (https://github.com/Ortlund-Laboratory/DMS_IgG1Fc/tree/main/Deposited_Data/Enrichment_Escape_Data; and

MAVEdb accession <u>tmp:adb0c602-00be-40e7-a2ee-69558ae0b330</u>). Will need to change MAVEdb accession upon publication of data.

- Effective *K*_A_ values calculated for IgG1 Fc against each FcγR (https://github.com/Ortlund-Laboratory/DMS_IgG1Fc/tree/main/Deposited_Data/Titration_Data; and MAVEdb accession tmp:adb0c602-00be-40e7-a2ee-69558ae0b330). Will need to change MAVEdb accession upon publication of data.
- The complete computational workflow to generate tighter/weaker binding scores (https://github.com/Ortlund-Laboratory/DMS_IgG1Fc/tree/main/example_enrichment_escape).
- The complete computational workflow to generate effective *K*_A_ values (https://github.com/Ortlund-Laboratory/DMS_IgG1Fc/tree/main/example_titration).

### EXPERIMENTAL MODEL AND SUBJECT DETAILS

#### Cells and viruses

All surface-display experiments were performed using “Viral Production Cells” from the ThermoFisher LV-MAX Lentiviral Production system, a derivative of the HEK 293F cell line. Cells were cultured in LV-MAX Production medium which supports growth at high cell densities (up to 10^7^ cells/mL).

### METHOD DETAILS

#### IgG1 Fc cell line generation and mammalian surface display

A codon-optimized IgG1 Fc (residues 216-447; EU numbering, encompassing the hinge and Fc regions) sequence (UniProt ID: P01857) was generated with N-terminal signal peptide sequence derived from IgG4 (MEFGLSWVFLVALFRGVQC), followed by a 1xGGS linker, Myc-tag (EQKLISEEDL), and a 2xGGS linker, as well as a C-terminal 5xGGS linker, followed by a transmembrane helix derived from PDGFR (AVGQDTQEVIVVPHSLPFKVVVISAILALVVLTIISLIILIMLWQKKPR). This sequence was cloned into pLVX-IRES-ZsGreen1 (TakaraBio) at the EcoRI and NotI sites using Gibson assembly (NEB 2x Gibson Assembly Master Mix). The resulting construct was packaged into a lentivirus using the packaging vectors psPAX2 and pMD2G and the LV-MAX lentiviral production system (ThermoFisher) in a 1:1:0.5 pLVX:psPAX2:pMD2G ratio. Lentivirus was titered using the GFP reporter and a stable cell line was generated by infecting Viral Production Cells (from the LV-MAX lentiviral production system) at an MOI of 0.1 so that >90% of cells are infected with a single viral particle. Cells were then sorted for GFP- and Myc-positive-expression using a FACS ARIA II instrument (BD) at the Emory Flow Cytometry Core.

#### Site-saturation library generation

A site-saturation library containing all possible single amino acid mutations at positions 216-447 of the IgG1 protein sequence (UniProt ID: P01857) was synthesized (TwistBioscience). The library was generated as described in Frank *et al*.^33^ Specifically, 15-nucleotide barcodes were added by PCR using 5 amplification cycles using primers pLVX_IgG4_FW and pLVX_BC_PDGFR_RE (see key resources table). The resulting DNA was assembled into pLVX-IRES-ZsGreen at the EcoRI and NotI sites using Gibson assembly (NEB). The Gibson assembly reaction was electroporated using Endura^TM^ ElectroCompetent Cells (Lucigen), plated on LB + Ampicillin plates at an estimated 150,000 colonies per replicate library to limit library complexity, and grown overnight at 30 °C. The next day cells were washed off the plates and plasmids were purified using the HiSpeed Plasmid Maxi kit (Qiagen). The resulting replicate libraries were packaged into a lentiviral library using the packaging vectors psPAX2 and pMD2G and the LV-MAX lentiviral production system (ThermoFisher). Lentivirus preparations were titered using the GFP reporter and stable cell lines were generated by infecting 200 million viral production cells (from the LV-MAX lentiviral production system, ThermoFisher) at an MOI of 0.1 so that >90% of cells were infected with a single viral particle. The cells were harvested 48 hours post-transfection and filtered with 0.45 μM, 25 mm PES filters. Cells were then sorted for GFP-expression using a FACS ARIA II instrument (BD) at the Emory Flow Cytometry Core to collect at least 5 million GFP-positive cells. GFP-positive cells were expanded and cells expressing functional Fc mutants were selected by sorting cells stained with a rabbit Alexa647-anti-Myc antibody (CellSignaling) at a 1:200 dilution (**Figure S2**). At least 5 million GFP-positive and Myc-positive cells were sorted and expanded for each replicate library. These cell lines were then used to screen against FcγRs.

#### Flow cytometry to characterize IgG1 Fc WT binding

Binding to the six FcγRs studied (FcγR1, FcγR2a-131H, FcγR2a-131R, FcγR2b, FcγR3a-158F, and FcγR3a-158V) was individually measured using the cell line stably expressing wild type IgG1 Fc on the cell surface. In a 96-well format, 3-fold dilution series of FcγR was prepared in binding buffer (1X PBS plus 2% FBS and 10 mM HEPES, pH 7.5). Incubation of biotinylated FcγR with ∼5 x 10^4^ cells was carried out at room temperature for 1 hour. Cells were then washed four times with binding buffer, followed by staining with rabbit PE-labeled anti-Myc antibody and PerCP-eFluor™ 710-labeled streptavidin. Data were collected on a Northern Lights (Cytek) instrument equipped with violet and blue lasers. GFP^+^/Myc^+^ - gated cells were analyzed for biotinylated FcγR binding using the median fluorescence intensity (MFI) of PerCP-eFluor710. MFI values were normalized to the highest signal. *GraphPad Prism* was used to plot normalized MFI across 12 wells of serially diluted FcγR concentrations (log10 scale), followed by non-linear regression analysis.

#### PacBio library sequencing and analysis

Inserts containing the IgG1216-447 coding sequences and associated barcodes were cut from the mutational plasmid library *via* EcoRI (*NEB*) and NotI (*NEB*) digestion, gel purified using a QIAquick Gel Extraction Kit (*Qiagen*) and PacBio sequenced (University of Georgia Genomics & Bioinformatics Core). A lookup table linking unique barcodes with their associated mutations was generated with reference to the PacBio circular consensus sequences (CSSs). Identification of the varying IgG1216-447 sequences contained in the CCSs was achieved by first determining the constant regions at the 5’ (containing the SP, Myc and GGS linkers) and 3’ (containing the TM domain and the stop codon) ends.

This varying sequence was then translated and aligned with our IgG1 reference sequence (https://github.com/Ortlund-Laboratory/DMS_IgG1Fc/tree/main/Deposited_Data/IgG_WT_Ref_Sequences). These steps were achieved using the computational notebook created by Starr *et al.*^34^ (available on GitHub at https://github.com/jbloomlab/SARS-CoV-2-RBD_DMS/blob/master/results/summary/process_ccs.md) with minor adaptations. For the alignment step, this notebook uses alignparse,^51^ version 0.6.0, which in turn depends on minimap2,^52^ version 2.17. At this stage, a filtering step was set up so that only CCSs with no more than 45 nucleotide mutations, an expected barcode length of 15, no more than 10 nucleotide mutations in the 5’ region, no more than 4 nucleotide mutations in the 3’ region and no more than 1 nucleotide mutation in the spacer region were retained. We then generated a codon variant table from these processed CCSs using a minimally adapted version of a computational notebook created by Starr *et al*.^34^ (available on GitHub at https://github.com/jbloomlab/SARS-CoV-2-RBD_DMS/blob/master/results/summary/build_variants.md). A filter was first applied to ensure that only those CCSs with ccs-reported accuracies of 99.9% or greater in both the IgG1216-447 and barcode regions were retained. A filter (details at https://jbloomlab.github.io/alignparse/alignparse.consensus.html#alignparse.consensus.empirical_accuracy) was then applied to select for cases where there was concordance between the IgG1216-447 gene sequence called by CCSs with the same barcode. If indels (which are filtered out in a subsequent step anyway) are ignored, the empirical accuracy of the total region of the CCS covering the IgG1216-447 sequence in the library was 98.0%. In our library, on average, the barcodes were covered by 2.15 CCSs. Therefore, if the CCSs only differed marginally, then a consensus sequence was built, and if they differed widely, then the barcode was discarded, according to the criteria implemented in https://jbloomlab.github.io/alignparse/alignparse.consensus.html#alignparse.consensus.simple_mutconsensus. Any variants with indels in IgG1216-447 were then discarded. The codon variant lookup table (which links barcodes to IgG1216-447 mutations) was then generated using the dms_variants algorithm (https://jbloomlab.github.io/dms_variants/, version 1.4.3). The resulting lookup table contained 4,359 (98.9%) of the 4,408 total possible single-site amino acid substitutions, with only two positions (242 and 431) largely missing where the input library had failed to generate mutants.

Our own version of the process_ccs.ipynb and build_variants.ipynb notebooks are available on our GitHub at https://github.com/Ortlund-Laboratory/DMS_IgG1Fc/blob/main/PacBio_Library_Sequencing/Lib01/process_ccs.ipynb and https://github.com/Ortlund-Laboratory/DMS_IgG1Fc/blob/main/example_enrichment_escape/build_variants.ipynb, respectively.

#### Fluorescence-activated cell sorting of library to select mutants that cause tighter/weaker FcγR binding

Fc mutants that resulted in either tighter or weaker binding to Fcγ receptors were identified using the GFP^+^/Myc^+^ IgG1 Fc library cells. For each receptor, 30 million cells were washed with binding buffer (1X PBS plus 2% FBS and 10 mM HEPES, pH 7.5) and incubated with the receptor at a concentration equivalent to the *K*D of the relevant FcγR-Fc interaction, in a total volume of 0.5 mL, on a rotating wheel at room temperature for one hour. Subsequently, the cells were washed three times with additional binding buffer and stained with rabbit anti-Myc antibody and PerCP-eFluor™ 710 streptavidin (see **key resources table**). Cell sorting was performed using a BD FACSMelody^TM^ cell sorter (*BD Biosciences*) or a BD FACS Aria II instrument (*BD Biosciences)* at the Emory Flow Cytometry Core. Using the positive anti-Myc signal as a measure of expression, gates were drawn above the diagonal to capture cells with the highest 15% of signal (tighter-binding mutations) or below the diagonal to capture cells with the lowest 15% of signal (weaker-binding mutations) on an anti-Myc vs receptor signal plot. Typically, for each receptor experiment, between 3x10^5^ and 1x10^6^ cells were collected in LV-MAX production medium for both the tighter- and weaker-binding populations. In addition to these sorted populations, each experiment collected a reference cell population (5x10^6^ of the complete library) for sequencing. These reference cells were washed once in binding buffer and prepared for next-generation sequencing.

#### Fluorescence-activated cell sorting titrations of library to determine effective *K*_A_s for each mutant against each FcγR

The *K*A of each Fc mutant for every FcγR was determined by titrating GFP+/Myc+ library cells with serially diluted FcγR, using a dilution factor of 3. This approach established a range of receptor concentrations spanning several orders of magnitude above and below the respective effective *K*As for the WT, in addition to a 0M FcγR sample. The concentrations used, and the number of cells collected in each bin, for each receptor are provided in our GitHub at https://github.com/Ortlund-Laboratory/DMS_IgG1Fc/blob/main/Deposited_Data/FACS/FACS_BD_Melody_Data_titration_DMS_data_GitHub.xlsx. For each receptor titer, 10 million cells were washed with binding buffer (1X PBS plus 2% FBS and 10 mM HEPES, pH 7.5) and incubated with 200 μL of receptor prepared to the concentration relevant for that titer. The cells were then washed three times with binding buffer and stained with PerCP-eFluor™ 710 streptavidin (see **key resource table**). The 0M FcγR control sample was treated identically to other samples but was not stained with PerCP-eFluor™ 710 streptavidin before sorting. Cell sorting was performed using a BD FACSMelody^TM^ cell sorter (*BD Biosciences*) or using a BD FACS Aria II instrument (*BD Biosciences)* at the Emory Flow Cytometry Core. For analysis, streptavidin-positive cells were gated and plotted in a histogram. The range gate tool was used to define four bins based on streptavidin positivity, representing Fc-FcγR binding strength: Bin 1 for the weakest binding and Bin 4 for the strongest. The sample with the highest [FcγR] concentration defined the width of Bin 4, while the 0M FcγR sample established the range for Bin 1. Both bins captured 85-95% of streptavidin-positive cells. Bins 2 and 3 were set to equal widths. Presort analysis was performed to calculate cell percentages in each bin, which were multiplied by 1x10^6^ to determine the required cell numbers per titer. Sorting specific cell quantities was challenging at both low and high [FcγR] concentrations, particularly bin 4 (strong binding) under low [FcγR] and bin 1 (weak binding) under high [FcγR]. Any bin containing less than 500 cells was not advanced to sequencing analysis due to insufficient material for downstream RNA purification. The collection medium used was DMEM/F-12 with 10% FBS.

#### High-throughput sequencing of sorted cell populations

The following applies both to cells sorted for tighter/weaker-binding populations, and for binned populations. We washed the cells once with 1X PBS and extracted RNA using the GeneJet RNA Purification Kit (*Thermofisher)*. cDNAs were then prepared using the High-Capacity cDNA Reverse Transcription Kit (*Thermofisher*) with a primer specifically designed to anneal immediately downstream of the barcode sequence (Fc_Lib_RT; see **key resources table**). Amplification of the barcodes using Platinum^TM^ SuperFi^TM^ DNA Polymerase (*Thermofisher*) with Fc_Lib_FW01 and Fc_Lib_RT primers with Nextera-compatible overhang sequences (see **key resources table**) was then carried out over 8 rounds of PCR, with this increasing up to 15 rounds if necessary. The QIAquick PCR Purification Kit (*Qiagen*) was then used to purify the amplicons. Dual-indexed barcodes were then appended to the purified amplicons with the Illumina NexteraXT DNA Library Preparation Kit using the library amplification protocol but omitting the tagmentation protocol. Validation was achieved by capillary electrophoresis on an Agilent 4200 TapeStation, before pooling of samples and sequencing on an Illumina NovaSeq 6000 at PE100 or PE26x91 to achieve an approximate read-depth of 10 million reads per tighter- or weaker-binding sample, 50 million reads per reference sample, or a maximum of 5 million reads per binned sample.

#### Calculation of tighter/weaker binding scores from sequencing data analysis

The dms_variants package (see below) uses an *Illuminabarcodeparser* algorithm (https://jbloomlab.github.io/dms_variants/dms_variants.illuminabarcodeparser.html#dms_variants.illuminabarcodeparser.IlluminaBarcodeParser) which requires sequences to be of the form 5’-[R2_start]-upstream-barcode-downstream-[R1_start]-3’, where R1 is required, anneals downstream of the barcode and reads backwards, and R2 is optional, anneals upstream of the barcode and reads forwards. This setup was incompatible with our construct, which contains barcodes only at the R2 end. We wrote a custom python script (available on our GitHub at https://github.com/Ortlund-Laboratory/DMS_IgG1Fc/blob/main/example_enrichment_escape/R2_to_R1.py) to reverse complement our Illumina sequences for compatible parsing, and made sure to also reverse the read accuracies. All files were kept in zipped format to avoid memory blow-up.

Variant counting was then carried out using the computational notebook developed by Greaney *et al*.^50^ (available on GitHub at https://github.com/jbloomlab/SARS-CoV-2-RBD_MAP_Crowe_antibodies/blob/master/results/summary/count_variants.md) with minor modifications. With this notebook, we used the dms_variants package (https://jbloomlab.github.io/dms_variants/, version 1.4.3) to transform the Illumina sequences to counts for each barcode, and then used our lookup table (see **PacBio library sequencing and analysis**) to map these barcodes to IgG1216-447 mutations.

This mapping process generated a variant_counts.csv file which we manipulated (see details at our GitHub at https://github.com/Ortlund-Laboratory/DMS_IgG1Fc/tree/main/example_enrichment_escape/scores_and_visualization) to give three-column files (detailing the barcode identified, the mutation it maps to, and the number of times it was counted in the population) for the tighter-/weaker-binding and reference populations, respectively. These files provide the input for our custom script, BarcodeMapping.R (adapted from our previous study,^33^ and also available on our GitHub). In this script, we group our barcode counts cumulatively according to the mutations they are associated with, and then convert into tighter or weaker binding scores. Following grouping, the score for weaker binding for an individual mutation, *Ei* was defined as:

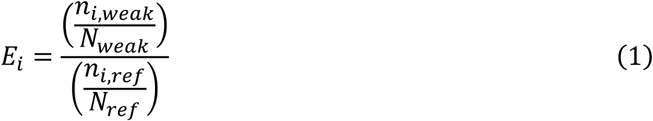

where *ni,weak* and *ni,ref* are the number of times mutation *i* is counted in the weaker binding and reference populations, respectively. *Nweak* is the sum of all mutation counts in the weaker binding population, Σi*ni,weak*, and *Nref* the sum of all mutation counts in the reference population, Σi*ni,ref*. These terms are included to normalize for the total fraction of the library that weakens receptor binding. To elude artificially high weak-binding scores caused by division with potential small numbers, the lowest 5% of abundance values in the reference population were set to the 5th-percentile value. To remove any further potential outliers, the top 1% of scores were set to the 99th-percentile value. To determine tighter-binding scores, the same equation is used, with the tighter-binding population used in place of the weaker binding population.

We then applied a number of transformations (also encoded in our BarcodeMapping.R file on our GitHub) to aid comparison of results between experiments. First, we normalized all scores to values between 0 and 1. We then applied an arcsine square root transformation to have our distribution follow Gaussian characteristics more closely, in line with our previous methods.^33^ In order to achieve a final distribution with mean 0 and standard deviation 1, we then performed Z-normalization on our data. We applied Fisher’s exact test to our data to determine adjusted p-values for each mutation, and which we quote with our tighter/weaker-binding scores.

### Calculation of effective *K*_A_s from sequencing data analysis

Variant counting followed the same process as for **Calculation of tighter/weaker binding scores from sequencing data analysis**. The position of the barcodes likewise necessitated the use of the custom script, R2_to_R1.py, and our lookup table (see **PacBio library sequencing and analysis**) was also used to map the identified barcodes to IgG1216-447 mutations. Furthermore, the dms_variants package (https://jbloomlab.github.io/dms_variants/, version 1.4.3) used to transform the Illumina sequences to counts for each barcode does not recognize bins with nil populations, but requires a consistent number of bins per titer. Occasionally (and particularly for low [FcγR], strong binding and high [FcγR], weak binding binning combinations), we found that the population does not produce sufficient cDNA from RNA extraction and reverse transcription for sequencing and therefore when it comes to barcode counting, those populations are empty. To resolve this, we wrote ‘placeholder’ files with just one sequence. These sequences consisted entirely of guanines, so chosen because this, plus its reverse complement, do not match any sequences in our library, and so ultimately will be filtered out. An example placeholder file is provided in our GitHub at https://github.com/Ortlund-Laboratory/DMS_IgG1Fc/tree/main/example_titration.

To construct titration curves and derive *K*D values from our counted barcodes, we used an adapted version of an R markdown script introduced by Starr *et al*.^34^ (the original computational notebook is provided on GitHub at https://github.com/jbloomlab/SARS-CoV-2-RBD_DMS/blob/master/results/summary/compute_binding_Kd.md, and our adapted version at https://github.com/Ortlund-Laboratory/DMS_IgG1Fc/blob/main/example_titration/FcgR2b_compute_binding_Kd.Rmd). At each receptor concentration, we calculated the mean bin response variable as a simple mean value, weighted according to the bin integer:

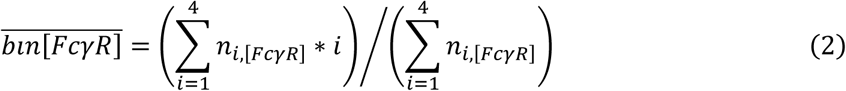

where *ni,[FcγR]* is the number of cells in bin *i* at a given FcγR concentration, with *i* ranging from 1 (corresponding to cells with weakest Fc-FcγR binding) to 4 (cells with strongest binding). We then used nonlinear least-squares regression to fit curves to the titration series for each barcode. This regression followed a standard non-cooperative Hill equation which related the mean bin response to the receptor concentration:

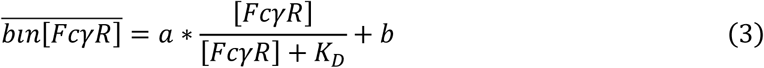

where *a* and *b* are free parameters for the titration response range (starting value set to 3) and the titration curve baseline (starting value set to 1), respectively. The starting *K*D value was always set to the sixth value in the 10-point concentration series. To aid fitting titration curves with aberrant data points, minimum and maximum bounds were applied for parameter *a* (1.5 and 3) and for parameter *b* (1 and 1.5). Our calculated *K*D values were also constrained to remain within the concentration range of our titration. To ensure our analyses were not affected by barcodes with few hits (which are more prone to statistical fluctuations) we applied a filter to remove barcodes where the average cell count was <5 across the concentration range, or if the cell count was <2 for 4 or more of the samples. Our final filter was applied to remove the top 5% of curves with the highest normalized mean square residual, where the normalization of the residual to between 0 and 1 had been achieved with parameter *a*. We quote effective association constant, *K*A, values in addition to *K*D values, converting using the relation log*K*A= -log*K*D.

To convert from per-barcode *K*D values to per-mutation *K*D values, we first removed all non-fitted barcodes, barcodes corresponding to wild type and barcodes corresponding to more than one nonsynonymous mutation from binding_Kds.csv, the output file from compute_binding_Kd.Rmd described above. We then used a further R script, (provided at https://github.com/Ortlund-Laboratory/DMS_IgG1Fc/blob/main/example_titration/results/binding_Kds/FcgR2b_combining_barcodes.R) to group barcodes, weighted by their average count from all concentrations, to their corresponding mutations.

#### Data visualization of tighter/weaker binding scores

Heatmaps were generated using custom scripts provided in our GitHub at https://github.com/Ortlund-Laboratory/DMS_IgG1Fc/tree/main/example_enrichment_escape/scores_and_visualization. For each heatmap, the *x-*axis corresponds to the IgG1 site, and ranges from 216 to 447, and the *y*-axis corresponds to amino acid type, organized into polar, nonpolar, aromatic, positive and negative groups. The wild type sequence for IgG1216-447 was defined by Fc_prot.fasta, also found in our GitHub example, and is represented by black dots on the heatmap. Tighter/weaker binding scores are represented by dots which deepen in hue as the score increases, and in size in proportion with the p-value. A bar chart which averages tighter/weaker binding scores per site is also produced.

#### Data visualization of effective *K*_A_s

Custom scripts, provided in our GitHub at https://github.com/Ortlund-Laboratory/DMS_IgG1Fc/tree/main/example_titration/heatmap, are used to generate heatmaps of similar format to those described in **Data Visualization of tighter/weaker binding scores**. Color-coding scales according to log*K*A values, centered on the wild type log*K*A Fc-FcγR interaction.

#### Comparative analysis of IgG1 Fc binding against different FcγRs

Custom scripts, provided in our GitHub at https://github.com/Ortlund-Laboratory/DMS_IgG1Fc/tree/main/example_differential_analysis, are used to generate tables of differential log*K*A values and associated heatmaps. The differential log*K*A values are generated by simple subtraction, and the heatmap color-coding is centered on the difference between the two wild type log*K*A Fc-FcγR interactions.

For a meta-analysis, differential log*K*A values were calculated for the FcγR2b (inhibitory) interaction versus the four studied lower-affinity activating receptor (FcγR2a-131H, FcγR2a-131R, FcγR3a-158F and FcγR3a-158V) interactions. These log*K*A values are therefore I:A ratios for FcγR2b binding versus the four studied activating receptors. To identify the most and least broadly inhibitory mutations, the results from the four differential analyses were averaged for each mutation. Mutations which have a log*K*A value differing by a factor greater than 2 away from the average for any of the individual differential analyses were then filtered out, in order to avoid selecting mutations which do not have consistent I:A ratios between the different activating receptors. Candidates at site 297 were also avoided since this is a glycosylation site, and will be the subject of future work. The top performing candidates were then selected for recombinant expression. A spreadsheet detailing this selection process is provided in our GitHub at https://github.com/Ortlund-Laboratory/DMS_IgG1Fc/tree/main/Deposited_Data/Differential_Titration_Data/Collected_Differential_Titration_Data.

#### Recombinant Fc and FcγR protein expression and purification

Proteins were expressed in Expi293 cells. The plasmids were transfected into Expi293 cells according to the manufacturer’s protocol (MAN0007814, Thermo Fisher Scientific) with the addition of penicillin/streptomycin mix 24 hours after transfection. The cells were cultured for 96 hours before harvesting.

A codon-optimized IgG1 Fc (residues 216-447) of rituximab was cloned into pcDNA3.4-TOPO vector (Azzam et al., 2024). Individual mutations were introduced using a Q5 site-directed mutagenesis kit. All sequences were confirmed by Sanger sequencing (Genewiz). IgG1 Fcs were purified using protein A resin (Thermo Fisher Scientific), with phosphate-buffered saline (PBS) (pH 7.4) being used as the binding and wash buffer and 100 mM sodium citrate buffer (pH 3.0) as the elution buffer. Eluted fractions were neutralized with 1 M Tris-HCl (pH 9.0), buffer-exchanged into PBS and stored at 4°C until ready for use.

Human FcγRIIIa (Uniprot P08637, 158V and 158F variants, residues 17-175), FcγRIIa (Uniprot P12318, 131R and 131H variants, residues 34-203), FcγRIIb (Uniprot 31994, residues 43-215), FcγRI (Uniprot P12314, residues 16-186) were cloned into the pTWIST CMV vector (TWIST Bioscience) with inclusion of a C-terminal biotin-attachment peptide (GLNDIFEAQKIEWHE) and N-terminal His6-tag. The proteins were purified using a HisTrap excel column (Cytiva) with PBS (pH 7.4) being used as the binding buffer and 500 mM imidazole, 150mM NaCl, 50 mM Tris-HCl pH 7.4 as the elution buffer. Eluted proteins were dialyzed against 50mM bicine pH 8.3, and biotinylated with BirA biotin ligase kit (Avidity) according to manufacturer instructions. The proteins were then buffer-exchanged into PBS, snap-frozen in liquid nitrogen and stored at -80°C for later use.

#### Thermal stability analysis

Thermal stability of the IgG Fc mutants was analyzed using a label-free method on a Prometheus Panta instrument (NanoTemper Technologies, Inc.). Each of the Prometheus standard glass capillaries were filled with 10 μL of the sample in PBS buffer at the protein concentration of 5 μM and loaded onto the instrument. Tryptophan emission was recorded at two wavelengths, 330 nm (emission maximum of the native state) and 350 nm (emission maximum of the unfolded state). A temperature range between 25 °C and 95 °C was tested with the temperature increasing at the rate of 2 °C per minute. The first derivative of the 350/330 nm ratio was plotted against the temperature yielding an unfolding profile curve, and the inflection points were calculated automatically using the PR.Panta Analysis 1.7.1 software.

#### Surface plasmon resonance (SPR) analysis

Surface plasmon resonance (SPR) assays were performed on an Octet SF3 SPR system (Sartorius) with 10 mM HEPES, pH 7.4, 150 mM NaCl, 0.05% Tween-20 as a running buffer, at a temperature of 25°C. Approximately 1000 RU of recombinant protein A (Pierce) were immobilized by standard amine coupling with EDC/NHS (Amine Coupling Kit, Cytiva) to all 3 channels of a CDH SPR sensor chip (Sartorius). Two different Fc mutants (ligands) were captured onto the surface of channel 1 and channel 3 directly from the protein expression cell culture supernatants to the level of approximately 150 RU, while channel 2 was used as a reference. Subsequently, each FcγR (analyte) was injected in a series of six 3-fold dilutions, starting from 5 µM, with two blank cycles performed per each pair of ligands captured. After each cycle, the chip surface was regenerated with 20 mM HCl, and a fresh portion of the ligand was captured. Resulting sensorgrams were double-referenced and analyzed by fitting to a 1:1 steady-state affinity model.

#### Quantification and statistical analysis

Deep mutational sequencing analyses were performed using a mixture of adapted^33,34,50^ and custom code, which is available on GitHub at https://github.com/Ortlund-Laboratory/DMS_IgG1Fc.

## Supplemental Information

**Figure S1.**
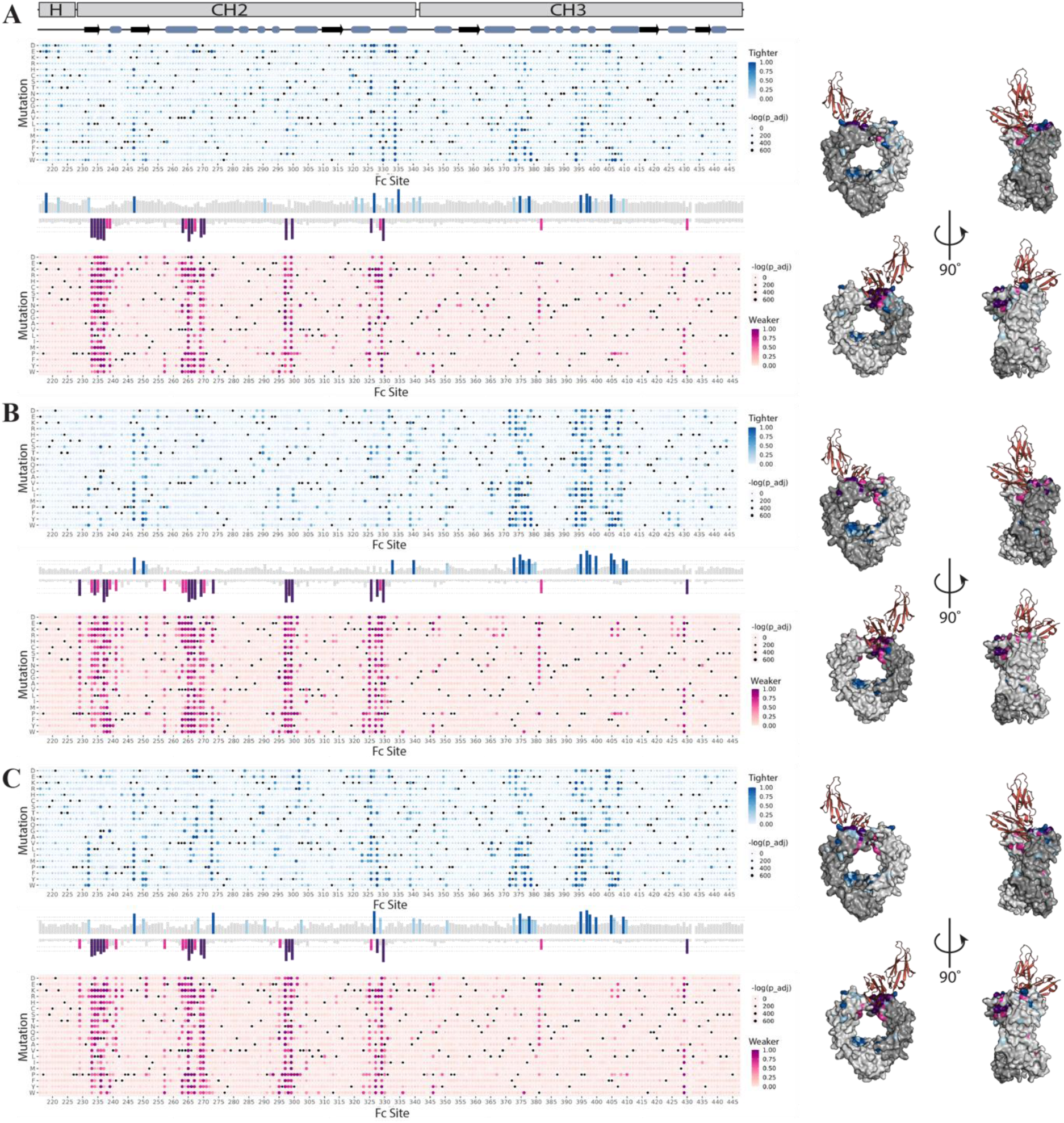
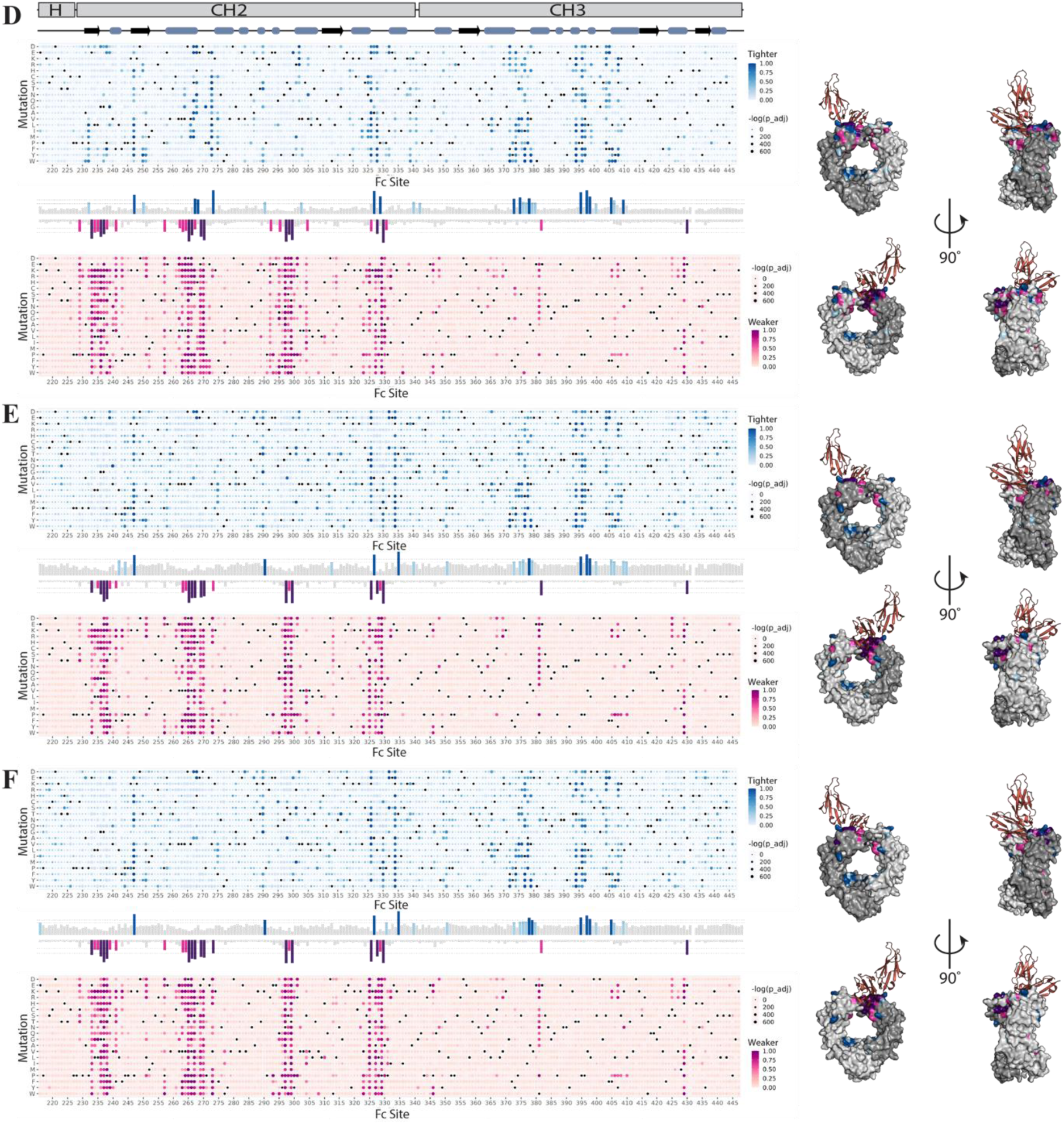
A-C. Single concentration tighter/weaker binding scores for all FcγRs. Heatmaps (*left*) give the tighter (blue) and weaker (magenta) binding scores for all single-point variants of an individual receptor. Average scores per site are given in bar charts with each heatmap. Per-site average scores exceeding one (light blue/pink) or two (dark blue/magenta) standard deviations above the mean are highlighted on Fc dimer structures (*right*). Receptors depicted are FcγR1 (A), FcγR2a-131H (B) and FcγR2a-131R (C). **D-F. Single concentration tighter/weaker binding scores for all FcγRs.** Heatmaps (*left*) give the tighter (blue) and weaker (magenta) binding scores for all single-point variants of an individual receptor. Average scores per site are given in bar charts with each heatmap. Per-site average scores exceeding one (light blue/pink) or two (dark blue/magenta) standard deviations above the mean are highlighted on Fc dimer structures (*right*). Receptors depicted are FcγR2b (D), FcγR3a-158F (E) and FcγR3a-158V (F).

**Figure S2.**
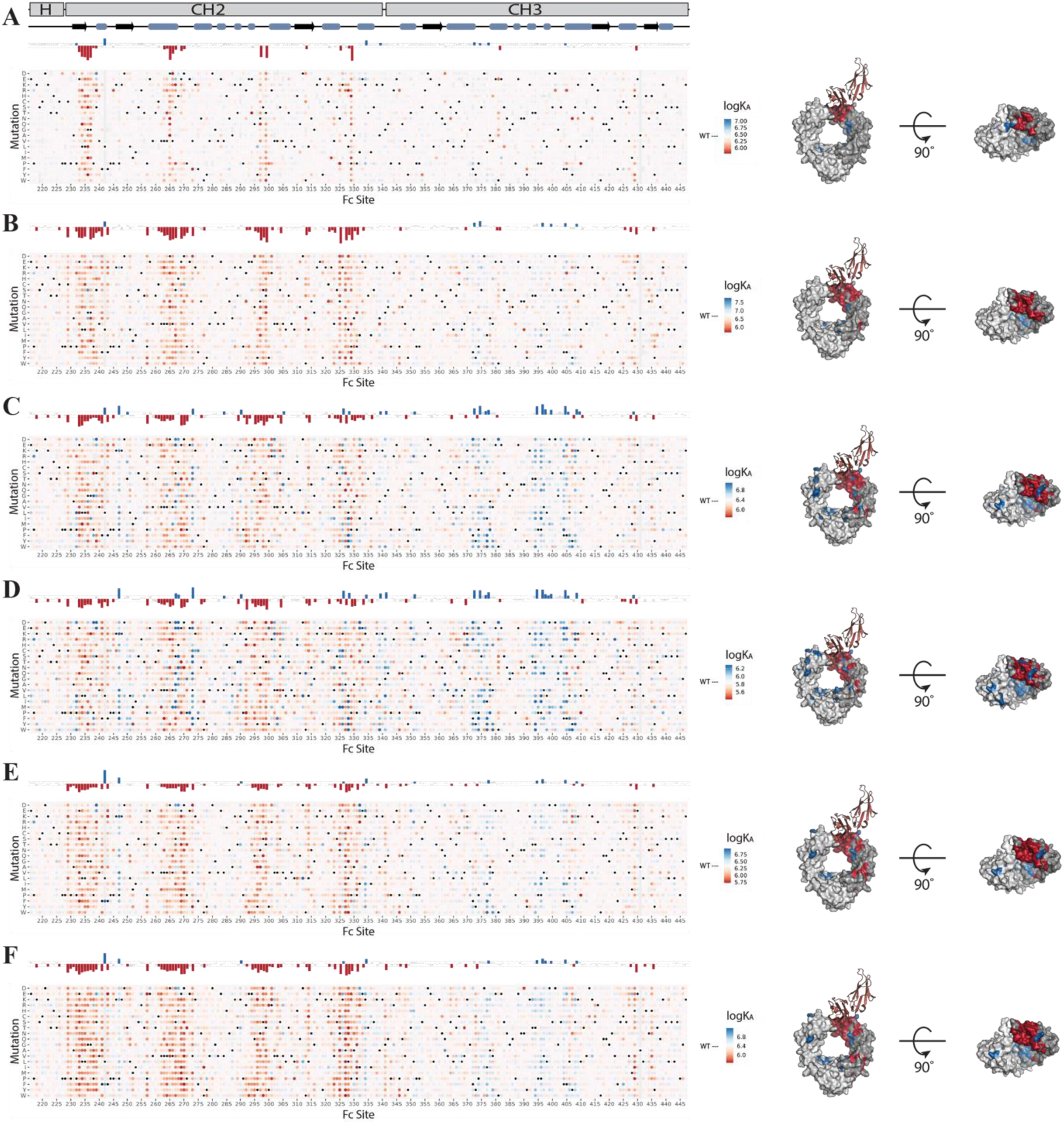
Titration data for all FcγRs. Heatmaps (*left*) give the log*K*A values for all single-point variants of an individual receptor. The color scale is normalized to the wildtype log*K*A for the particular receptor, with positive scores (blue) indicating enhanced binding with respect to the wildtype, and negative scores (red) indicating reduced binding. Average scores per site are given in bar charts above each heatmap. Per-site average scores exceeding 0.75 standard deviations above (blue) or below (red) the mean are highlighted on Fc dimer structures (*right*). Receptors depicted are FcγR1 (A), FcγR2a-131H (B), FcγR2a-131R (C), FcγR2b (D), FcγR3a-158F (E) and FcγR3a-158V (F).

**Figure S3.**
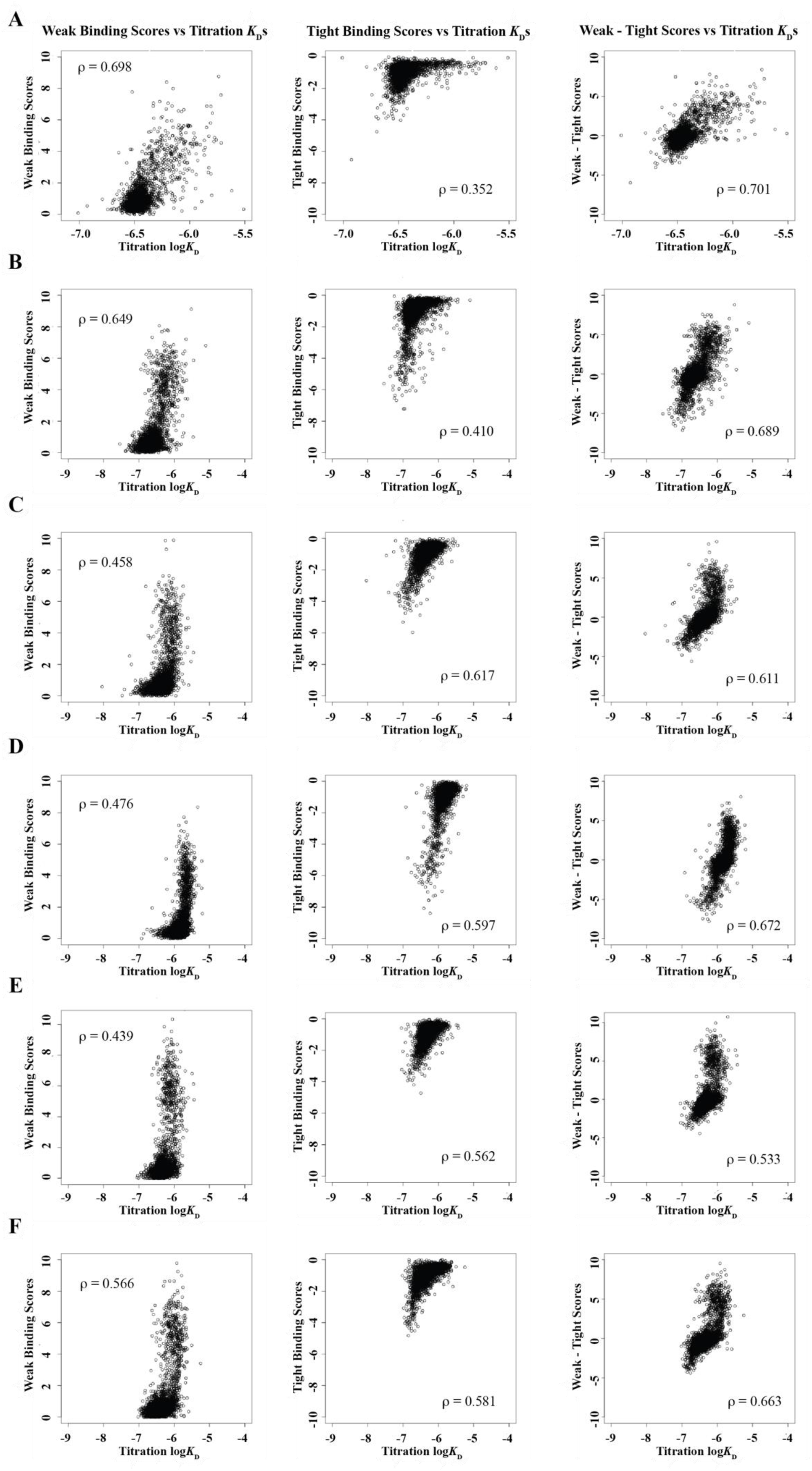
Correlation measurements between the tighter/weaker binding and titration approaches. Scatterplots give the correlations between variant weaker binding population scores and titration log*K*Ds (*left*), variant tighter binding population scores and titration log*K*Ds (*center*), and variant calculated weaker minus tighter binding values and titration log*K*Ds (*right*) for experiments with FcγR1 (A), FcγR2a-131H (B), FcγR2a-131R (C), FcγR2b (D), FcγR3a-158F (E) and FcγR3a-158V (F), respectively. A Pearson correlation coefficient, ρ, is provided for each comparison. Apart from FcγR1 tighter binding vs titration, these coefficients are all moderate-to-high, suggesting the tighter/weaker binding approach provides a moderately effective approximation of the variants most likely to enhance/diminish binding identified by the titration approach.

**Table S1.** Strongest- and weakest-binding IgG1 Fc variants for all receptors studied. Results are provided as log*K*As, where a larger value indicates stronger Fc-FcγR binding, and a lower value corresponds to weaker binding. Cells are color-coded according to how often the particular variant appears across the receptors studied: yellow corresponds to 4 instances; green to 3; blue to 2; and magenta to 1.

| | Rank | FcγR1 | | FcγR2a-131H | | FcγR2a-131R | | FcγR2b | | FcγR3a-158F | | FcγR3a-158V | |
| --- | --- | --- | --- | --- | --- | --- | --- | --- | --- | --- | --- | --- | --- |
| | | Variant | $\log K_A$ | Variant | $\log K_A$ | Variant | $\log K_A$ | Variant | $\log K_A$ | Variant | $\log K_A$ | Variant | $\log K_A$ |
| **Strongest:** | 1<sup>st</sup> | S239D | 6.928 | G236S | 7.538 | S267E | 8.040 | S239D | 6.923 | D312L | 7.048 | S267D | 6.987 |
|  | 2<sup>nd</sup> | K334P | 6.742 | G236A | 7.365 | L328W | 7.410 | L328W | 6.865 | I332D | 7.035 | I332D | 6.966 |
|  | 3<sup>rd</sup> | G371P | 6.714 | V379W | 7.288 | W381D | 7.264 | S267E | 6.765 | S239D | 7.007 | L242F | 6.950 |
|  | 4<sup>th</sup> | K334E | 6.703 | P396I | 7.285 | L328Y | 7.161 | L328Y | 6.666 | I377W | 6.967 | S239E | 6.945 |
|  | 5<sup>th</sup> | I332E | 6.698 | P374F | 7.248 | F372D | 7.151 | S267Q | 6.655 | L242F | 6.933 | D312L | 6.930 |
|  | 6<sup>th</sup> | S239E | 6.692 | G402Q | 7.244 | I377Y | 7.131 | H268E | 6.609 | I377F | 6.928 | C425F | 6.923 |
|  | 7<sup>th</sup> | I332D | 6.685 | F405E | 7.228 | L328F | 7.120 | P232L | 6.574 | P247V | 6.878 | K334E | 6.917 |
|  | 8<sup>th</sup> | P217R | 6.679 | P374L | 7.217 | P396I | 7.119 | V273F | 6.550 | K334E | 6.869 | T393K | 6.909 |
|  | 9<sup>th</sup> | C425F | 6.660 | P217R | 7.195 | P232L | 7.099 | A378P | 6.541 | P396I | 6.851 | K326E | 6.908 |
|  | 10<sup>th</sup> | D280I | 6.658 | K370H | 7.185 | S267A | 7.098 | I377W | 6.489 | H268D | 6.849 | V379Y | 6.897 |
| **Wildtype:** | N/A | . | 6.455 | . | 6.727 | . | 6.365 | . | 5.832 | . | 6.353 | . | 6.397 |
|  | 10<sup>th</sup> | Y407N | 5.807 | N297I | 5.727 | G236L | 5.739 | T299D | 5.425 | V273Y | 5.727 | D270F | 5.661 |
|  | 9<sup>th</sup> | G236L | 5.799 | S267I | 5.714 | A327S | 5.724 | G236Y | 5.425 | L328H | 5.726 | N325A | 5.659 |
|  | 8<sup>th</sup> | V266D | 5.793 | T299Q | 5.696 | A327Q | 5.688 | R292H | 5.420 | S298Q | 5.725 | W277I | 5.658 |
|  | 7<sup>th</sup> | N325P | 5.778 | G237K | 5.692 | E233L | 5.662 | V263F | 5.415 | N297A | 5.703 | A327W | 5.656 |
|  | 6<sup>th</sup> | D265A | 5.773 | D265R | 5.680 | E269P | 5.660 | Q295H | 5.409 | F243K | 5.684 | E269I | 5.652 |
|  | 5<sup>th</sup> | L234Q | 5.757 | V263K | 5.663 | P331G | 5.609 | D270N | 5.381 | E233F | 5.676 | Y300R | 5.636 |
|  | 4<sup>th</sup> | D265N | 5.736 | D265Q | 5.646 | I332R | 5.574 | E269P | 5.371 | E233S | 5.644 | E269P | 5.635 |
|  | 3<sup>rd</sup> | L328R | 5.716 | Q295D | 5.619 | N297A | 5.537 | R292P | 5.365 | E269T | 5.601 | S298D | 5.632 |
|  | 2<sup>nd</sup> | G236M | 5.714 | N297A | 5.515 | L234M | 5.521 | N297A | 5.334 | I332R | 5.491 | D270V | 5.629 |
| **Weakest:** | 1<sup>st</sup> | E294T | 5.617 | P329Y | 5.104 | Q295D | 5.452 | G237K | 5.218 | L410P | 5.433 | A327Q | 5.501 |

**Figure S4.**
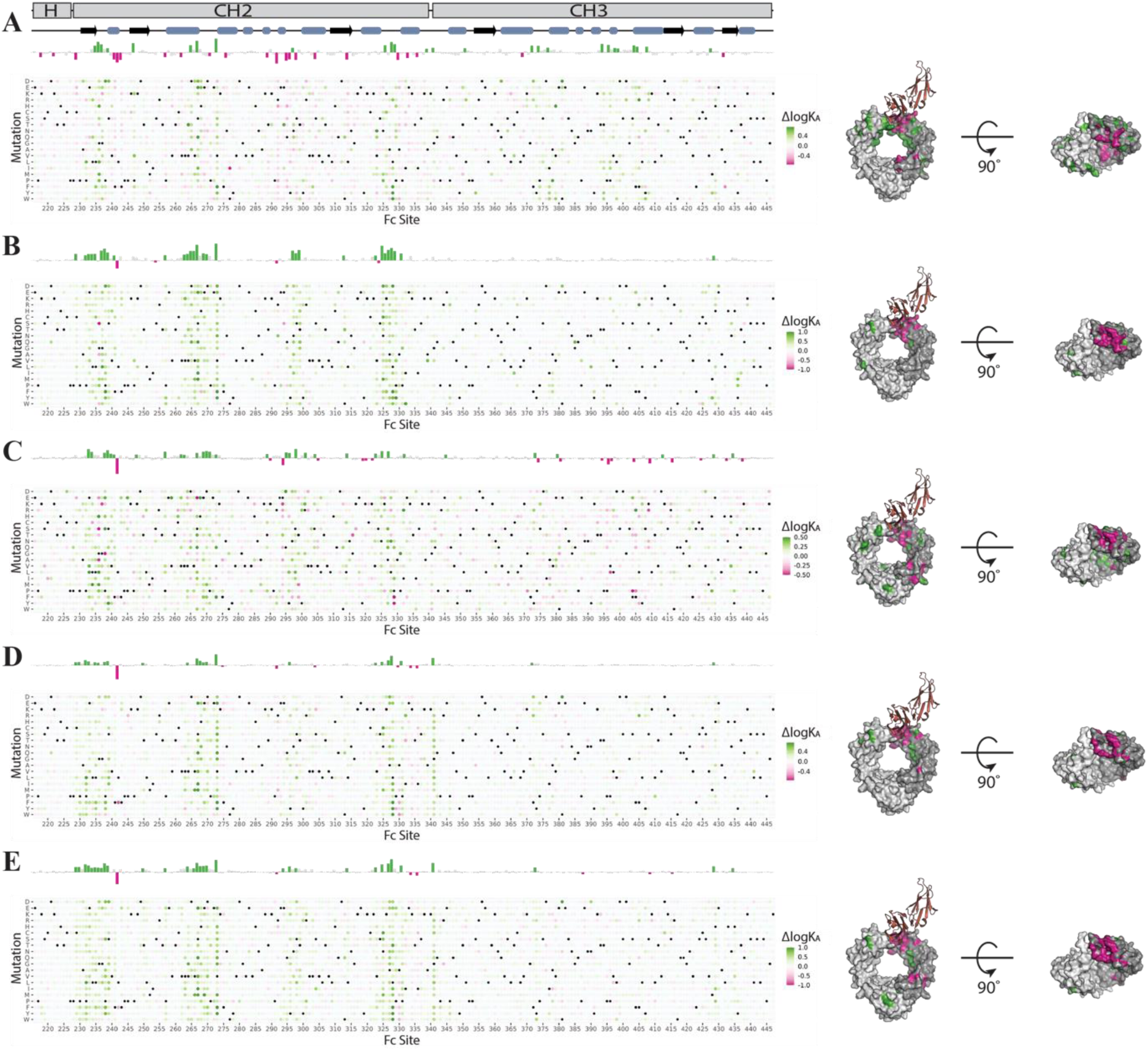
Differential titration data for FcγR2b vs all other FcγRs. Heatmaps (*left*) give the difference in log*K*A values for all single-point variants of FcγR2b vs a selected receptor. The color scale is normalized to the difference in wildtype log*K*A values for FcγR2b and the selected receptor, respectively. Positive scores (green) indicate stronger binding to FcγR2b, and negative scores (pink) indicate stronger binding to the other receptor. Average scores per site are given in bar charts above each heatmap. Per-site average scores exceeding 1.0 standard deviation above (green) or below (pink) the mean are highlighted on Fc dimer structures (*right*). Receptors depicted are FcγR2b minus: FcγR1 (A), FcγR2a-131H (B), FcγR2a-131R (C), FcγR3a-158F (D) and FcγR3a-158V (E).

**Figure S5.**
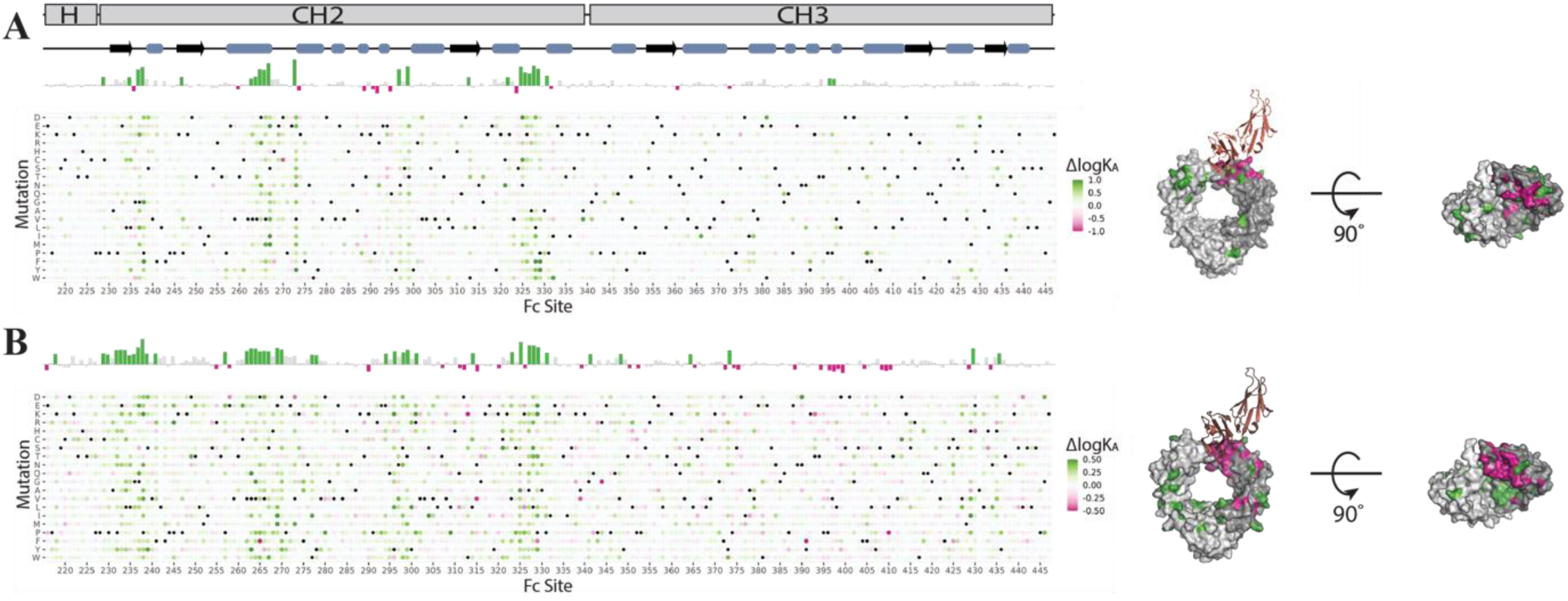
**Differential titration data for FcgR2a and FcgR3a polymorphs.** (A) FcgR2a-131R - FcgR2a-131H. (B) FcgR3a-158F - FcgR3a-158V. Heatmaps (*left*) give the difference in log*K*A values for all single-point variants of the two selected receptors. The color scale is normalized to the difference in wildtype log*K*A values for the two selected receptors. Positive scores (green) indicate stronger binding to the first receptor listed, and negative scores (pink) indicate stronger binding to the second receptor listed. Average scores per site are given in bar charts above each heatmap. Per-site average scores exceeding 1.0 standard deviations above (green) or below (pink) the mean are highlighted on Fc dimer structures (*right*).

**Table S2.** IgG1 Fc variants giving the highest and lowest I:A ratios, where 2b is compared against all activating receptors (2a-131H, 2a-131R, 3a-158F and 3a-158V) and just 2a-131R and 3a-158F. The latter case is a polymorphic combination associated with worse health outcomes for patients suffering from follicular lymphoma.^52^

|  | **2b vs All 2a and 3a** | **2b vs 2aR and 3aF** |
| --- | --- | --- |
| **Lowest I:A** | L242F | P329Y |
|  | D376A | L242F |
|  | S337G | G237K |
|  | Q438S | V397L |
|  | N389T | T299M |
|  | P245H | S354A |
|  | E388I | P245R |
|  | H285K | G371P |
|  | V302Y | K409N |
|  | G385Y | F275A |
| ... | ... | ... |
|  | S298L | S239I |
|  | P331G | A327N |
|  | A327N | S267T |
|  | L328E | S298L |
|  | S267T | H268E |
|  | H268E | I332R |
|  | V273Q | V273Q |
|  | F405R | F405R |
|  | V273F | V273F |
| **Highest I:A** | S267Q | S267Q |

**Figure S6.**
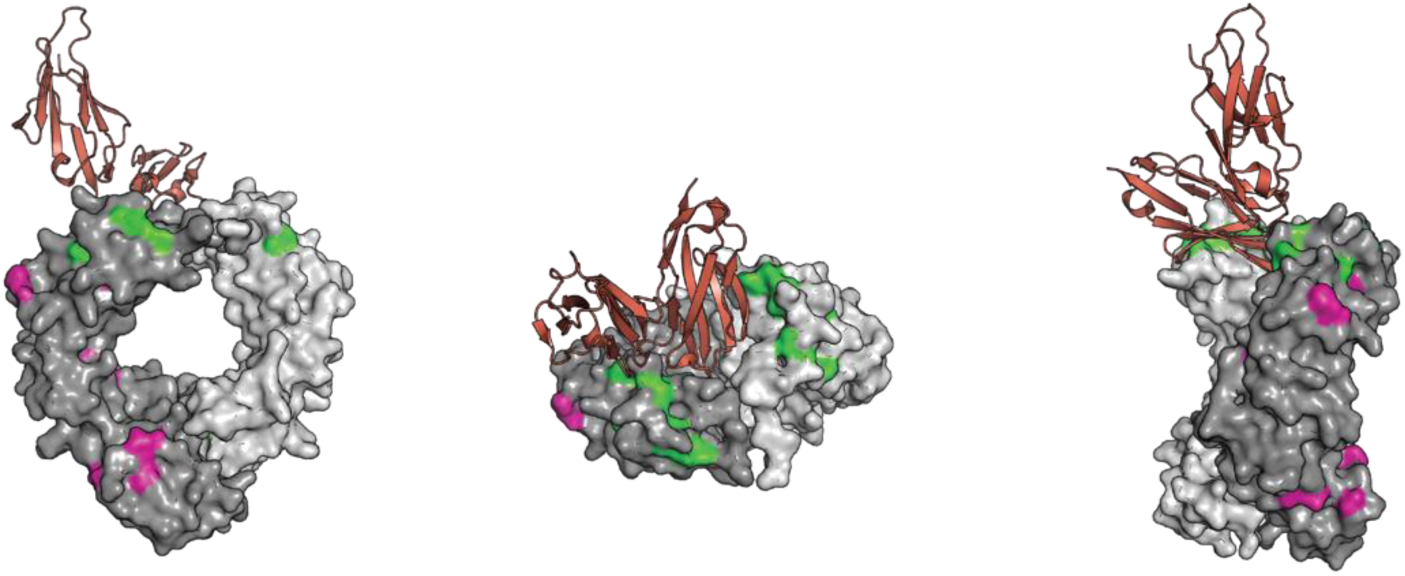
IgG Fc variants most strongly influencing the I:A ratio. The log*K*A values for Fc-FcγR2b binding were compared against the values generated by titrations with activating FcγRs, as described in the main text. IgG Fc variants which result in strongest binding to FcγR2b compared to the activating receptors (giving the highest I:A values) are color-coded in green, and those resulting in weakest binding (giving the lowest I:A values) are color-coded in pink.

**Table S3.**
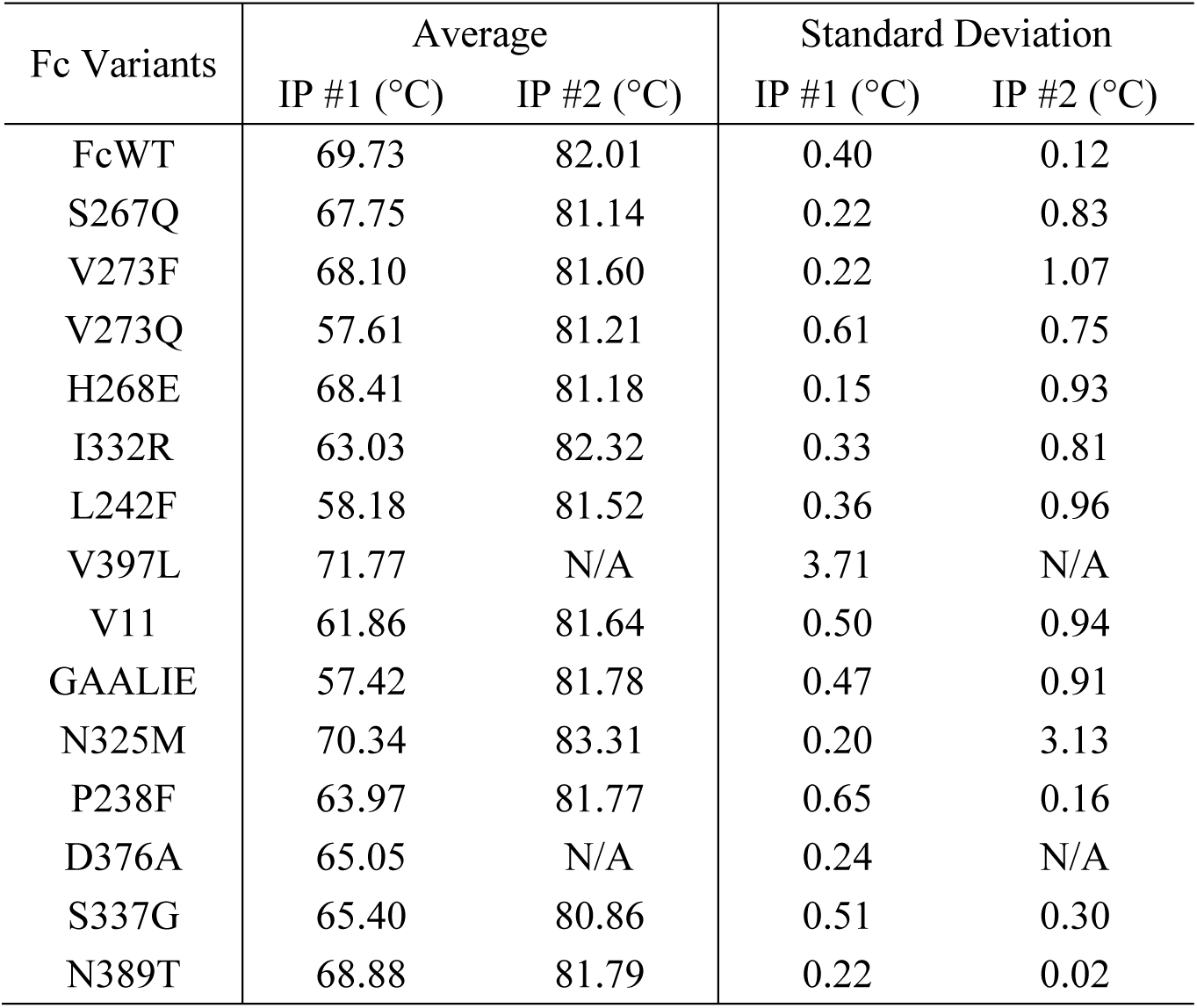
Thermostablities of selected IgG1 Fc mutants as determined by differential scanning fluorimetry.

**Figure S7.**
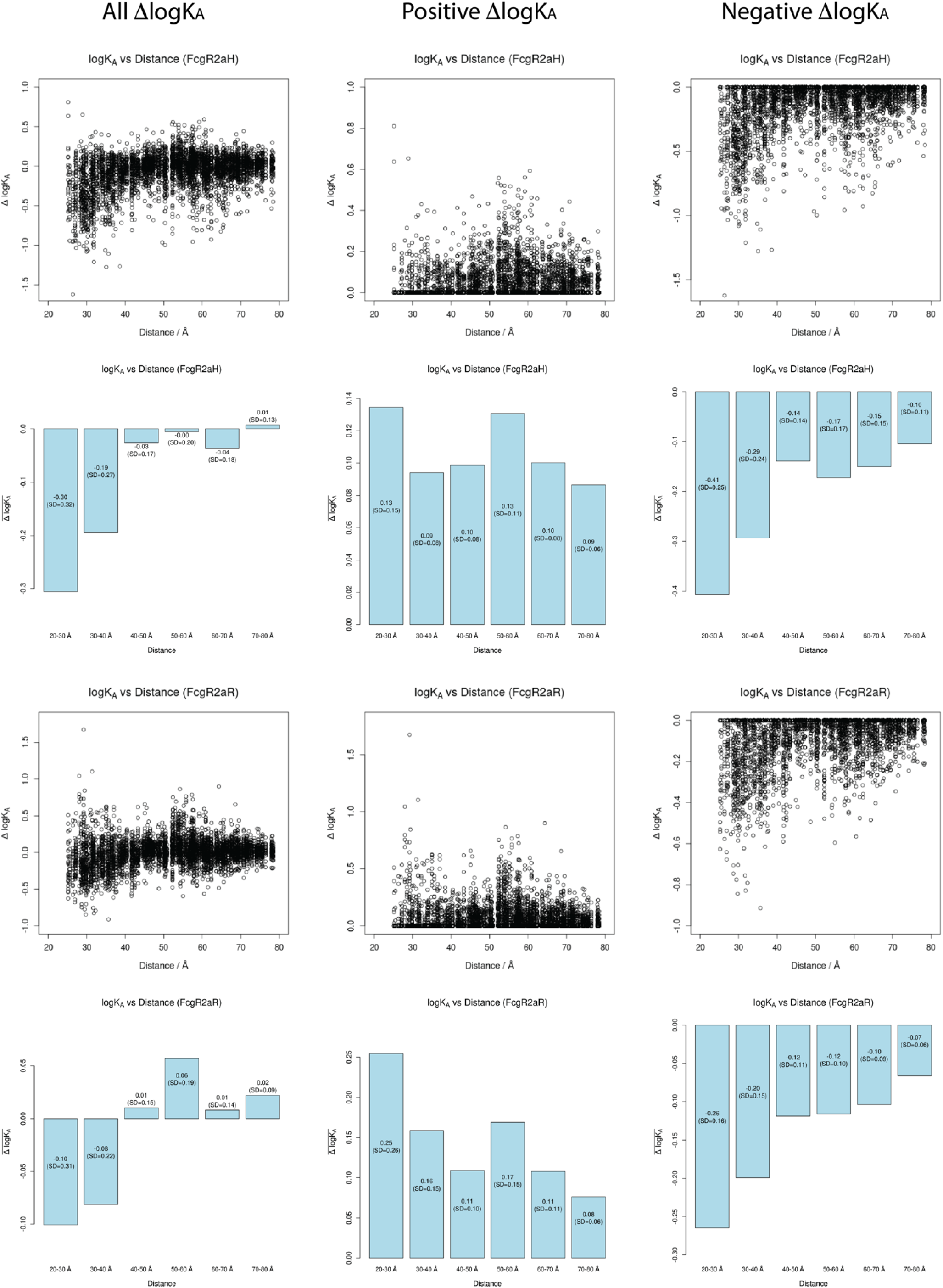
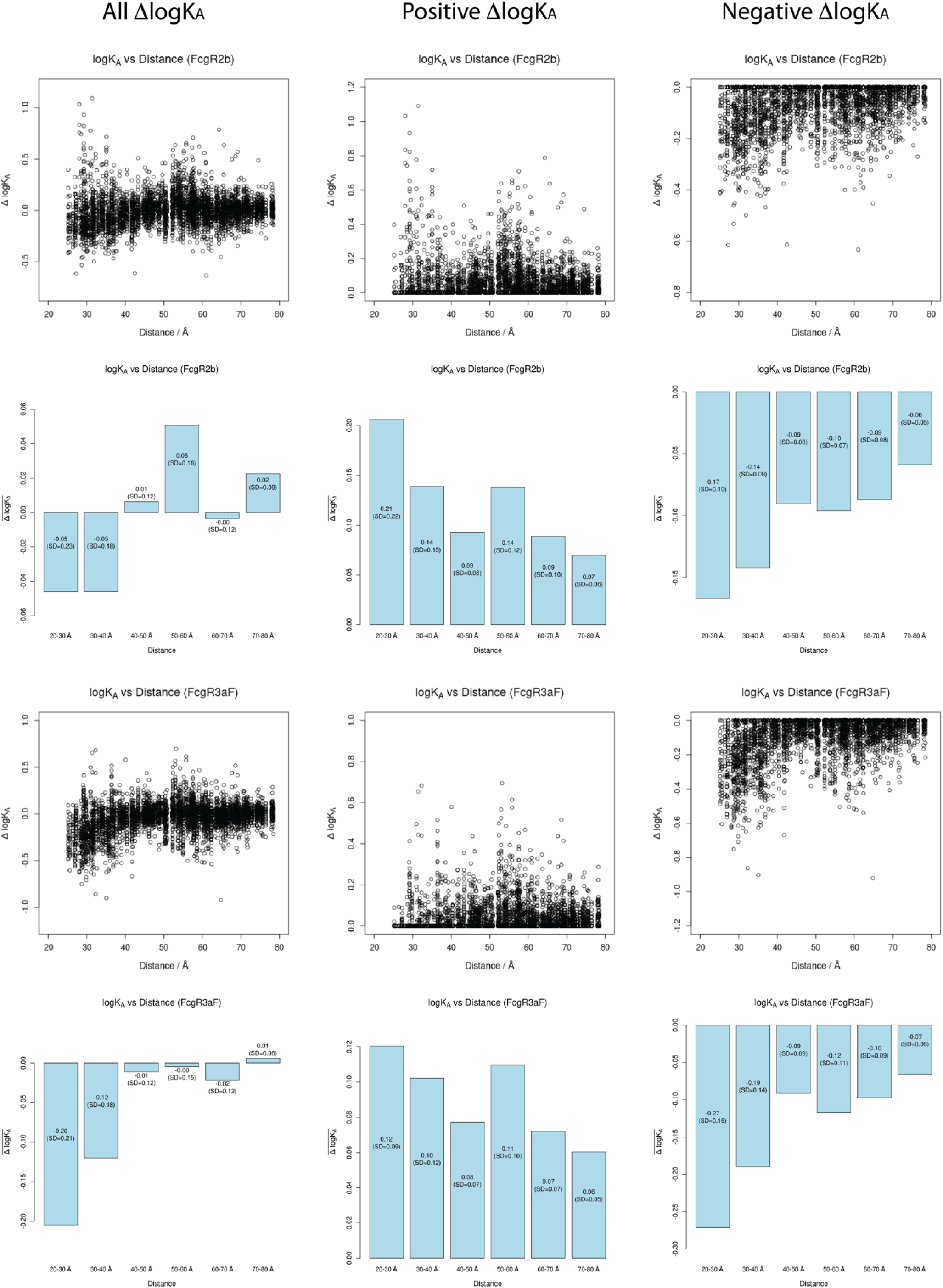
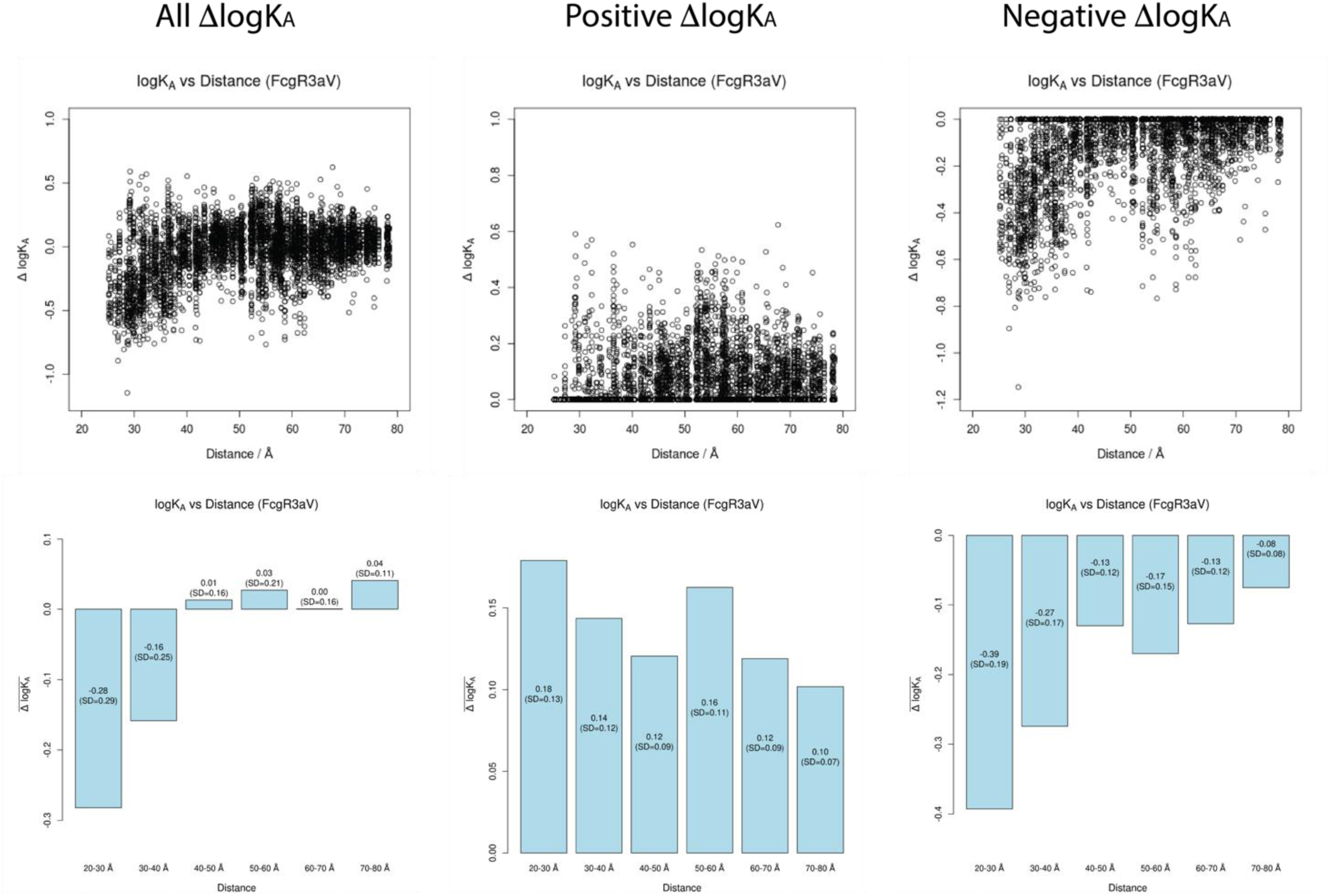
Quantification of ΔlogKA versus distance from Center of Mass (CoM) of receptor. Scatter plots of ΔlogKA (with respect to WT) were plotted against the distance between the CoM of the mutated residue on Fc against the CoM of the receptor, as measured using the only available crystal structure of a wild type Fc-FcγR complex, PDB code 3SGJ.^38^ The receptor for this structure is FcγR3aV, and was used as the basis for all CoM distance calculations, under the assumption that all receptors (except for FcγR1, which is known to be structured and behave differently)^43^ bind to the same position of Fc and are structurally the same. These assumptions were tested using Alphafold 3,^53^ which suggest they were justified (see Fig. S9). The scatter plot results were binned into six separate distances, and the ΔlogKA results within each bin averaged. Bar charts immediately underneath their respective scatter plots are plotted, showing the average ΔlogKA for each distance, within 10 Å increments. The mean value and standard deviation are plotted within each bar. In the left column, all data points are plotted; in the center column, only the positive data points are plotted; and in the right column, only the negative data points are plotted. This strategy was implemented to better identify trends. The standard deviations for each bar are typically large, and sometimes even exceed the mean values, which highlights the significant variability in the effects mutations have on changing the logKA. Standard deviations are typically highest within the 20-30 Å bin, showing that this variability is greatest the closer Fc is to the receptor. Also, standard deviations are typically lower for interactions with the FcγR1 receptor, suggesting less variation in the possible effects mutations have on Fc binding to this receptor. With all receptors, the average ΔlogKA is most negative within the 20-30 Å bin, suggesting mutations within the section of Fc most proximate to the receptor are, on average, most likely to decrease binding affinity between Fc and its receptor. Considering mutations which have a positive ΔlogKA only reveals that, for each receptor, binding affinities increase to the greatest extent, on average, within the 20-30 Å bin. These results show that, even though mutations to residues within this bin are most likely to lead to a decrease in binding affinity, there are also a significant number of possible mutations which could be applied that would markedly increase binding. Analysis of the positive ΔlogKA-only results also reveals that the average ΔlogKA for the 50-60 Å bin has the second-highest value for all of the strongly activating receptors, and is the third-highest bin (with the 30-40 Å bin coming second) for the mildly activating receptor, FcγR1, and the inhibitory receptor, FcγR2b. In the absence of allosteric effects, it would be expected that the effect of mutations on binding affinity would tail off with increased distance from the receptor, thus suggesting that some residues contained within the 50-60 Å bin are capable of exerting allosteric effects. Furthermore, since this bin does not constitute as significant an outlier when considering the negative ΔlogKA only, these results suggest that there is particularly significant scope within 50-60 Å bin for mutations which will increase binding affinity. Taken together, these data suggest that residues within the 20-30 Å bin will be the most promising targets for mutations if the goal is to either increase *or* decrease the binding affinity, due mainly to the direct contact between these residues and the receptor, and that residues within the 50-60 Å bin would be the second-most promising to target if the goal is to increase binding affinity, due to the allosteric effects these residues appear to exert on the Fc-FcγR binding interface.

**Figure S8.**
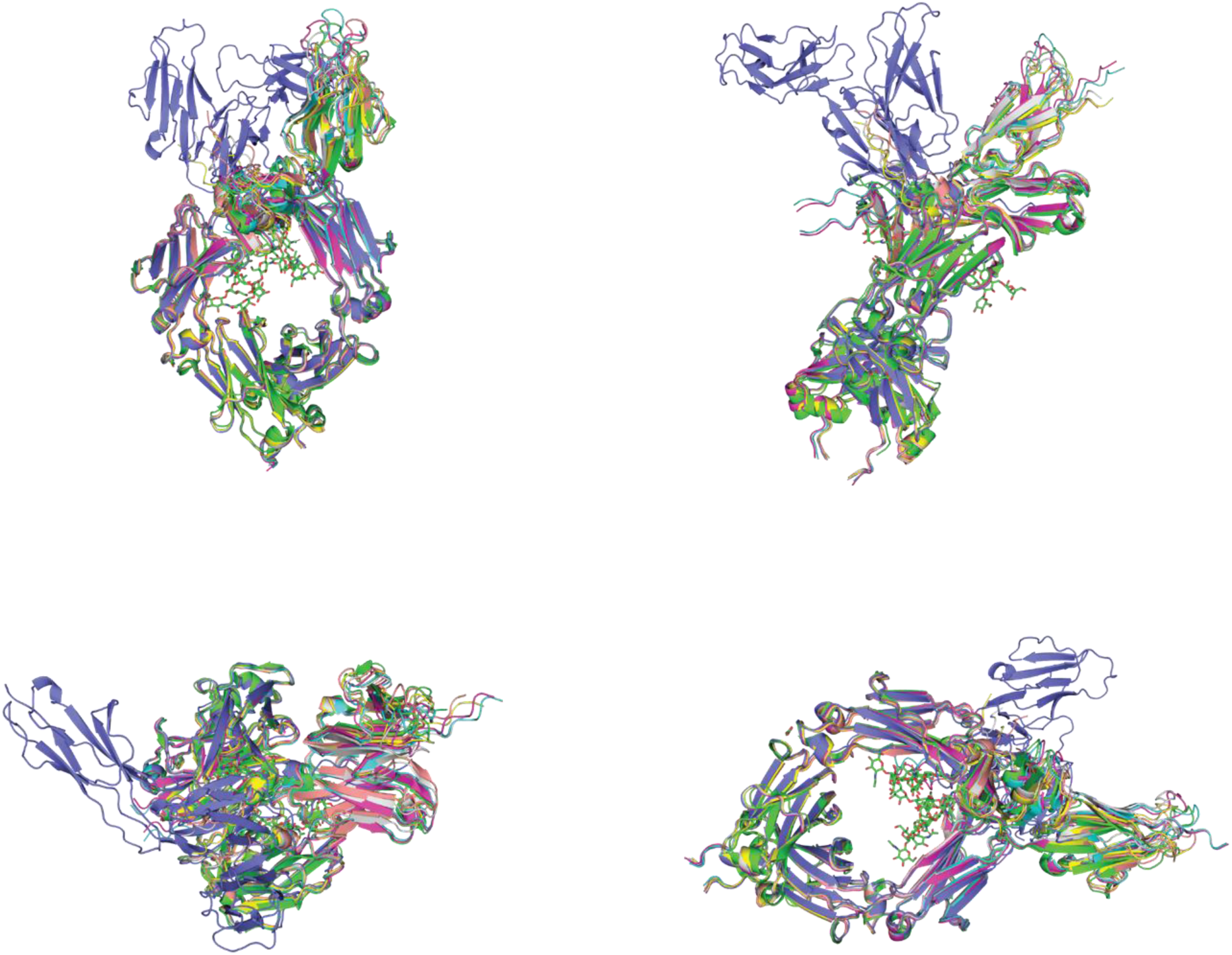
Representations of the overlaid structures of the Fc dimer bound to each receptor investigated in this study. The structure color-coded green represents the Fc dimer bound to FcγR3aV, determined from crystallography (PDB code 3SGJ).^38^ As a control, the sequence determined from this crystal structure was used as input for Alphafold 3,^53^ and the predicted structure mapped in cyan. To predict Fc-FcγR3aF binding, this same sequence was used as input to Alphafold 3, with the mutation V158F applied to the receptor; the predicted structure is mapped in magenta. To predict Fc-FcγR2b binding, the receptor sequence from PDB 3WJJ^41^ was used; the predicted structure is mapped in yellow. To predict Fc-FcγR2aR binding, the receptor sequence from PDB 3RY5^54^ was used; the predicted structure is mapped in salmon. To predict Fc-FcγR2aH binding, the receptor sequence from PDB 3RY4^54^ was used; the predicted structure is mapped in gray. To predict Fc-FcγR1 binding, the receptor sequence from PDB 3RJD^55^ was used; the predicted structure is mapped in blue. These alignments suggest that all receptors, other than FcγR1, fold to near-identical structures, and bind to the same sites of the Fc dimer, therefore justifying the approximations used to calculate receptor centers of mass.

**Table S4.** IgG1 Fc variants within the 20-30 Å and 50-60 Å layers which cause a positive ΔlogKA against the inhibitory FcγR2b receptor whilst simultaneously causing a negative ΔlogKA against all activating receptors (FcγR2a-131H, FcγR2a-131R, FcγR3a-158F and FcγR3a-158V), and *vice versa*. Quoted values are the ΔlogKA against FcγR2b minus the average ΔlogKA against all activating receptors. Therefore, a greater positive value denotes greater inhibition, and a greater negative value denotes greater activation. *= data for FcγR2a-131H and FcγR3a-158F are missing.

| **Mutation**<br>**20-30 Å Layer** | **$\Delta\log K_A$**<br>**20-30 Å Layer** | **Mutation**<br>**50-60 Å Layer** | **$\Delta\log K_A$**<br>**50-60 Å Layer** |
| --- | --- | --- | --- |
| P232K | +0.276 | K218F | +0.207 |
| E233T | +0.280 | K218L | +0.213 |
| L234W | +0.227 | K218N | +0.099 |
| G236L | +0.378 | H224N | +0.081 |
| G237E | +0.406 | K248D | +0.174 |
| G237L | +0.535 | K248P | +0.167 |
| G237R | +0.178 | K248W | +0.087 |
| P238A | +0.415 | T250L | +0.106 |
| P238I | +0.362 | T250Q | +0.107 |
| P238S | +0.929* | L251I | +0.164 |
| S267C | +0.277 | L251P | +0.201 |
| S267T | +0.550 | L251R | +0.243 |
| E269V | +0.273 | R255C | -0.099 |
| A327N | +0.522 | P257K | +0.267 |
| L328C | +0.220 | T307C | -0.126 |
| L328D | +0.550 | T307F | +0.108 |
| L328E | +0.537 | T307L | +0.104 |
| P329N | +0.267 | H310P | +0.055 |
|  |  | Q311A | -0.052 |
|  |  | W313T | +0.364 |
|  |  | L314I | +0.120 |
|  |  | G316H | +0.170 |
|  |  | K317V | -0.124 |
|  |  | G341Y | +0.315 |
|  |  | Q342F | -0.090 |
|  |  | Q342K | -0.106 |
|  |  | P343W | +0.155 |
|  |  | S375W | -0.082 |
|  |  | D376T | -0.191 |
|  |  | I377L | +0.068 |
|  |  | A378T | -0.108 |
|  |  | V379F | -0.154 |
|  |  | T393G | +0.110 |
|  |  | T393Y | +0.116 |
|  |  | P395V | -0.131 |
|  |  | V397S | +0.130 |
|  |  | F405R | +0.822 |
|  |  | E430F | +0.125 |
|  |  | E430Y | +0.251 |

**Table S5.**
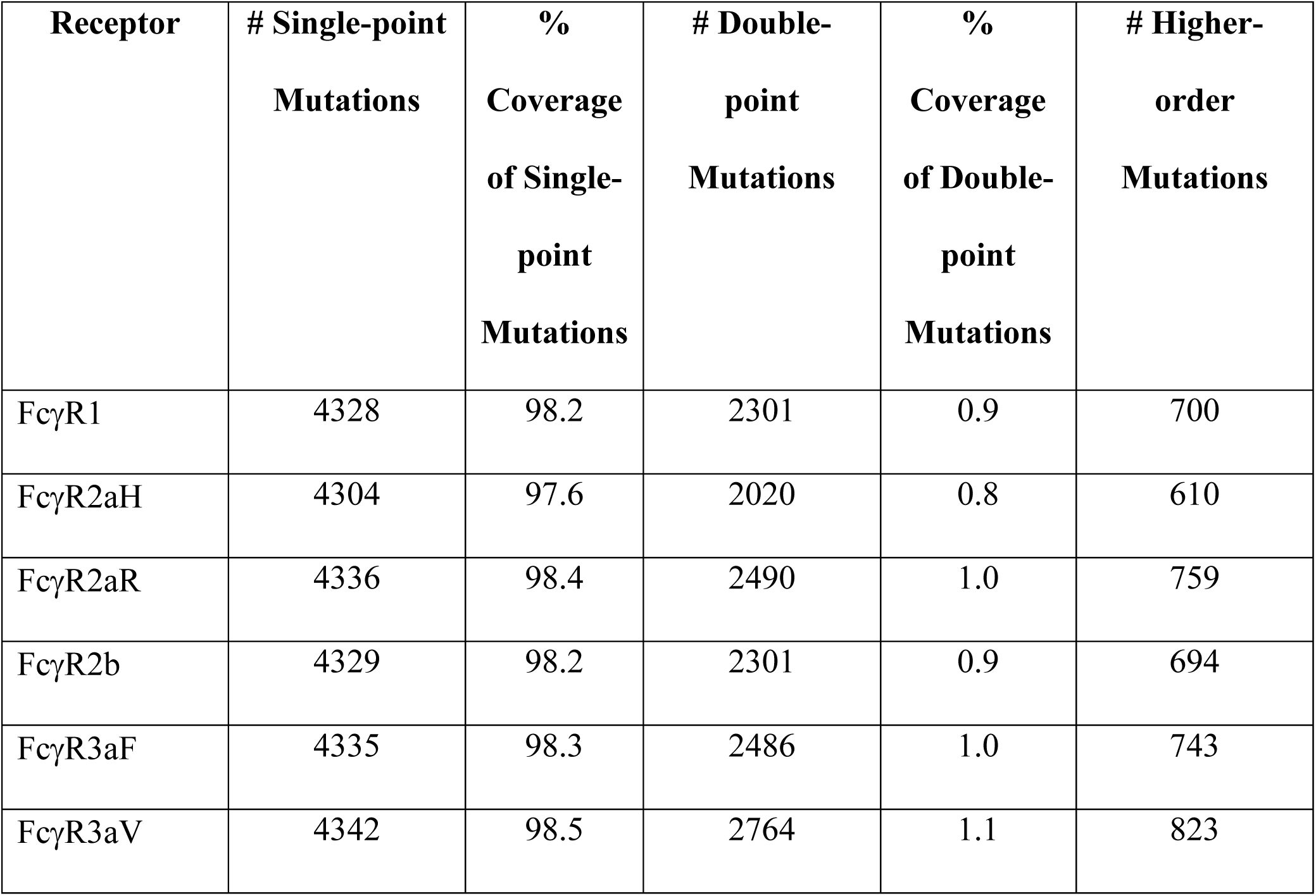
DMS coverage for all Fc-FcγR titrations. The theoretical maximum number of single-point mutations for Fc (IgG1216-447) is 4408, and for double-point mutations is 254,562. Coverage for single-point mutations is consistently close to 100%, as expected from the experimental design, which prioritizes these mutations. Coverage for double-point mutations consistently approximates to 1%. Higher-order mutations denote triple-point mutations and above.

**Figure S9.**
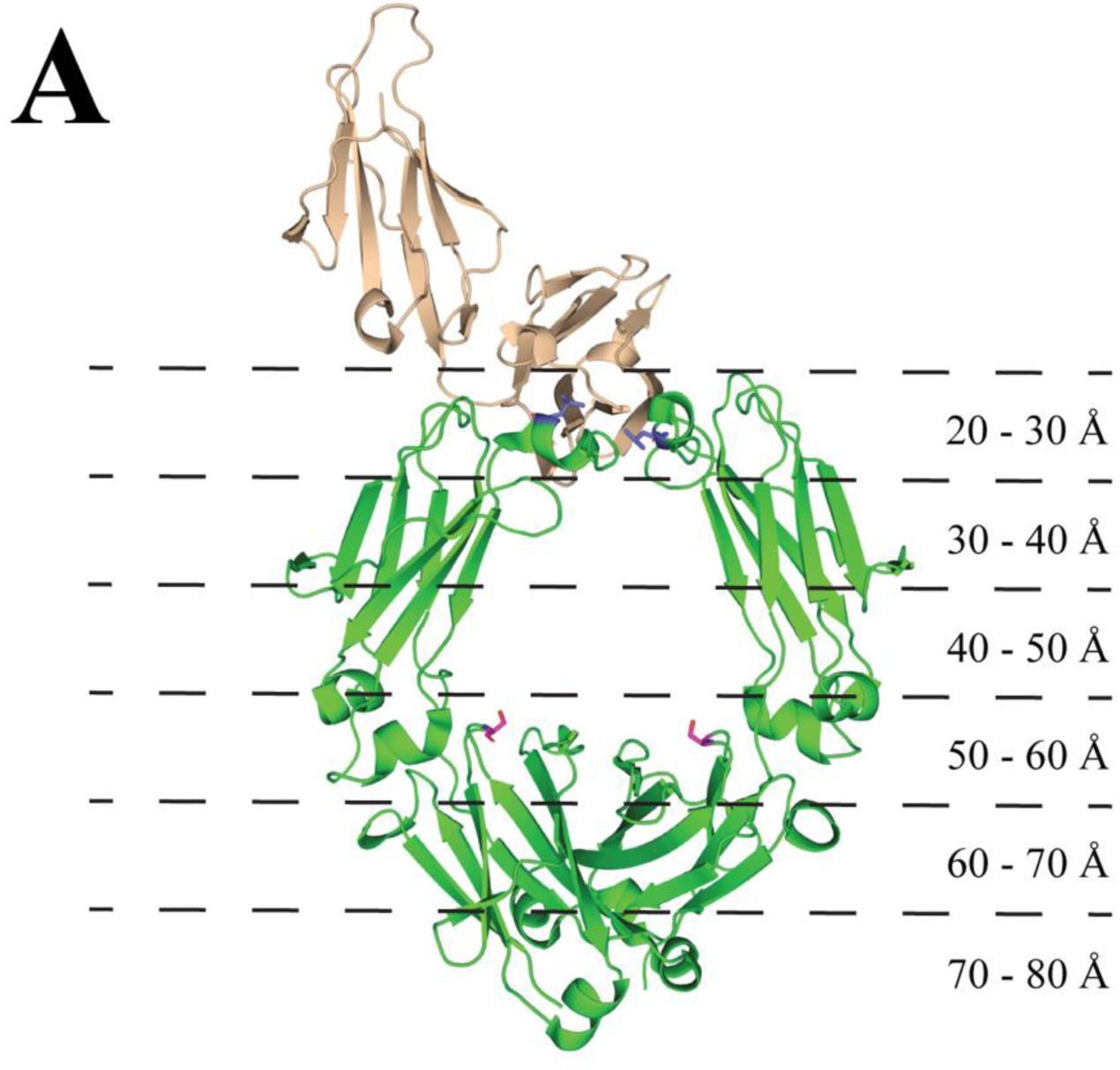
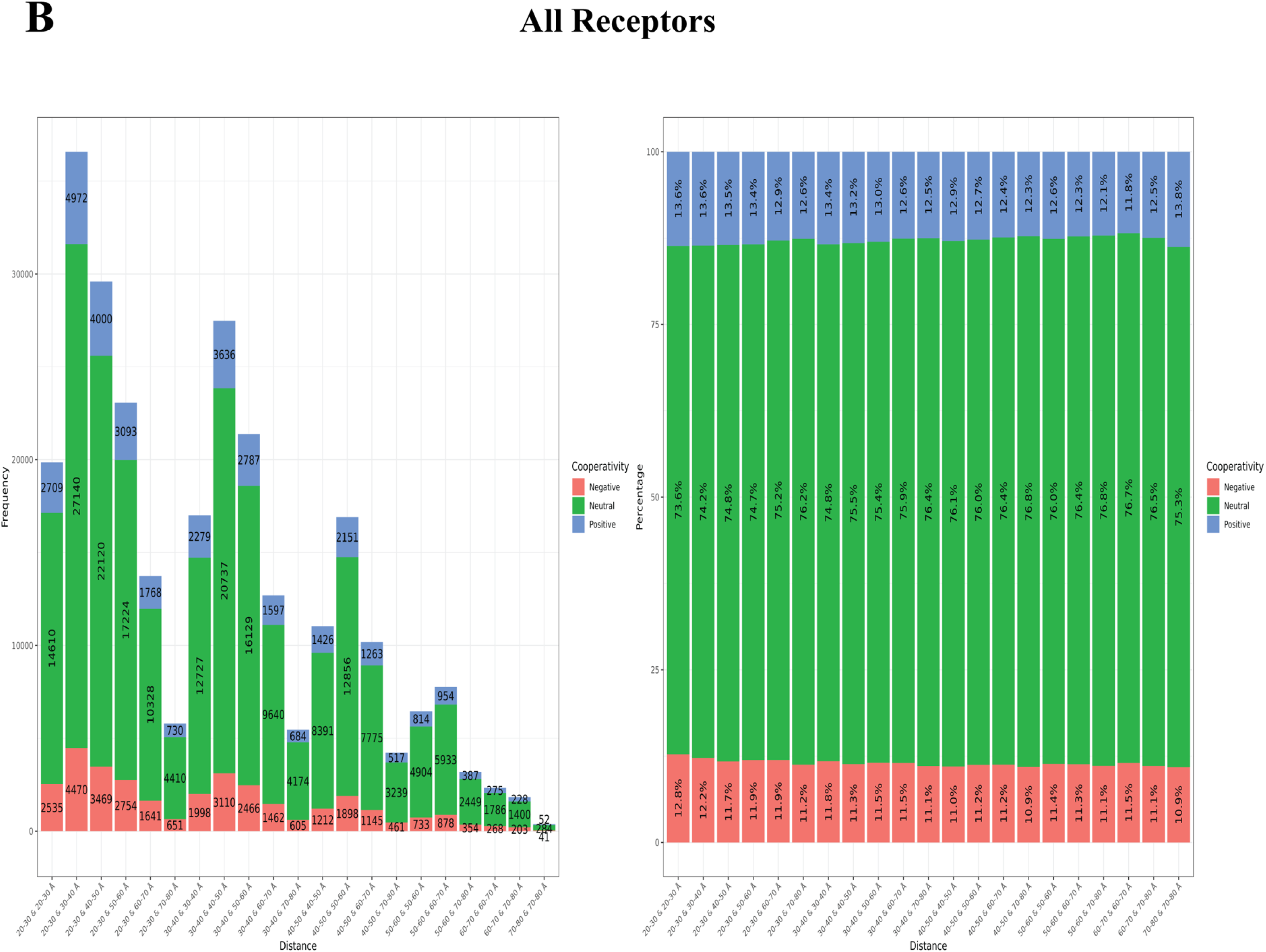
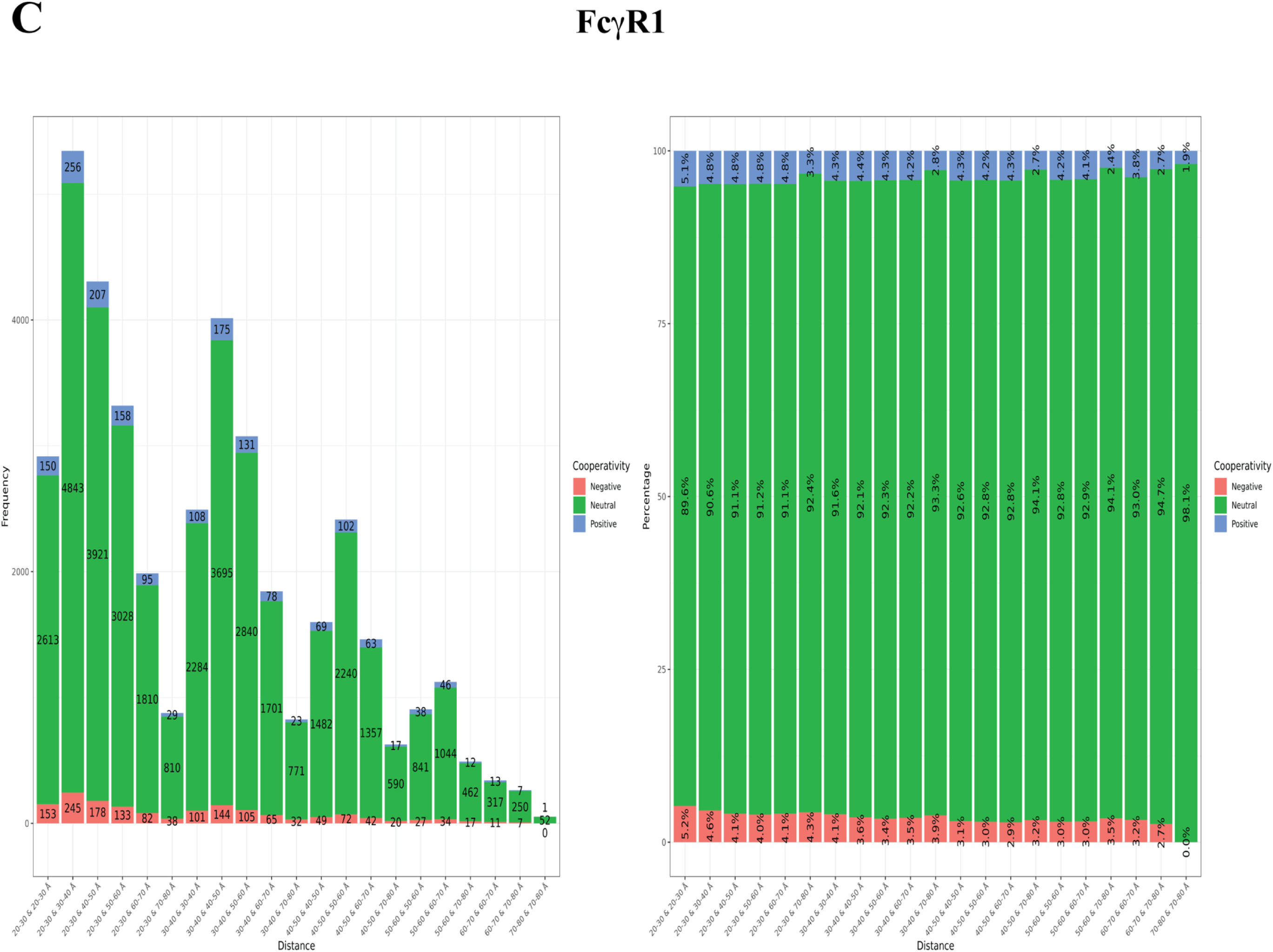
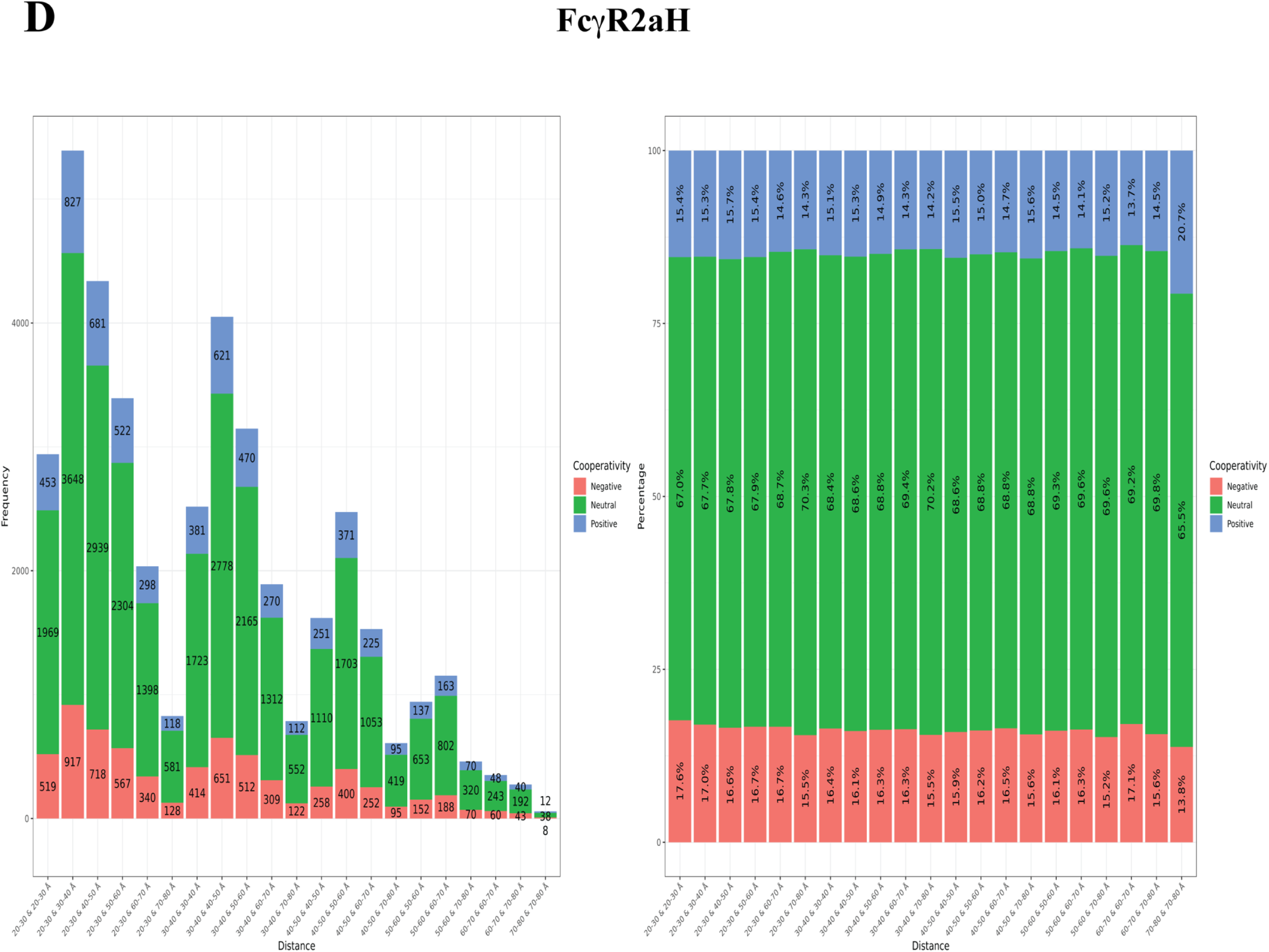
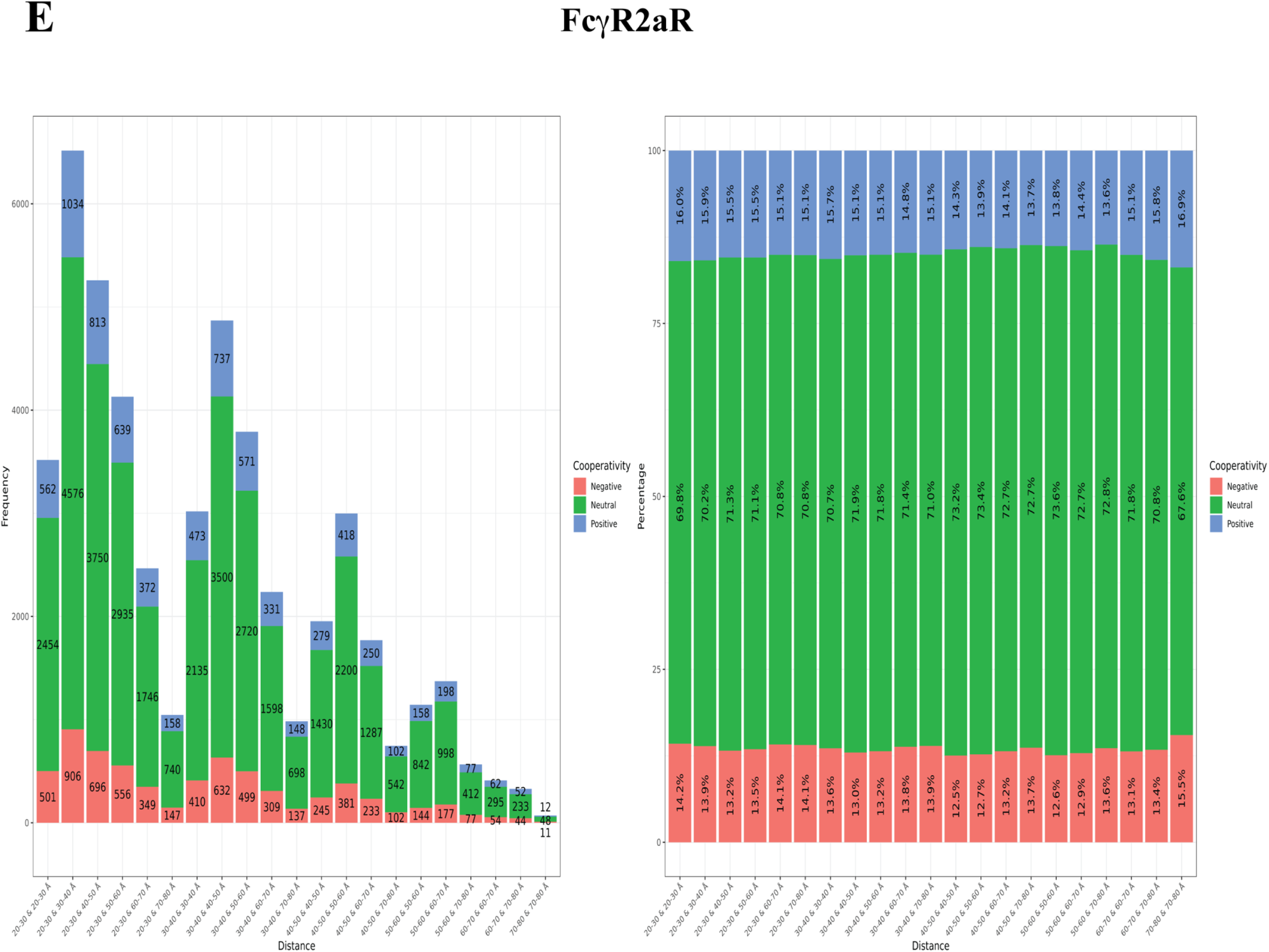
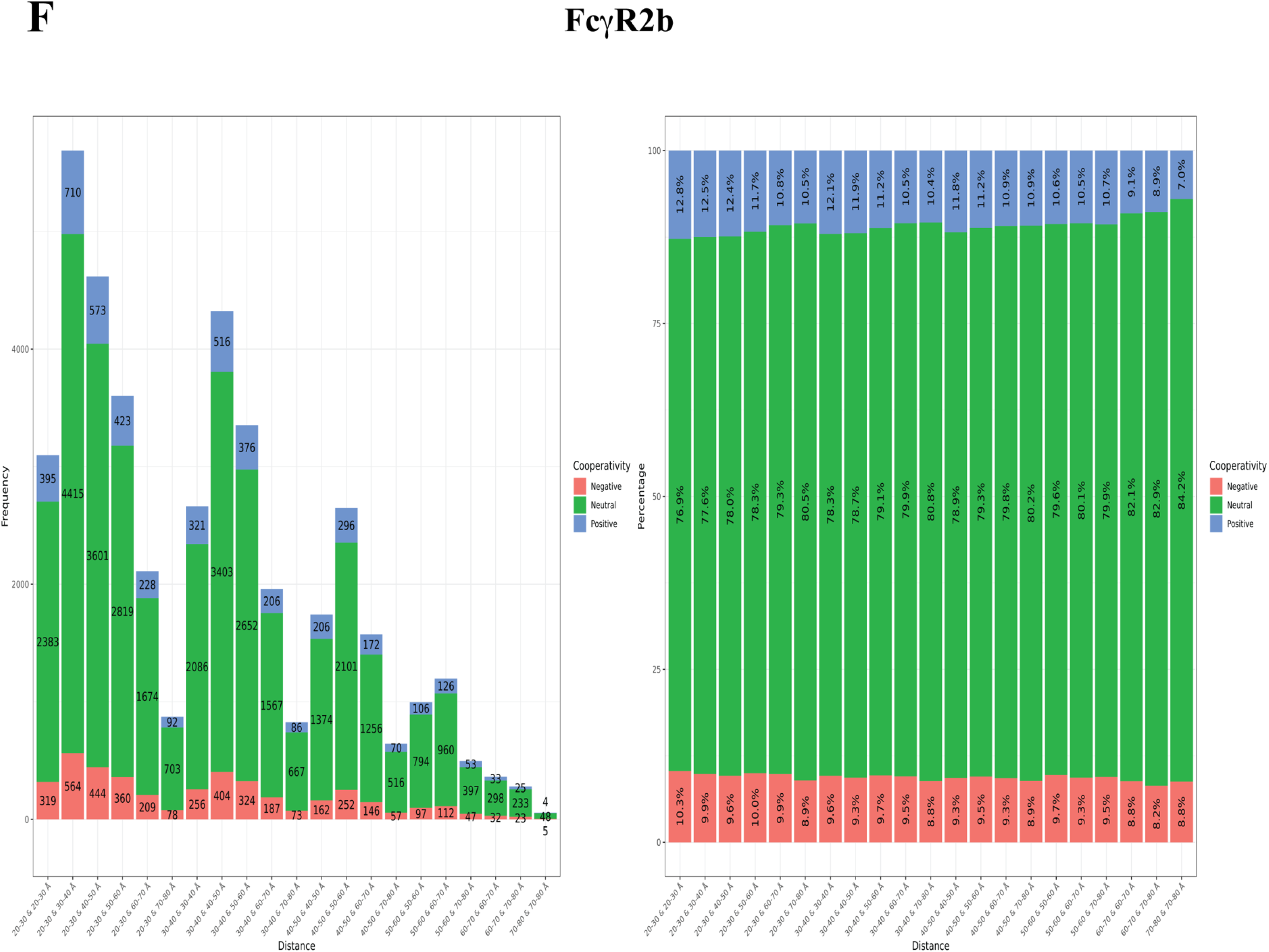
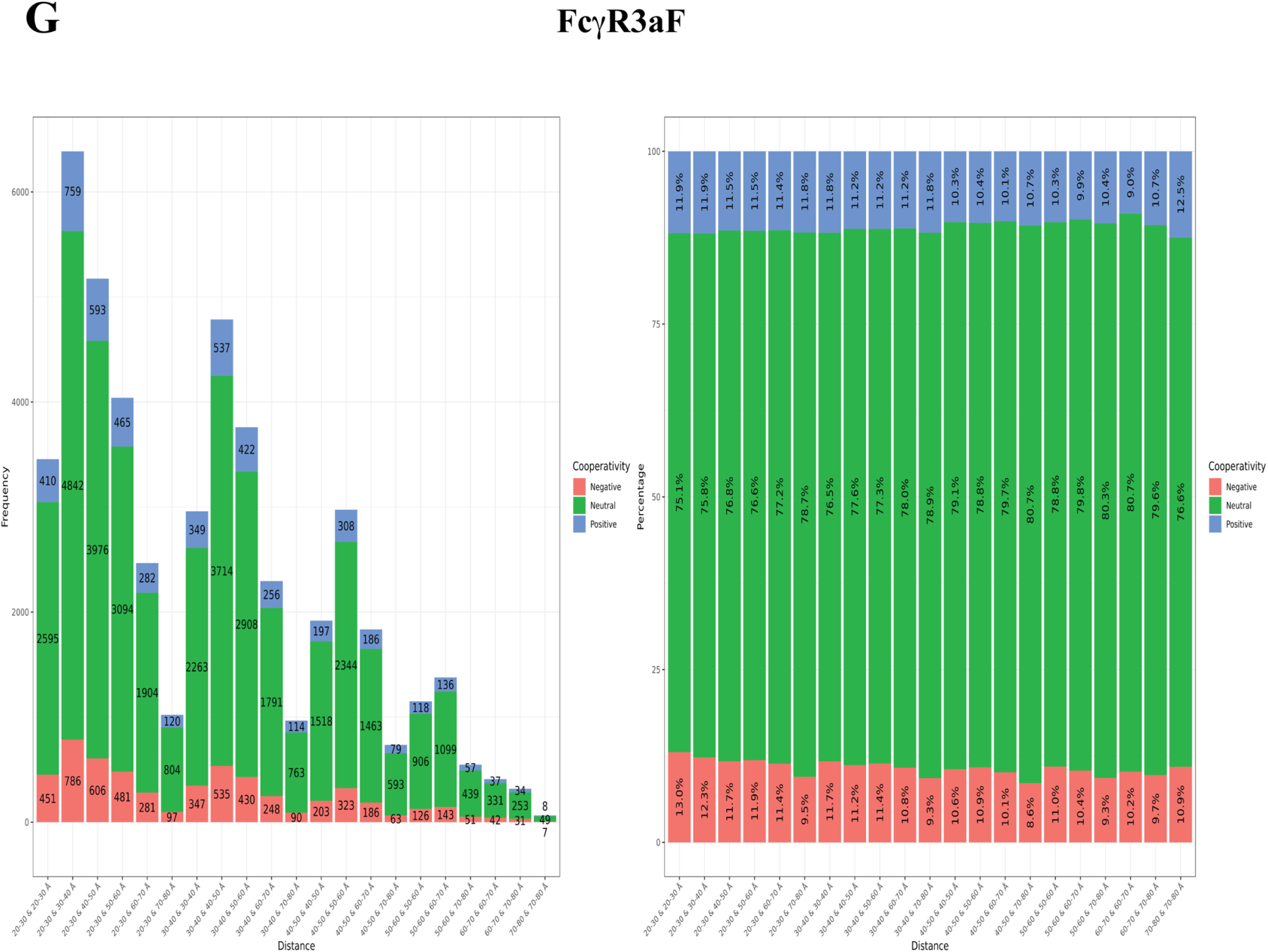
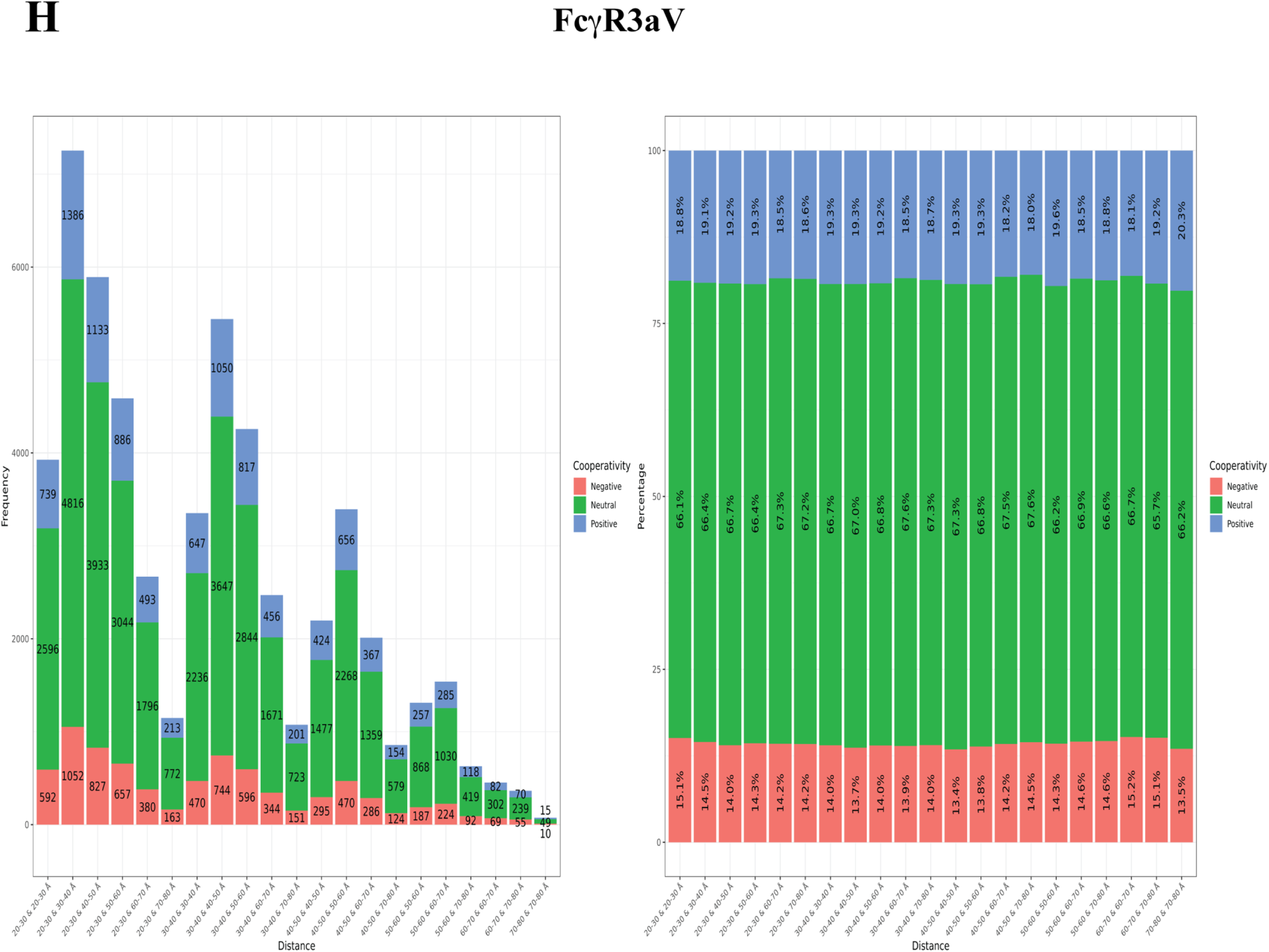
Cooperativity vs mutant-receptor and distances. For each double mutant, the distances of each mutating residue from the receptor are calculated, and the results binned. Bars are further split into whether the double mutation is positively cooperative (blue), additive (green) or negatively cooperative (red), where additivity is defined as being -0.25 < ΔlogKA,*ij* - (ΔlogKA,*i* + ΔlogKA,*j*) < 0.25 with *i* being one mutation and *j* being the other mutation constituting the overall double mutant. For (**B-H**), the *(left)* plot gives the frequencies for each bin and the (*right)* plot gives normalized data so that the percentage of double mutants with positive/neutral/negative cooperativity is displayed. (**A**) Schematic showing the binning scheme; (**B**) Tally of all receptors studied; (**C**) FcγR1; (**D**) FcγR2a-131H; (**E**) FcγR2a-131R; (**F**) FcγR2b; (**G**) FcγR3a-158F; (**H**) FcγR3a-158V.

## References

1. Spencer, D.A., Goldberg, B.S., Pandey, S., Ordonez, T., Dufloo, J., Barnette, P., Sutton, W.F., Henderson, H., Agnor, R., Gao, L., et al. (2022). Phagocytosis by an HIV antibody is associated with reduced viremia irrespective of enhanced complement lysis. Nat. Commun. 13, 662.

2. Zhang, A., Stacey, H.D., D’Agostino, M.R., Tugg, Y., Marzok, A., and Miller, M.S. (2023). Beyond neutralization: Fc-dependent antibody effector functions in SARS-CoV-2 infection. Nat. Rev. Immunol. 23, 381–396.

3. Cao, X., Chen, J., Li, B., Dang, J., Zhang, W., Zhong, X., Wang, C., Raoof, M., Sun, Z., Yu, J., et al. (2022). Promoting antibody-dependent cellular phagocytosis for effective macrophage-based cancer immunotherapy. Sci. Adv. 8, eabl9171.

4. Li, H., Somiya, M., and Kuroda, S. (2021). Enhancing antibody-dependent cellular phagocytosis by re-education of tumor-associated macrophages with resiquimod-encapsulated liposomes. Biomaterials 268, 120601.

5. Su, K., Wu, J., Edberg, J.C., Li, X., Ferguson, P., Cooper, G.S., Langefeld, C.D., and Kimberly, R.P. (2004). A promoter haplotype of the immunoreceptor tyrosine-based inhibitory motif-bearing FcγRIIb alters receptor expression and associates with autoimmunity. I. Regulatory *FCGR2B* polymorphisms and their association with systemic lupus erythematosus. J. Immunol. 172, 7186–7191.

6. Su, K., Wu, J., Edberg, J.C., Li, X., Ferguson, P., Cooper, G.S., Langefeld, C.D., and Kimberly, R.P. (2004). A Promoter Haplotype of the Immunoreceptor Tyrosine-Based Inhibitory Motif-Bearing FcγRIIb Alters Receptor Expression and Associates with Autoimmunity. II. Differential Binding of GATA4 and Yin-Yang1 Transcription Factors and Correlated Receptor Expression and Function. The Journal of Immunology 172, 7192–7199. 10.4049/jimmunol.172.11.7186.

7. Nimmerjahn, F., and Ravetch, J.V. (2008). Fcγ receptors as regulators of immune responses. Nat. Rev. Immunol. 8, 34–47.

8. Nagelkerke, S.Q., Schmidt, D.E., de Haas, M., and Kuijpers, T.W. (2019). Genetic variation in low-to-medium-affinity Fcγ receptors: functional consequences, disease associations, and opportunities for personalized medicine. Front. Immunol. 10, 2237.

9. Indik, Z.K., Park, J.-G., Hunter, S., and Schreiber, A.D. (1995). The molecular dissection of Fcy receptor mediated phagocytosis. Blood 86, 4389–4399.

10. Wang, X., Mathieu, M., and Brezski, R.J. (2018). IgG Fc engineering to modulate antibody effector functions. Protein Cell 9, 63–73. 10.1007/s13238-017-0473-8.

11. Gogesch, P., Dudek, S., van Zandbergen, G., Waibler, Z., and Anzaghe, M. (2021). The Role of Fc receptors on the effectiveness of therapeutic monoclonal antibodies. Int. J. Mol. Sci. 22, 8947.

12. Mandelboim, O., Malik, P., Davis, D.M., Jo, C.H., Boyson, J.E., and Strominger, J.L. (1999). Human CD16 as a lysis receptor mediating direct natural killer cell cytotoxicity. Proc. Natl. Acad. Sci. USA 96, 5640–5644.

13. van der Heijden, J., Breunis, W.B., Geissler, J., de Boer, M., van den Berg, T.K., and Kuijpers, T.W. (2012). Phenotypic variation in IgG receptors by nonclassical *FCGR2C* alleles. J. Immunol. 188, 1318–1324.

14. Su, K., Yang, H., Li, X., Li, X., Gibson, A.W., Cafardi, J.M., Zhou, T., Edberg, J.C., and Kimberly, R.P. (2007). Expression profile of FcγRIIb on leukocytes and its dysregulation in systemic lupus erythematosus. J. Immunol. 178, 3272–3280.

15. Tsang-A-Sjoe, M.W.P., Nagelkerke, S.Q., Bultink, I.E., Geissler, J., Tanck, M.W., Tacke, C.E., Ellis, J.A., Zenz, W., Bijl, M., Berden, J.H., et al. (2016). Fc-gamma receptor polymorphisms differentially influence susceptibility to systemic lupus erythematosus and lupus nephritis. Rheumatology 55, 939–948.

16. Nowicka, M., Hilton, L.K., Ashton-Key, M., Hargreaves, C.E., Lee, C., Foxall, R., Carter, M.J., Beers, S.A., Potter, K.N., Bolen, C.R., et al. (2021). Prognostic significance of *FCGR2B* expression for the response of DLBCL patients to rituximab or obinutuzumab treatment. Blood Adv. 5, 2945–2957.

17. Roghanian, A., Cragg, M.S., and Frendeus, B. (2016). Resistance is futile: targeting the inhibitory FcγRIIB (CD32B) to maximize immunotherapy. Oncoimmunology 5, e1069939.

18. Aguilar, O.A., Gonzalez-Hinojosa, M.D.R., Arakawa-Hoyt, J.S., Millan, A.J., Gotthardt, D., Nabekura, T., and Lanier, L.L. (2023). The CD16 and CD32b Fc-gamma receptors regulate antibody-mediated responses in mouse natural killer cells. J. Leukoc. Biol. 113, 27–40.

19. Sakai, H., Tanaka, Y., Tazawa, H., Shimizu, S., Verma, S., Ohira, M., Tahara, H., Ide, K., Ishiyama, K., Kobayashi, T., et al. (2017). Effect of Fc-gamma receptor polymorphism on rituximab-mediated B cell depletion in ABO-incompatible adult living donor liver transplantation. Transplant. Direct 3, e164.

20. Gavin, P.G., Song, N., Kim, S.R., Lipchik, C., Johnson, N.L., Bandos, H., Finnigan, M., Rastogi, P., Fehrenbacher, L., Mamounas, E.P., et al. (2017). Association of polymorphisms in *FCGR2A* and *FCGR3A* with degree of trastuzumab benefit in the adjuvant treatment of ERBB2/HER2-positive breast cancer: analysis of the NSABP B-31 trial. JAMA Oncol. 3, 335–341.

21. Wu, J., Edberg, J.C., Redecha, P.B., Bansal, V., Guyre, P.M., Coleman, K., Salmon, J.E., and Kimberly, R.P. (1997). A novel polymorphism of FcgammaRIIIa (CD16) alters receptor function and predisposes to autoimmune disease. J. Clin. Invest. 100, 1059–1070.

22. Koene, H.R., Kleijer, M., Algra, J., Roos, D., E.G.Kr. von dem Borne, A., and de Haas, M. (1997). FcγRIIIa-158V/F polymorphism influences the binding of IgG by natural killer cell FcγRIIIa, independently of the FcγRIIIa-48L/R/H phenotype. Blood 90, 1109–1114.

23. Mellor, J.D., Brown, M.P., Irving, H.R., Zalcberg, J.R., and Dobrovic, A. (2013). A critical review of the role of Fc gamma receptor polymorphisms in the response to monoclonal antibodies in cancer. J. Hem. Oncol. 6, 1.

24. Gauthier, L., and Vivier, E. (2020). Boosting cytotoxic antibodies against cancer. Cell 180, 822–824.

25. Zahavi, D., AlDeghaither, D., O’Connell, A., and Weiner, L.M. (2018). Enhancing antibody-dependent cell-mediated cytotoxicity: a strategy for improving antibody-based immunotherapy. Antib. Ther. 1, 7–12.

26. Mandó, P., Rivero, S.G., Rizzo, M.M., Pinkasz, M., and Levy, E.M. (2021). Targeting ADCC: a different approach to HER2 breast cancer in the immunotherapy era. Breast 60, 15–25.

27. Herbrand, U. (2016). Antibody-dependent cellular phagocytosis: the mechanism of action that gets no respect: a discussion about improving bioassay reproducibility. BioProcessing J. 15, 26–29.

28. Su, K., Li, X., Edberg, J.C., Wu, J., Ferguson, P., and Kimberly, R.P. (2004). A promoter haplotype of the immunoreceptor tyrosine-based inhibitory motif-bearing FcγRIIb alters receptor expression and associates with autoimmunity. II. Differential binding of GATA4 and Yin-Yang1 transcription factors and correlated receptor expression and function. J. Immunol. 172, 7192–7199.

29. Rieth, N., Carle, A., Muller, M.A., ter Meer, D., Direnberger, C., Pohl, T., and Sondermann, P. (2014). Characterization of SM201, an anti-hFcγRIIB antibody not interfering with ligand binding that mediates immune complex dependent inhibition of B cells. Immunol. Lett. 160, 145–150.

30. Iwata, Y., Katada, H., Okuda, M., Doi, Y., Ching, T.J., Harada, A., Takeiri, A., Honda, M., and Mishima, M. (2023). Preclinical *in vitro* evaluation of immune suppression induced by GYM329, Fc-engineered sweeping antibody. J. Toxicol. Sci. 48, 399–409.

31. Li, X., and Kimberly, R.P. (2014). Targeting the Fc receptor in autoimmune disease. Expert Opin. Ther. Targets 18, 335–350.

32. Yamaguchi, Y., and Barb, A.W. (2020). A synopsis of recent developments defining how N-glycosylation impacts immunoglobulin G structure and function. Glycobiology 30, 214–225. 10.1093/glycob/cwz068.

33. Frank, F., Keen, M.M., Rao, A., Bassit, L., Liu, X., Bowers, H.B., Patel, A.B., Cato, M.L., Sullivan, J.A., Greenleaf, M., et al. (2022). Deep mutational scanning identifies SARS-CoV-2 Nucleocapsid escape mutations of currently available rapid antigen tests. Cell 185, 3603–3616.e3613.

34. Starr, T.N., Greaney, A.J., Hilton, S.K., Ellis, D., Crawford, K.H.D., Dingens, A.S., Navarro, M.J., Bowen, J.E., Tortorici, M.A., Walls, A.C., et al. (2020). Deep mutational scanning of SARS-CoV-2 receptor binding domain reveals constraints on folding and ACE2 binding. Cell 182, 1295–1310.e1220.

35. Bruhns, P., Iannascoli, B., England, P., Mancardi, D.A., Fernandez, N., Jorieux, S., and Daeron, M. (2009). Specificity and affinity of human Fcγ receptors and their polymorphic variants for human IgG subclasses. Blood 113, 3716–3725.

36. Chen, W., Kong, L., Connelly, S., Dendle, J.M., Liu, Y., Wilson, I.A., Powers, E.T., and Kelly, J.W. (2016). Stabilizing the CH2 Domain of an Antibody by Engineering in an Enhanced Aromatic Sequon. ACS Chem Biol 11, 1852–1861. 10.1021/acschembio.5b01035.

37. Feige, M.J., Walter, S., and Buchner, J. (2004). Folding mechanism of the CH2 antibody domain. J Mol Biol 344, 107–118. 10.1016/j.jmb.2004.09.033.

38. Subedi, G.P., and Barb, A.W. (2015). The structural role of antibody N-glycosylation in receptor interactions. Structure 23, 1573–1583. 10.1016/j.str.2015.06.015.

39. Ferrara, C., Grau, S., Jäger, C., Sondermann, P., Brünker, P., Waldhauer, I., Hennig, M., Ruf, A., Rufer, A.C., Stihle, M., et al. (2011). Unique carbohydrate–carbohydrate interactions are required for high affinity binding between FcγRIII and antibodies lacking core fucose. Proc Natl Acad Sci USA 108, 12669–12674.

40. Richards, J.O., Karki, S., Lazar, G.A., Chen, H., Dang, W., and Desjarlais, J.R. (2008). Optimization of antibody binding to FcγRIIa enhances macrophage phagocytosis of tumor cells. Mol. Cancer Ther. 7, 2517–2527.

41. Chu, S.Y., Vostiar, I., Karki, S., Moore, G.L., Lazar, G.A., Pong, E., Joyce, P.F., Szymkowski, D.E., and Desjarlais, J.R. (2008). Inhibition of B cell receptor-mediated activation of primary human B cells by coengagement of CD19 and FcγRIIb with Fc-engineered antibodies. Mol. Immunol. 45, 3926–3933.

42. Mimoto, F., Katada, H., Kadono, S., Igawa, T., Kuramochi, T., Muraoka, M., Wada, Y., Haraya, K., Miyazaki, T., and Hattori, K. (2013). Engineered antibody Fc variant with selectively enhanced FcγRIIb binding over both FcγRIIa(R131) and FcγRIIa(H131). Protein Eng. Des. Sel. 26, 589–598.

43. Liu, R., Oldham, R.J., Teal, E., Beers, S.A., and Cragg, M.S. (2020). Fc-engineering for modulated effector functions - improving antibodies for cancer treatment. Antibodies 9, 64.

44. Kiyoshi, M., Caaveiro, J.M.M., Kawai, T., Tashiro, S., Ide, T., Asaoka, Y., Hatayama, K., and Tsumoto, K. (2015). Structural basis for binding of human IgG1 to its high-affinity human receptor FcγRI. Nat Commun 6.

45. Bournazos, S., Vo, H.T.M., Duong, V., Auerswald, H., Ly, S., Sakuntabhai, A., Dussart, P., Cantaert, T., and Ravetch, J.V. (2021). Antibody fucosylation predicts disease severity in secondary dengue infection. Science 372, 1102–1105. 10.1126/science.abc7303.

46. Shields, R.L., Namenuk, A.K., Hong, K., Meng, Y.G., Rae, J., Briggs, J., Xie, D., Lai, J., Stadlen, A., Li, B., et al. (2001). High resolution mapping of the binding site on human IgG1 for FcγRI, FcγRII, FcγRIII, and FcRn and design of IgG1 variants with improved binding to the FcγR. J. Biol. Chem. 276, 6591–6604.

47. Sondermann, P., Kaiser, J., and Jacob, U. (2001). Molecular basis for immune complex recognition: a comparison of Fc-receptor structures. J. Mol. Biol. 309, 737–749.

48. Ahmed, A.A., Keremane, S.R., Vielmetter, J., and Bjorkman, P.J. (2016). Structural characterization of GASDALIE Fc bound to the activating Fc receptor FcγRIIIa. J. Struct. Biol. 194, 78–89.

49. Weitzenfeld, P., Bournazos, S., and Ravetch, J.V. (2019). Antibodies targeting sialyl Lewis A mediate tumor clearance through distinct effector pathways. J. Clin. Invest. 129, 3952–3962.

50. Greaney, A.J., Starr, T.N., Gilchuk, P., Zost, S.J., Binshtein, E., Loes, A.N., Hilton, S.K., Huddleston, J., Eguia, R., Crawford, K.H.D., et al. (2021). Complete mapping of mutations to the SARS-CoV-2 Spike receptor-binding domain that escape antibody recognition. Cell Host Microbe 29, 44–57.e49.

51. Crawford, K.H.D., and Bloom, J.D. (2019). alignparse: A Python package for parsing complex features from high-throughput long-read sequencing. J. Open Source Softw. 4, 1915.

52. Li, H. (2018). Minimap2: pairwise alignment for nucleotide sequences. Bioinformatics 34, 3094–3100.

